# New insights into bryophyte arabinogalactan-proteins from a hornwort and a moss model organism

**DOI:** 10.1101/2025.03.14.643249

**Authors:** Kim-Kristine Mueller, Lukas Pfeifer, Linus Wegner, Katrin Ehlers, Birgit Classen

## Abstract

Two bryophyte models, the hornwort *Anthoceros agrestis* (*Anthoceros*) and the moss *Physcomitrium patens* (*Physcomitrium*), were analysed for presence of arabinogalactan-proteins (AGPs), as emergence of these signalling glycoproteins in evolution is still under debate. AGPs of both species had a galactan core structure similar to that of other bryophyte and fern AGPs, but different to angiosperm AGPs, as 1,6-linked pyranosidic galactose was almost absent. In the *Physcomitrium* AGP, furanosidic arabinose (Ara*f*) linkages were mainly terminal (10 %) or 5-linked (13 %), while in *Anthoceros*, terminal Ara*f* dominated (26 %) and was accompanied by very low amounts of 1,3-Ara*f* and pyranosidic terminal Ara. Unusual 3-*O*-methylated pyranosidic rhamnose, which has never been detected in cell walls of angiosperms, occurred in both bryophyte AGPs (5 % in *Anthoceros*, 10 % in *Physcomitrium* AGP), This was comparable to AGPs of other spore-producing land plants. Bioinformatic search in genomes of 14 bryophyte species revealed that most hornworts lack sequences encoding GPI-anchored classical AGPs. Generally, hornworts contained less sequences for AGP protein backbones compared to the liverwort *Marchantia polymorpha* and the moss *Physcomitrium patens*. All of them comprise sequences for chimeric AGPs, and among those surprisingly xylogen-like AGPs. Homologous sequences encoding glycosyltransferases and other enzymes involved in the synthesis and decoration of the AGP galactan framework were present in all bryophyte genomes. Immunocytochemistry of *Anthoceros* tissue detected AGPs at the plasma membrane/cell wall interface but also at vacuolar and vesicle membranes, suggesting new functions of AGPs in bryophytes.

**SIGNIFICANCE STATEMENT:** Extant bryophytes are key to infer evolution of the most recent common ancestor of all land plants. As cell walls were important for adaptation to life on land, we analysed arabinogalactan-proteins from the hornwort *Anthoceros agrestis* and the moss *Physcomitrium patens* and detected terminal 3-*O*-methylrhamnose residues, which also occur in fern AGPs but not in angiosperms. Bioinformatic search for AGP protein backbones and glycosyltransferases in bryophyte genomes further strengthens understanding of AGP evolution during terrestrialization.

## INTRODUCTION

The split of the two major land plant clades, bryophytes and tracheophytes, is dated more than 470 million years back (Yadav *et al*., 2023). Since then, the bryophytes with their remarkable biodiversity, formed by more than 20,000 extant species, adapted to almost every habitat (Morris *et al*., 2018; Degola *et al*., 2022, Wang *et al*., 2022). Recent phylogenomic evidence considers bryophytes as monophyletic (Zhang *et al*., 2020; Wang *et al*., 2022; Bechteler *et al*., 2023) with hornworts as sister clade to the so-called setaphytes. The latter group consists of both bryophyte groups carrying a stalked spore capsule, namely liverworts and mosses. As hornworts are the deepest-diverging phylum of all extant land plants, they are important to reconstruct the evolution of land plants (Schaffran *et al*., 2025). In order to investigate the molecular basis of various biological processes, many researchers supported the establishment of model organisms for each bryophyte clade. Milestones for this ambitious aim were the successive sequencing of genomes from all of the three bryophyte groups (Rensing *et al*., 2008, Bowman *et al*., 2017; Li *et al*., 2020; Zhang *et al*., 2020, Schafran *et al*., 2025).

Bryophytes show an extreme tolerance against abiotic stressors, like e.g. changing light status, temperature and water availability (de Vries & Archibalt, 2018; Fürst-Jansen *et al*., 2020; Bowles *et al*., 2021; Yadav *et al*., 2023). This is reflected by the enormous diversity of the natural habitats, ranging from west tropical forests to the Arctic (Hodgetts *et al*., 2019; Wang *et al*., 2022). The extreme stress tolerance can also be seen as evolutionary heritage of the most recent common ancestor (MRCA) of all embryophyta. That MRCA had to cope with numerous environmental changes going hand-in-hand with plant terrestrialization and this flexibility formed the basis for the flourishing of all land plant taxa (de Vries & Archibalt, 2018, Domozych & Bagdan, 2022, McCourt *et al*., 2023).

One of the many important adaptation steps was the modification of the cell wall (Harholt *et al*., 2016). This highly complex and dynamic system protects the cell from various stressors, both abiotic and biotic. Furthermore, it mechanically stabilizes the plant. The main components of the cell wall are polysaccharides - mainly cellulose, hemicelluloses and pectic polysaccharides - as well as cell wall (glyco)proteins. All of these macromolecules were described to be generally present in bryophytes (Geddes & Wilkie, 1971; Kremer *et al*., 2004; Carafa *et al*., 2005; Berry *et al*., 2016; Pfeifer *et al*., 2022a). Although a general similarity to tracheophyte walls is visible (for reviews, see Ye & Zhong, 2022; Pfeifer *et al*., 2022a), in-detail studies gave rise to some exciting new discoveries. Some examples are altered basic motifs in xyloglucans of mosses and liverworts (Peña *et al*., 2008) or arabinoglucans from *Physcomitrium patens*, which show structural features of mixed-linked glucans (Roberts *et al*., 2018).

Cell wall (glyco)proteins are key players in plant cell wall biochemistry. The most dominant types are hydroxyproline-rich glycoproteins (HRGPs). These can be further divided into arabinogalactan-proteins (AGPs), extensins (EXT) and proline-rich proteins (PRPs) with differing degrees of glycosylation. While AGPs show huge glycan parts of around 90 % of the molecule, only 50 % of EXTs and even lower portions of PRPs are built up of carbohydrates (Johnson *et al*., 2018). Both moieties (protein and carbohydrates) are covalently connected *via* the 4-*O*-hydroxylated derivative of proline (= hydroxyproline, Hyp). Glycosylation in the polysaccharide part shows the characteristics of type II arabinogalactans, formed by (1→3)- linked pyranosidic β-D-galactose (Gal*p*) units which are connected to (1→6)-β-D-galactan side chains at position O-6. This galactan backbone is further decorated with furanosidic α-L-arabinose (Ara*f*) and other monosaccharides in minor amounts, such as L-rhamnose (Rha), L-fucose (Fuc) or D-glucuronic acid (GlcA; Kitazawa *et al*., 2013; Strasser *et al*., 2021). The distribution of AGPs among land plants is widespread and also bryophytes were investigated very early in the 1970s by Clarke *et al*. (1978). The authors used the interaction between the AGP glycan part and β-glucosyl Yariv reagent for precipitation (βGlcY; Yariv *et al*., 1962). AGPs from all major land plant lineages, i.e. ferns, gymnosperms, as well as monocots and dicots (Classen *et al*., 2004; Goellner *et al*., 2011, Bartels & Classen, 2017; Baumann *et al*., 2021, Mueller *et al*., 2023) were analyzed over the last decades. Furthermore, few studies focused on bryophytes, but characterized only AGPs of mosses and liverworts (Lee *et al*., 2005; Fu *et al*., 2007; Bartels *et al*., 2017; Happ & Classen, 2019) and not of hornworts. Beside a general similarity of spore plant AGPs to angiosperm AGPs, significant structural peculiarities, namely high amounts of methylated rhamnose (3-*O*-MeRha; acofriose; Fu *et al*., 2007; Bartels *et al*., 2017; Bartels & Classen, 2017; Happ & Classen, 2019; Mueller *et al*., 2023), were found.

Seed plant AGPs are involved in many developmental processes, including xylem differentiation, cell proliferation, growth, salt and drought tolerance, as well as interaction with plant microbes (Seifert & Roberts, 2007; Ma *et al*., 2018; Mareri *et al*., 2018). Bryophyte AGPs were described to contribute to water balance in mosses (Ligrone *et al*., 2002; Kobayashi *et al*., 2011; Cui *et al*., 2012) as well as to apical cell extension in protonemata of the moss *P. patens* (Lee *et al*., 2005).

In this study, we hypothesize that AGPs in bryophytes are important for the successful adaptation to life on land. Bryophyte AGPs of the model moss *P. patens* and the model hornwort *A. agrestis* were purified to further understand the nature of AGP glycans in bryophytes. No hornwort AGP has been isolated and characterized up to now. Our investigations are complemented by immunocytochemical detection of AGPs in hornwort tissue. Bioinformatic search for AGP protein backbones and glycosyltransferases in genomes of all bryophyte lineages (Rensing *et al*., 2008, Bowman *et al*., 2017; Schafran *et al*., 2025) further strengthens understanding of AGP evolution during terrestrialization.

## RESULTS

### Gel diffusion assay

A gel diffusion assay was used to search for AGPs in the high molecular weight, water-soluble fractions (AE) of the bryophytes. AGPs contain 1,3-galactan chains that precipitate with red-colored Yariv phenyl glycosides, such as β-D-glucosyl Yariv-reagent (βGlcY; Kitazawa *et al*., 2013; Paulsen *et al*., 2014). This precipitation can be perceived as a red line in a gel diffusion assay (Clarke *et al*., 1979) and was positive for *Anthoceros* and *Physcomitrium* AEs, comparable to AE from *Marchantia polymorpha* (Figure S1), which has been investigated in a previous study (Happ & Classen, 2019).

### Isolation and yield of AGPs

The extraction of the high molecular weight, water-soluble fractions (AE) yielded 12.0 ± 3.1 % for *Anthoceros* plants (n=2), 10.9 % for *Anthoceros* cell suspension culture and 2.1 % of the dry plant material for *Physcomitrium* (n=1). The fractions comprised different monosaccharides, with Glc, Gal, Xyl and Ara dominating in these fractions (Tables S1 and S2). In both AE samples from *Anthoceros* (plant and cell culture), Gal and Ara accounted for over 50 % of the neutral monosaccharides with higher amounts of Ara in the cell culture. In *Physcomitrium*, Glc was dominating, followed by Gal and Ara. It has to be taken into account that at least part of the Glc probably originates from starch due to lack of amylase treatment (Figure S2).

Based on the high amount of Ara and Gal in the AE fractions (Table S1) as well as the positive gel diffusion assays (Figure S1), the AE fractions were treated with βGlcY-reagent to gain AGP fractions. The yields of the *Anthoceros* AGP fractions were very high with 0.31 % ± 0.08 % (plants, n=2) and 0.57 % (cell culture, n=1) of the dry plant material compared to that of *Physcomitrium* (0.09 %, n=1).

### Carbohydrate moiety of bryophyte AGPs

The neutral monosaccharides Ara and Gal were clearly dominant in *Anthoceros* AGP fractions (Table 1), accounting together for 85.7 % (plant) and 90.1 % (cell culture). The content of Ara was remarkably high in AGP from *Anthoceros* cell culture, leading to a ratio of 1 : 0.8 (Ara:Gal) which is unusual for AGPs. In contrast, these ratios in AGPs from *Anthoceros* plants and *Physcomitrium* were quite similar and typical for AGPs (1:1.7 and 1:1.6). High amounts of Glc were additionally present in *Physcomitrium* AGP. Rha was part of all AGPs with a higher amount in *Physcomitrium* and mainly present as 3-*O*-methyl derivative (3-*O*-MeRha).

**Table 1.** Neutral monosaccharide composition of AGPs from *Anthoceros agrestis* plants and cell cultures and *Physcomitrium patens* in % (mol mol^-1^).

| Neutral monosaccharide | *Anthoceros* (plant) |  | *Anthoceros* (cell culture) | *Physcomitrium* |  |
| --- | --- | --- | --- | --- | --- |
|  | AGP (n=1) | AGP<sub>UR</sub> (n=3) | AGP (n=3) | AGP (n=1) | AGP<sub>UR</sub> (n=2) |
| 3-*O*-Me-Rha | 5.6 | 4.3 ± 0.1 | 5.1 ± 0.1 | 10.0 | 10.6 ± 0.4 |
| Rha | tr | tr | tr | 3.4 | 2.8 ± 0.1 |
| Fuc | tr | tr | tr | 2.0 | 1.5 ± 0.1 |
| Ara | 31.8 | 35.6 ± 0.7 | 49.3 ± 0.6 | 21.8 | 21.1 ± 0.0 |
| Xyl | 4.0 | 3.3 ± 0.4 | tr | 2.8 | 2.7 ± 0.3 |
| Man | 1.6 | 1.9 ± 0.3 | 3.4 ± 0.1 | 3.0 | 2.4 ± 0.0 |
| Gal | 53.9 | 50.3 ± 0.8 | 40.8 ± 0.5 | 35.2 | 35.6 ± 0.0 |
| Glc | 3.1 | 4.6 ± 1.0 | 1.4 ± 0.1 | 21.8 | 23.3 ± 0.0 |
| Ara : Gal | 1 : 1.7 | 1 : 1.7 | 1 : 0.8 | 1 : 1.6 | 1 : 1.7 |

tr: trace value < 1 %

The uronic acid content of the AGPs determined by colorimetric quantification was comparable between 5 % and 7 % (Table S2). Furthermore, AGPs were subjected to uronic acid reduction with sodium borodeuteride (AGP_UR_). In the mass spectrum analysis, deuterated fragments were detected only in the peak of Glc (see below, Table 2). This, together with the slight increase of Glc in the neutral monosaccharide analysis after uronic acid reduction (Table 1, AGP_UR_) identified GlcA as part of bryophyte AGPs.

**Table 2.** Linkage type analysis of AGP and AGP_UR_ samples from *Anthoceros agrestis* (plant) and *Physcomitrium patens* in % (mol mol^-1^, n=1).

| Mono-saccharide | linkage type | *Anthoceros* |  | *Physcomitrium* |  |
| --- | --- | --- | --- | --- | --- |
|  |  | AGP | AGP<sub>UR</sub> | AGP | AGP<sub>UR</sub> |
| Galp | 1,3,6- | 28 | 24 | 22 | 18 |
|  | 1,3,4- | 7 | 7 | - | - |
|  | 1,6- | tr | 1 | 1 | 1 |
|  | 1,3- | 20 | 18 | 11 | 11 |
|  | t- | 1 | 1 | 2 | 2 |
| Glcp | 1,4- | - | 2* | 23 | 20 |
|  | t- | 1 | 2* | 2 | 4* |
| Manp | 1,6- | tr | tr | tr | 2 |
| Hexp | 1,3,4,6- | 7 | 7 | - | - |
| Rhap | t- | 8 | 7 | 17 | 18 |
| Fucp | t- | - | - | 1 | 1 |
| Araf | 1,5- | tr | tr | 11 | 13 |
|  | 1,3- | 2 | 2 | tr | tr |
|  | t- | 24 | 26 | 10 | 10 |
| Arap | t- | 2 | 2 | tr | tr |
| Xylp | t- | tr | 1 | tr | tr |

tr: trace value < 1 %; UR: uronic acid-reduced; \* including deuterated fragments

### Structure elucidation of the arabinogalactan (AG) moieties of bryophyte AGPs

For structural characterization, the AGPs and the uronic acid-reduced AGPs (AGP_UR_) were subjected to linkage-type analysis (Table 2). Figure 1 shows the GC-chromatogramms of both samples after *per*-methylation and acetylation together with structural proposals for *Anthoceros* and *Physcomitrium* AGPs. In all samples, Gal*p* was the main monosaccharide and present mainly in 1,3,6-linkage (18 % – 28 %) and 1,3-linkage (11 % – 20 %). In *Anthoceros* AGP, 1,3,4-linked Gal was present additionally. The linkage-types of Ara also differed in the two investigated bryophytes. In *Anthoceros* AGP, terminal Ara*f* was strongly dominant (24 % / 26 %), whereas in *Physcomitrium* AGP, more 1,5-Ara*f* was present and accompanied by almost similar amounts of terminal Ara*f*.

**Figure 1.**
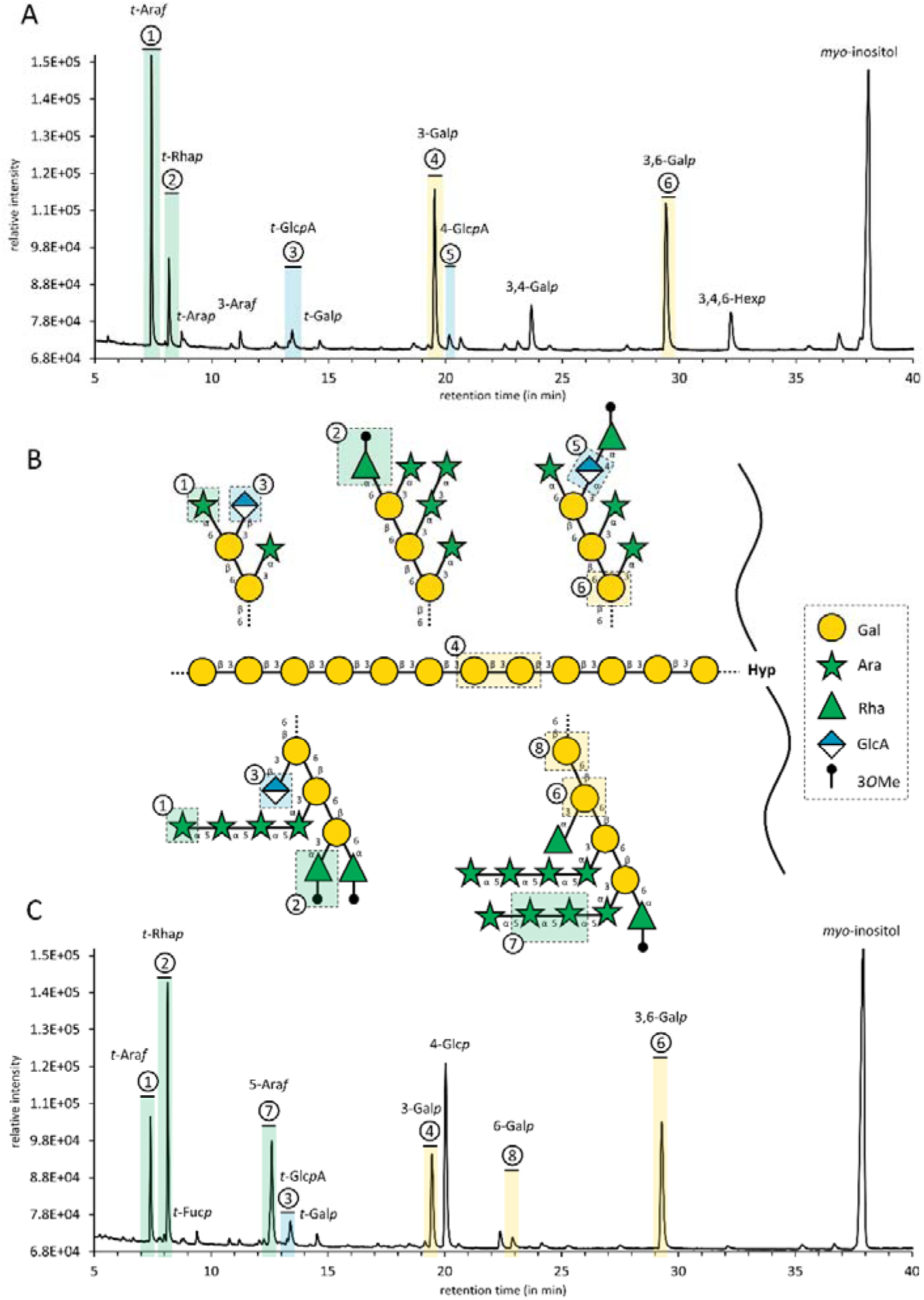
Results of the linkage-type analysis of *Anthoceros agrestis* and *Physcomitrium patens* AGPs. The highlighted structural features are numbered with the same numbers in all three panels of the diagram. **A**: Gas-chromatogram of methylated and *per*-acetylated monosaccharides derived from *Anthoceros* AGP_UR_. **B**: Structural proposal of fragments inferred from linkage analysis. The upper half of this panel represents *Anthoceros* fragments while the lower half represents *Physcomitrium* fragments. The central 1,3-linked Gal*p* backbone is shared. **C:** Gas-chromatogram of methylated and *per*-acetylated monosaccharides derived from *Physcomitrium* AGP_UR_.

Further terminal monosaccharides of both AGPs were t-Rha*p* with up to 18 % in *Physcomitrium*, terminal Gal*p* and terminal Glc*p*A. In the AGP samples, uronic acids are not detectable due to the method used. After reduction of uronic acids with borodeuteride (AGP_UR_), deuterated fragments were detected in the peak of terminal Glc and the amount of this peak slightly increased, indicating presence of terminal GlcA in both native AGPs (Table 2). Further deuterated fragments were found in 1,4-linked Glc present in *Anthoceros* AGP in small amounts, possibly originating from 1,4-GlcA. Additionally, an 1,3,4,6-Hex*p* peak was identified in *Anthoceros* AGP. Since it cannot be clearly determined whether this is an artificial peak due to undermethylation, it was not included in the structural proposal (Figure 1). In the *Physcomitrium* AGP sample, high amounts of 1,4-linked Glc (20 %) were detected. As this might originate from accompanying starch, it was also not included in the structural proposal (Figure 1). Fuc, Man and Xyl were not found in significant amounts in both bryophyte AGPs.

### Characterization of AG epitopes with antibodies

To obtain more information about the structure of the AG moieties of the two bryophyte AGPs, they were tested by ELISA for their binding affinity to different antibodies directed against AGP glycan epitopes (JIM13, KM1, LM2 and LM6; Figure 2; Table S3). The antibody JIM13 (Figure 2A; Yates *et al*., 1996; Pfeifer *et al*., 2022b) is generally considered to bind to AGPs, and the trisaccharide structure β-D-Glc*p*A-(1→3)-α-D-Gal*p*A-(1→2)-α-L-Rha has been suggested as an epitope. The AGPs of *Physcomitrium* showed a strong and that of *Anthoceros* a moderate affinity to JIM13. KM1, directed against (1→6)-β-D-Gal*p* units in AGs type II, (Figure 2B; Classen *et al*., 2004; Ruprecht *et al*., 2017) did not bind to the *Anthoceros* AGPs and had very weak reactivity with the *Physcomitrium* AGPs. The antibody LM2 (Smallwood *et al*., 1996, Figure 2C), identifying (1→6)-β-D-Gal*p* units with terminal β-D-Glc*p*A in AGPs (Ruprecht *et al*., 2017), reacted strongly with both bryophyte AGPs, but even more with *Anthoceros* AGP. Only the AGP of *Physcomitrium* showed a moderate binding to the antibody LM6 (Figure 2D) which detects oligomers with (1→5)-α-L-Ara*f* linkages in arabinans or AGPs (Verhertbruggen *et al*., 2009). Thus, the ELISA experiments underlined the structural differences between the hornwort and the moss AGPs (Figure 1).

**Figure 2.**
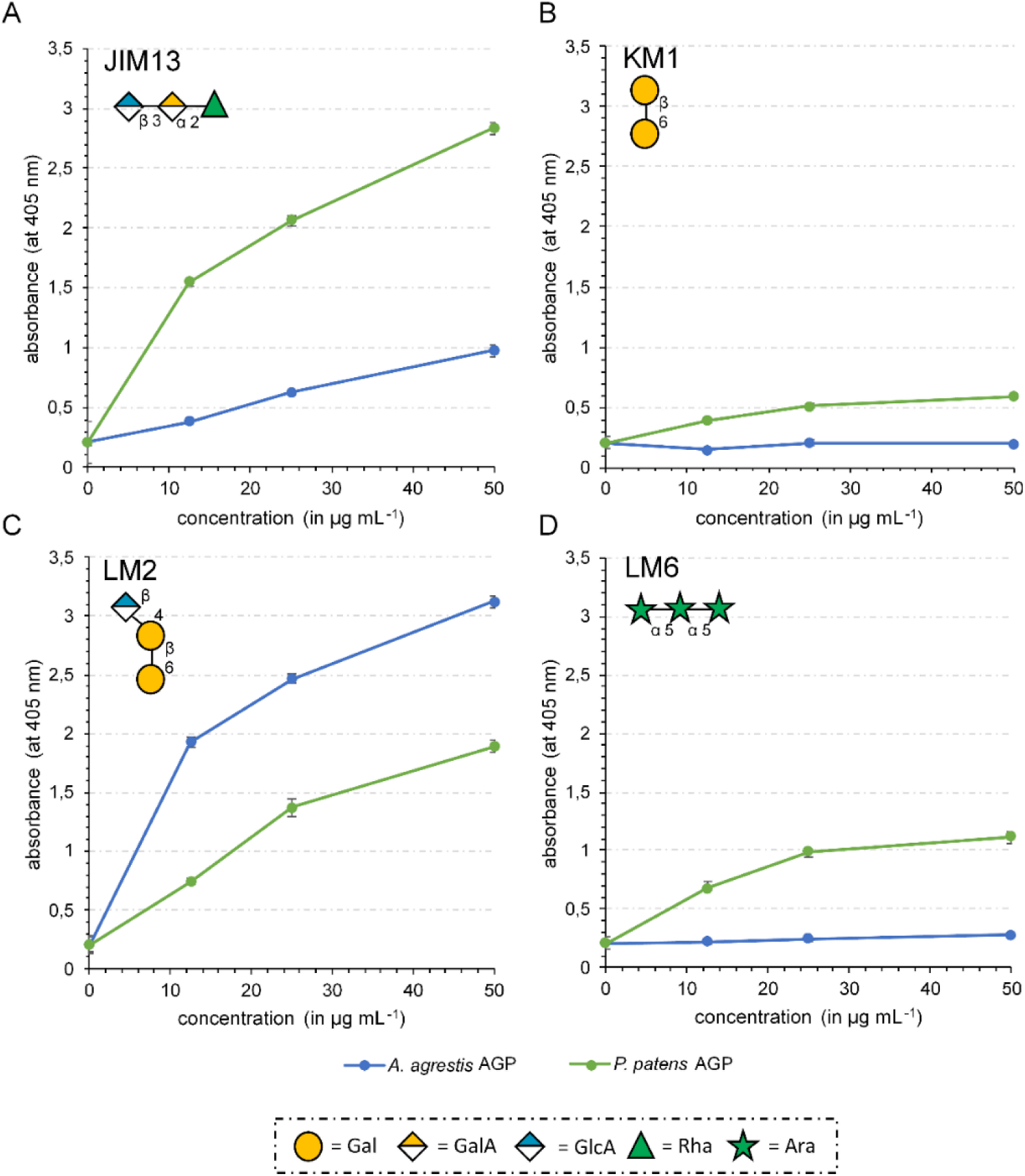
Reactivity of AGPs of *Anthoceros agrestis* (blue) and *Physcomitrium patens* (green) with antibodies against epitopes of AGP glycans tested by ELISA. **A**: JIM13; **B**: KM1; **C**: LM2; **D**: LM6. For epitopes of the antibodies, see Table S3.

### Enzymatic toolkit for AGP glycosylation in bryophytes

Twelve hornwort genomes (Schafran *et al*., 2025) as well as the genomes of *Marchantia* p*olymorpha* (Bowman *et al*., 2017) and *Physcomitrium patens* (Rensing *et al*., 2008, Lang *et al*., 2018) were searched for enzyme homologs involved in AGP glycosylation (Figure 3, Table S4, Data S1-12). As described above, glycosylation of AGPs is very complex, including a galactan framework attached to the protein moiety *via* Hyp and further decoration of the galactan with mainly Ara and different other monosaccharides like GlcA, Rha, Fuc or Xyl (for reviews, see Showalter & Basu, 2016; Silva *et al*., 2020; Strasser *et al*., 2021). Prolyl-4-hydroxylases (P4Hs) are responsible for hydroxylation of proline. Phylogenetic analysis revealed the presence of three to five putative AGP-related P4H homologs in hornwort genomes; *Marchantia* contained only two, whereas *Physcomitrium* had six (Data S1). Furthermore, we searched for glycosyltransferases (GTs, Data S3-7) and glycosylhydrolases (GHs, Data S8-12) involved in AGP biosynthesis and remodelling, which are classified according to carbohydrate-active enzyme nomenclature (CAZy; http://www.cazy.org). Several members of the GT31 family are described to be responsible for AGP galactosylation. *Arabidopsis* GALT 2-6 (Basu *et al*., 2015a,b) as well as HPGT1-3 (Ogawa-Ohnishi & Matsubayashi, 2015) are responsible for the first step of AGP glycosylation: the transfer of the initial Gal onto Hyp. Genomes of all hornworts, *Marchantia* and *Physcomitrium* had corresponding sequences in comparable numbers (three to five sequences). GALT31A, UPEX1/KNS4, GALT9 (family GT31) and GALT 29A (family GT29) are involved in synthesis of the 1,3-, 1,6-linked galactan framework in *Arabidopsis*, and all bryophyte genomes contained homologs of these sequences in comparable numbers (one to four sequences, with a high number of 4 GALT31A, UPEX1/KNS4 homologs in *Physcomitrium*).

**Figure 3.**
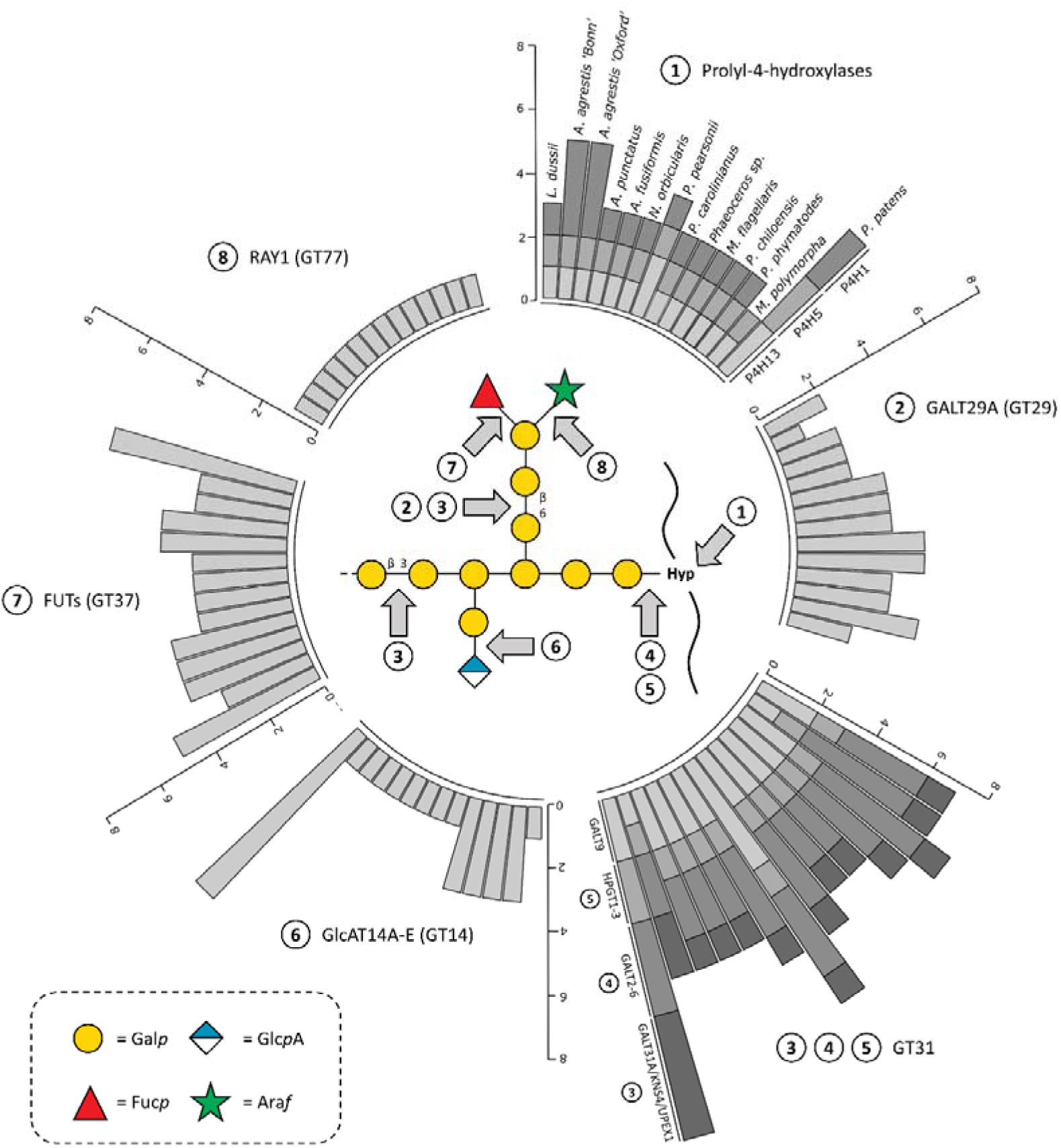
Circular stacked bar plot showing homolog numbers of characterized enzymes involved in AGP biosynthesis. A hypothetical model of AGP structure based on published enzyme specificities is shown in the middle. Grey arrows represent linkages resulting from the respective enzyme activities. Hornworts: *Leicosporoceros dussii*, *Anthoceros agrestis* ‘Bonn’, *Anthoceros agrestis* ‘Oxford’, *Anthoceros punctatus*, *Anthoceros fusiformis*, *Notothylas orbicularis*, *Paraphymatoceros pearsonii*, *Phaeoceros carolinianus*, *Phaeoceros* sp., *Megaceros flagellaris*, *Phaeomegaceros chiloensis*, *Phymatoceros phymatodes*; liverwort: *Marchantia polymorpha*; moss: *Physcomitrium patens*. For phylogenetic trees of all enzyme families, see Data S1-12.

Members of GT14 family of glucuronosyltransferases are acting on AGPs, especially GlcAT14A-E, which adds terminal Glc*p*A to the AG moieties. Homologs were detected in all bryophyte genomes with lower numbers in hornworts and the liverwort (one to three homologs) compared to *Physcomitrium* (7 homologs).

Arabinosyl- and rhamnosyltransferases responsible for decoration of the galactan backbone of AGPs are mainly unknown up to now (Silva *et al*., 2020). Only one homolog of the β-arabinofuranosyltransferase RAY1 (Gille *et al*., 2013) is present in all investigated hornwort and setaphyte genomes. No enzyme homologs of AGM1+2 occur in bryophyte genomes (Data S2). This corresponds to our biochemical data, as these enzymes transfer methylethers at C4 of GlcA in *Arabidopsis* AGPs (Temple *et al*., 2019) and we did not find any methylated GlcA in bryophyte AGPs. Homologs of GT37 occur in all bryophytes and comprise fucosyltransferases. As bryophyte AGPs contain no or only traces of Fuc (see Tables 1 and 2), members of the GT37 family possibly mainly act on other polysaccharides in bryophytes.

GHs known to be involved in AGP remodelling are RsAraf (GH3), AGAL2-3 (GH27), AtBGAL8 (GH35), AtGUS2 (GH79) and members of GH43. Homologous protein sequences coding for these enzymes were present in all bryophytes.

### Protein moiety of bryophyte AGPs

Since AGPs consist of a protein moiety covalently linked *via* Hyp to the carbohydrate part, the protein and the Hyp content of the bryophyte AGPs were determined. From the nitrogen content measured by elemental analysis (2.6 % for *Anthoceros* and 2.7 % for *Physcomitrium* AGP) the amount of protein in the AGPs was calculated using the Kjeldahl factor (x 6.25) and determined to be 16.3 % and 16.9 % of the dry weight of the AGPs, respectively. The Hyp amount of the *Anthoceros* AGP was slightly higher (0.31 %) compared to that of *Physcomitrium* (0.25 %). This results in a Hyp proportion of 1.9 % of the protein in *Anthoceros* AGP and 1.5 % for that of *Physcomitrium*.

### AGP protein sequences present in bryophyte genomes

Bryophyte genomes were also used to search for genes coding for AGP protein backbones. Sequences for classical AGPs with and without GPI-anchor as well as hybrid AGP sequences with additional motifs from other groups of hydroxyproline-rich glycoproteins (HRGPs) were identified (Figure 4A, Table S5) using the established motif and amino acid bias ‘MAAB’ classification system (Johnson *et al*., 2017a,b). It is striking that 10 of the hornwort genomes completely lack sequences for GPI-anchored classical AGPs; only the two genomes of *Leicosporoceros dusii* and *Anthoceros punctatus* contain one of these sequences. This is in strong contrast to the liverwort *Marchantia polymorpha* (20 sequences) and the moss *Physcomitrium patens* (16 sequences). All hornworts with the exception of *Leicosporoceros dusii* possess genome sequences for AGPs without GPI-anchor (1 to 7 sequences, in mean 2.9 sequences). These GPI-free AGPs are also more abundant in the setaphytes *(Marchantia* with 9 sequences; *Physcomitrium* with 18 sequences). Only few hybrid AGPs with an additional extensin or PRP motif were present in the investigated bryophyte genomes. The hornworts *Notothylas orbicularis*, *Phaeoceros* sp. and *Phaeoceros carolinianus* as well as *Marchantia* and *Physcomitrium* had one or two sequences, whereas all other hornwort genomes lacked hybrid AGP sequences.

**Figure 4.**
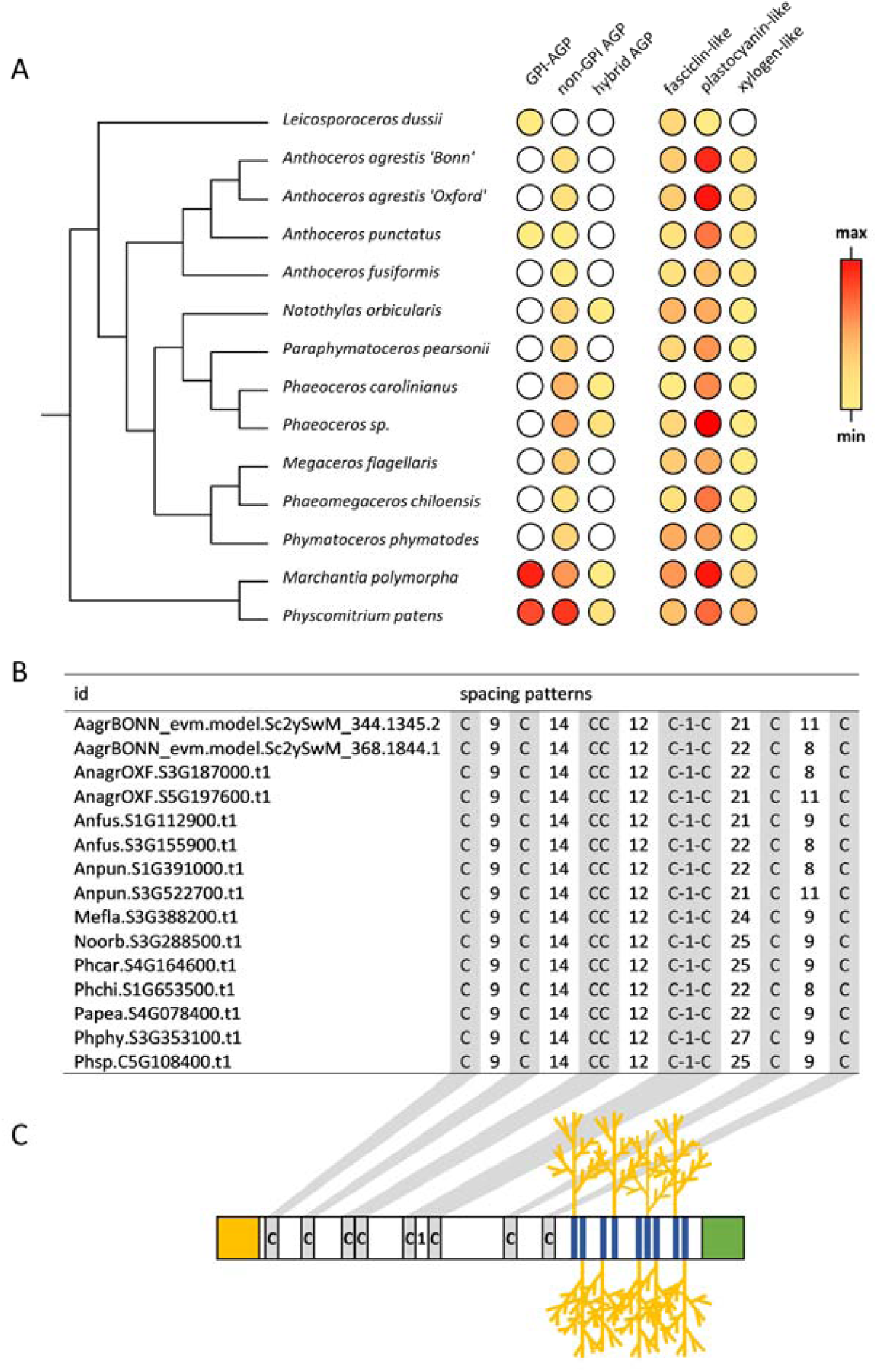
Identified AGP protein backbone sequences in 14 bryophyte genomes (12 hornworts, 1 liverwort and 1 moss). **A**: Heatmap representation of numbers for classical AGPs (GPI-anchored, non-GPI-anchored), hybrid AGPs, as well as selected chimeric AGPs (fasciclin-like, plastocyanin-like, and xylogen-like). **B**: Detailed structural analysis of spacing patterns between the eight conserved cysteine residues in the nsLTP2-domain (pfam14368) being characteristic for xylogen-like chimeric AGPs. **C**: Illustration of xylogen-like AGPs with the conserved cysteine residues shaded in grey, AG-glycomotifs shaded in blue. The *N*-terminal signal peptide and a GPI-anchor sequence are represented as yellow and green squares, respectively.

In chimeric AGPs (Figure 4A, Table S6), AGP motifs are accompanied by a signal peptide and one or more known functional protein domains (as classified in the pfam database, http://pfam.xfam.org/). Typical chimeric AGPs known from angiosperms were detected in the bryophyte genomes, namely plastocyanin-like AGPs (containing the Cu bind-like domain, pfam02298), fasciclin-like AGPs (containing the FAS1 domain, pfam02469) and xylogen-like AGPs (containing the LTP-2 domain, pfam14368; Ma *et al*., 2017). Most abundant in all bryophytes were the plastocyanin-like AGPs with in mean 11.2 sequences in hornworts, 21 sequences in *Marchantia* and 13 sequences in *Physcomitrium* (Figure 4A, Table S6). In comparison to the hornworts (range 1-7, in mean 3.4 sequences,), the fasciclin-like AGPs were slightly more abundant in *Marchantia* (9 sequences) and *Physcomitrium* (5 sequences). Within the bryophytes, xylogen-like AGP sequences dominated in the setaphytes (*Marchantia* with 3 sequences, *Physcomitrium* with 6 sequences); but some also occurred in the hornworts (one or two sequences, no sequence in *Leicosporoceros*). Xylogen-like AGP sequences contain lipid-transfer-like domains (Figures 4B and C), which only occur in land plants and are very stable due to four disulfide bridges formed by eight conserved cystein residues (Edstam *et al*., 2014). Detailed analyses of this non-specific lipid transfer protein (nsLTP) domain was performed by looking at the spacing patterns between the eight conserved cysteines (Figure 4B, Table S7) together with a check for putatively present GPI-anchors. These analyses revealed presence of only the G and G/D types (Figures 4B and 4C; for detailed classification of nsLTPs, see Liu *et al*., 2015). Further comparison with other genomic data (2 angiosperms, 2 gymnosperms, 3 ferns, 2 lycophytes, *Physcomitrium* and *Marchantia*) confirmed sole occurrence of these domain types (Table S8) in xylogen-like AGPs. G-type LTPs were initially classified by presence of GPI-anchors as described in Edstam *et al*. (2011). It has to be noted that the sequences labelled with “G/D” could possibly be G-type LTPs lacking these anchor structures.

### Immunocytochemical detection of *Anthoceros* AGPs by TEM

The JIM 13 antibody was also used to detect AGPs immunocytochemically in mature *Anthoceros* tissue (Figure 5 and Figures S3, S4). For better orientation, Figure S3A shows a detail of a plasmodesmos (asterisk) traversing the cell wall (CW) of *Anthoceros* thallus cells with complete structural preservation of e.g. the bilayered plasma membrane due to conventional fixation which, however, hinders immunoreactivity. To estimate the specificity of the immuno-labelling, TEM micrographs in Figure 5 and Fig. S3B-D, showing thalli (fixed for immunocytochemistry and treated with the primary antibody JIM 13 and gold-conjugated secondary antibodies), were compared to corresponding negative controls lacking JIM13 treatment (Figures S3E-H and Figure S4).

**Figure 5.**
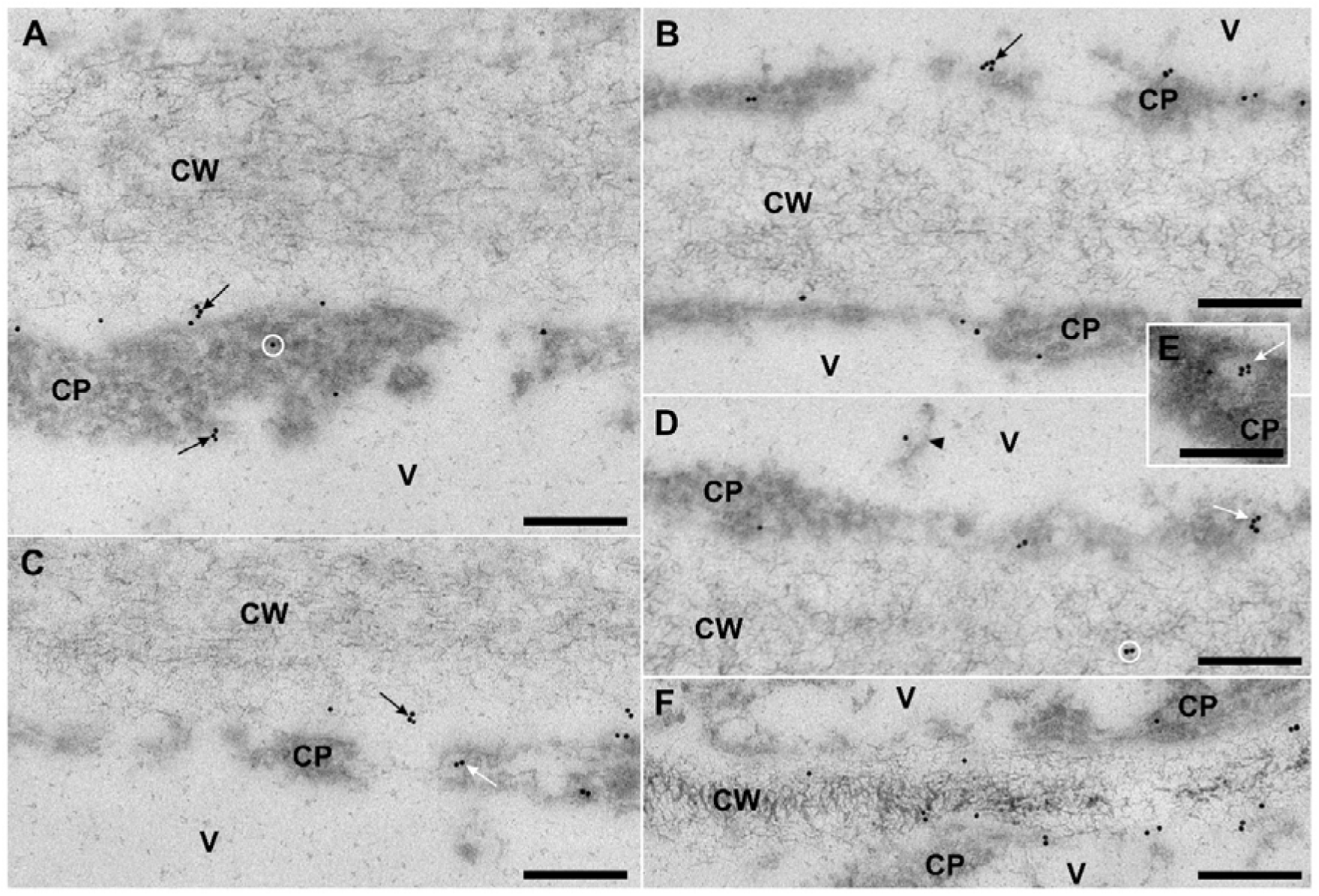
Immunolabeling of AGPs detected with JIM13 primary antibodies and 10 nm gold-conjugated secondary antibodies in epidermal cells of *Anthoceros agrestis* gametophyte thalli observed by transmission electron microscopy. **A**: Single gold labels and small clusters of 2-5 gold particles (black arrows) were regularly found at the plasma membrane bordering the outer, externally oriented epidermis cell wall (CW), but also at the innermost, youngest layers of that wall, and at the tonoplast membrane of the vacuole (V). Single labels were occasionally found in the cytoplasm (CP, circle). **B**: Cell wall between two adjacent cells with particular strong labeling of AGPs at the tonoplast (arrow). **C**: Many labels at the border between the old central wall layers, which have a fibrillar appearance, and the youngest electron-lucent wall layers closer to the plasma membrane (black arrow). The white arrow marks label on a vesicle. **D,E**: Rarely, labeling occurred in the older cell wall layers (white circle in **D**). Vesicles and their membranes in particular were frequently labeled (white arrows), as well as a dark material inside the vacuoles (arrowhead in **D**). **F**: Younger, still developing cell walls exhibited a higher immunoreactivity and labels also occurred in the central fibrillar wall layers. Scale bars: 200 nm, CW: cell wall, CP: cytoplasm, V: vacuole. Corresponding negative controls without JIM13 primary antibody treatment are shown in Figure S3.

Specific AGP labeling frequently appeared at the youngest layers of the outer epidermal wall (upper arrow, Figure 5A), of intercellular contact walls (Figures 5B,C,F, Figure S3B) and of cell walls bordering on intercellular spaces (IC; Figure S3C). The border between these youngest, electron-lucent wall layers close to the plasma membrane and the old central wall layers, which have a fibrillar appearance, also had a strongly immunoreactivity (black arrow in Figure 5C, Figure S3B). Rare single gold labels, which most likely represent the non-specific background, also appeared at the central layers of the wall (white circle in Figure 5D) and in the cytosol/cytoplasm (CP; white circle in Figure 5A). In the central fibrillar layers of young and still developing cell walls, however, immunoresponses appeared regularly (Figure 5F) which indicates a distinct distribution of AGPs in young and mature cell walls. Plasmodesmata (asterisk in Figure S3B) and their directly adjacent cell wall areas were never labelled.

Furthermore, specific AGP labels were found at the plasma membrane (Figure 5, black arrow in Figure S3C), which is a typical location of GPI-anchored AGPs, but also at the tonoplast of the vacuoles (V, black arrows in Figures 5A,B, Figure S3B), at vesicles and their membranes (white arrows in Figure 5C,D,E), as well as on a dark and presumably proteinaceous material inside the vacuoles (arrowhead in Figure 5D, Figure S3D).

## DISCUSSION

The step of terrestrialization, about 470 million years ago, with its significant change in environmental conditions, forced plants to develop coping strategies (Domozych & Bagdan, 2022). Comparative studies on extant bryophytes and their direct sister group, the tracheophytes, enable today’s research to gain more insights into strategies which had already evolved in their MRCA (Bechteler *et al*., 2023). Cell wall composition was most likely one of the features which had been strongly influenced during this process. There is a lack in knowledge on bryophyte AGPs when it comes to analytical characterization exceeding immunocytochemical screening of these important signalling molecules. In order to fill this gap, *Anthoceros* and *Physcomitrium* were analyzed in this study, focussing on in-depth analysis of the AGPs which represent a major class of signalling glycoproteins in plant cell walls.

### Yield of AGPs in bryophytes

In the past, AGPs have been detected in different bryophytes, but mostly on the basis of gel diffusion assays or antibody reactivities (Clarke *et al*., 1978; Basile & Basile, 1987; Ligrone *et al*., 2002; Kremer *et al*., 2004; Lee *et al*., 2005; Berry *et al*., 2016). Only a few bryophyte AGPs have been isolated, three from mosses (Lee *et al*., 2005; Fu *et al*, 2007; Bartels *et al*., 2017) and one from a liverwort (Happ & Classen, 2019). In this study, the first AGP from a hornwort has been characterized and compared to an AGP from *Physcomitrium*.

The AGP yield of *Physcomitrium* (0.09 %) is comparable to that of other bryophytes and ferns (0.06 % – 0.19 %; Bartels & Classen, 2017; Bartels *et al*., 2017; Happ & Classen, 2019; Mueller *et al*., 2023), while the AGP yield of *Anthoceros* is higher (0.31 % for plants, 0.57 % for cell culture). This feature might be specific for the species or for hornwort AGPs. Another possibility is a dependence on culture conditions, as *Anthoceros* plants were cultivated on agar medium and not on soil (Figure S4) and AGP yield from *Anthoceros* suspension cultures was highest. AGP content in *Physcomitrium* also increased when plants were grown in liquid culture (0.23 %; Bartels *et al*., 2017).

### Carbohydrate part of AGPs

The polysaccharide part of the AGP of *Anthoceros* mainly consists of Ara, Gal and 3-*O*-MeRha, which together make up over 90 %. In contrast, these monosaccharides account for only 70 % in *Physcomitrium* AGP, due to a high content of Glc. The high Glc content confirms the results obtained by other authors (Fu *et al*., 2007), but is contradictory to other studies, in which *Physcomitrium* AGP contained only small amounts of Glc (Lee *et al*., 2005, Bartels *et al*., 2017). Similar to our study, linkage-type analyses revealed dominance of 1,4-Glc in α-anomeric linkage, indicating that Glc is principally starch, that was resistant against starch-removal by DMSO and persisted through the precipitation with βGlcY (Fu *et al*., 2007). This result is unexpected, as normally βGlcY yields pure AGP fractions and no report is known of covalent linkages of AGPs to starch. Preliminary results from enzyme-assisted release of glucose with following quantification using a glucose-oxidase test with colorimetric detection also hinted towards presence of starch in *Physcomitrium* (Figure S2).

The typical galactan core of angiosperms consists of 1,3-, 1,6- and 1,3.6-linked Gal*p*. In the bryophyte AGPs, mainly 1,3- and 1,3,6-linked Gal*p* were present, accompanied by only low amounts of 1,6-linked Gal. Low amounts of 1,6-Gal*p* linkages were also characteristic for other AGPs of spore-producing land plants and indicate a high degree of branching of these AGPs (Lee *et al*., 2005; Bartels & Classen, 2017; Bartels *et al*., 2017; Happ & Classen, 2019; Mueller *et al*., 2023). Only the structure of the investigated peatmoss *Sphagnum* sp. AG moiety is an exception with a content of 1,6-Gal*p* linkages in the range of seed plants (Bartels *et al*., 2017; Thude & Classen, 2005). Further Gal branching points have been detected in bryophyte AGPs, e.g. 1,3,4-Gal*p* in the hornwort *Anthoceros* (this study) and 1,2,3-Gal*p* in the moss genera *Polytrichastrum* and *Sphagnum* (Bartels *et al*., 2017).

In angiosperm AGPs, Ara*f* is the dominant terminal residue, accompanied by terminal (4-*O*-Me)Glc*p*A, Gal*p* and sometimes Fuc*p* (Tryfona *et al*., 2012). Ara*f* is also the dominant terminal residue in *Anthoceros* and *Marchantia* (Happ & Classen, 2019) AGPs, but in *Physcomitrium*, the unusual terminal (3-*O*-Me)Rha*p*, which is also present in the *Anthoceros* and *Marchantia* AGPs, is the main terminal monosaccharide (see also Lee *et al*., 2005). Terminal 3-*O*-MeRha*p* has previously been found in cell walls and also AGPs of algae, bryophytes, ferns and gymnosperms (Akiyama *et al*., 1987; Ogawa *et al*., 1997; Popper & Fry, 2003; Pfeifer *et al*., 2022b; Bartels & Classen, 2017; Bartels *et al*., 2017; Happ & Classen 2019; Baumann *et al*., 2021; Mueller *et al*., 2023), but it has obviously been lost during evolution of angiosperms, since it has never been detected in any cell wall component of angiosperm species.

In angiosperm AGPs, Ara*f* is further present in linear linkage types, especially in 1,5-linkage (Classen *et al*., 2000; Thude & Classen, 2005, Wack *et al*., 2005). This is also the case for the *Physcomitrium* AGP in our study and in correspondence to other investigations on *Physcomitrium* (Lee *et al*., 2005; Fu *et al*., 2007), *Sphagnum* (Bartels *et al*., 2017) and *Equisetum* (horsetail; Bartels & Classen, 2017) AGPs. In contrast, the AGP of *Anthoceros* revealed an 1,3-Ara*f* linkage type similar to *Marchantia* and *Polytrichastrum* AGPs (Happ & Classen, 2019; Bartels *et al*., 2017). The 1,2-linkage type of Ara*f*, which occurs mainly in leptosporangiate ferns (Bartels & Classen, 2017; Mueller *et al*., 2023), was detected only in traces in AGPs of both bryophyte species investigated in the present study.

Results obtained by ELISA with several antibodies raised against epitopes of AGP-glycans support the analytical findings. The antibody JIM13 is commonly used to detect AGPs in plant tissue (e.g. Knox *et al*., 1991) The exact epitope is still unknown, but the trisaccharide β-D-Glc*p*A-(1→3)-α-D-Gal*p*A-(1→2)-α-L-Rha has been shown to bind this antibody (Yates *et al*., 1996). Furthermore, a rhamnogalactan-protein from *Spirogyra* algae (Zygnematophyceae) immuno-reacted with JIM13, underlining that Rha is possibly an integral part of the epitope (Pfeifer *et al*., 2022b). Since the AGP of *Physcomitrium* showed a higher affinity compared to that of *Anthoceros*, it is furthermore supported that Rha in particular is involved in binding. The low amounts of 1,6-Gal*p* linkages were verified by the results with the antibody KM1, which did not bind to the *Anthoceros* AGP and showed only weak interaction with *Physcomitrium* AGP. The epitope of this antibody was raised against an AGP of the angiosperm *Echinacea purpurea*, which is rich in 1,6-Gal*p* linkages (Classen *et al*., 2004). The presence of 1,5-Ara*f* specifically in *Physcomitrium* was confirmed by affinity of this AGP to LM6. The AGP of *Anthoceros* did not bind, confirming that other Ara*f* linkage types are present. LM6 was also used in the study of Lee *et al*. (2005). There, LM6 had a higher affinity to protonemal cells of *Physcomitrium* than JIM13. In particular, LM6 showed binding to the plasma membrane and cell wall surfaces at the tip of the growing protonemata, indicating that AGPs are involved in apical cell expansion. The antibody LM2 recognizes a carbohydrate epitope of an AGP from *Oryza sativa* (rice) rich in Ara and Gal and minor amounts of GlcA. As monomeric D-GlcA was effective to inhibit LM2 binding (Smallwood *et al*., 1996) and a Gal-oligosaccharide with a terminal GlcA showed effective binding to LM2 (Ruprecht *et al*., 2017), a combination of Gal with a terminal GlcA is possibly responsible for strong LM2 binding of both AGPs investigated in this study.

### Enzymes involved in biosynthesis of AGP glycans

The prerequisite for glycosylation of AGP protein backbones is proline hydroxylation. Presence of sequences coding for P4Hs in all bryophyte genomes revealed that this step is possible in hornworts, liverworts and mosses. Genes for Hyp-*O*-galactosyltransferases, β-1,3- galactosyltransferases and a β-1,6-galactosyltransferase are also present in the investigated bryophyte genomes. This corresponds to our biochemical investigations (see above) and confirms that the typical AGP galactosylation machinery of seed plants (Silva *et al*., 2020) and ferns (Mueller *et al*., 2023) acts similarly in bryophytes. In contrast to this conserved nature of the galactan framework, there is strong variation in monosaccharides present at the periphery of AGP molecules. *Anthoceros* and *Physcomitrium* AGPs contain small amounts of GlcA, which are added to Gal chains by GlcATs of CAZy family GT14 (Knoch *et al*., 2013; Dilokpimol & Geshi, 2014) for which genes are also present in genomes of the investigated hornworts, *Marchantia* and *Physcomitrium*. Unfortunately, only one AraT (RAY1, member of family GT77) working on AGPs has been described up to now (Gille *et al*., 2013). One single close homolog of RAY1 has been detected in all investigated bryophyte genomes and may be responsible for arabinosylation of bryophyte AGPs. Although *Arabidopsis* knockout mutants of RAY1 showed reduced Ara content, the influence was strongest for 1,3-linked Ara compared to other linkage types (Gille *et al*., 2013). As *t*-Ara is more dominant in bryophytes and *Physcomitrium* AGP has mainly 1,5-linked Ara residues, further AraTs must be involved in arabinosylation of AGPs. To date, rhamnosyltransferases acting on AGPs are completely unknown. Bryophyte AGPs (especially *Physcomitrium* AGP) contain terminal Rha, therefore search for these enzymes is an interesting task for the future. Furthermore, Rha is partially methylated at O-3 in bryophyte AGPs; thus, a further challenge will be search for the corresponding methyltransferases. In *Arabidopsis* AGP, the enzymes AGM1+2 belonging to the DUF579 family are responsible for methylation of terminal GlcA (Temple *et al*., 2019). Bryophytes did not contain methylated GlcA and also close homologs of *AGM1+2* are lacking in their genomes. Other members of the DUF579 family are present (Data S2) and might be responsible for methylation of Rha in bryophyte AGPs.

### Protein moiety of AGPs

The moss and the hornwort AGPs contain a similar protein content of around 16 % – 17 % (*w*/*w*), comparable to the protein content of the moss *Polytrichastrum* (Bartels *et al*., 2017) and two water ferns, *Azolla* and *Salvinia* (14.6 % and 19.4 %; Mueller *et al*., 2023). Other studies revealed a high variety of the protein amount in bryophyte AGPs ranging from 4.6 % (*Sphagnum*; Bartels *et al*., 2017) to 24.9 % (*Marchantia*, Happ & Classen, 2019). With exception of the moss *Sphagnum*, the protein content of bryophyte AGPs seems to be higher compared to angiosperms. In addition, the Hyp amount relative to the total protein of *Physcomitrium* and *Anthoceros* AGPs was in the same range compared to the bryophytes and aquatic ferns mentioned above (0.9 % – 2.5 %; Bartels *et al*., 2017; Happ & Classen, 2019; Mueller *et al*., 2023) and lower compared to angiosperms. The only exception was *Sphagnum* AGP with 5.8 % Hyp (Bartels *et al*., 2017). Apparently, a higher protein content results in lower amounts of Hyp, whereas a lower protein content requires higher portions of Hyp to ensure enough binding positions for the AG moieties.

### Bioinformatic search for AGP protein backbones in bryophytes

All hornwort genomes as well as those of *Marchantia* and *Physcomitrium* contain sequences coding for AGP protein backbones. The protein backbones of AGPs are rich in the amino acids hydroxyproline/proline (Hyp/Pro), alanine (Ala), serine (Ser) and threonine (Thr), which are regularly arranged as Ala–Pro, Ser–Pro, and Thr–Pro (for review, see Ma *et al*., 2018).

Classical AGPs occur without and with GPI-anchors, which link AGPs to the plasma membrane and facilitate co-location of signalling partners in lipid microdomains (Konrad & Ott, 2015). Our results reveal an impressive difference between hornworts and the setaphytes: GPI-AGPs are mostly missing in hornwort genomes (ten species without sequence, 2 species with one sequence), whereas *Marchantia* and *Physcomitrium* genomes contain 20 and 16 sequences, respectively. This is comparable to *Arabidopsis* with 17 sequences coding for GPI-AGPs (Johnson *et al*., 2017a). The low content of GPI-AGPs in hornworts compared to liverworts and mosses and also lycophytes, ferns, gymno- and angiosperms has also been shown for transciptomes (Johnson *et al*., 2017b). The same trend is obvious for AGPs without GPI-anchor: less sequences are detected in the hornwort genomes (in mean 2.9 sequences; range 0-7) compared to *Marchantia* (9 sequences) and *Physcomitrium* (18 sequences), the latter being in the range of *Arabidopsis* (12 sequences). To date, monophyly of hornworts, liverworts and mosses with hornworts as sister to the setaphytes is generally accepted (Puttick *et al*., 2018; Donoghue *et al*., 2021; Su *et al*., 2021). As genes for GPI-AGPs have already been detected in transcriptomes of green algae (Johnson et al., 2017a) and have obviously been present in the common ancestor of all bryophytes, it seems that they have been mostly lost in extant hornworts.

AGP protein backbones are characterized by high heterogeneity. Besides classical AGPs discussed above, hybrid AGPs combine AGP domains with sequences of other HRGPs (e.g. AGP domain + extension domain), and chimeric AGPs are characterized by one or more additional functional protein domains (Ma *et al*., 2017). Their gene homologs are also present in at least some of the investigated members of all bryophyte taxa, with chimeric AGPs being clearly more abundant and widespread. Fasciclin-like AGPs contain a fasciclin domain capable of binding multiple ligands and often involved in cell adhesion (Seifert, 2021). In some streptophyte algae, it has been shown that AGPs most likely function as key adhesion molecules (Palacio-Lopez *et al*., 2019; LoRicco *et al*., 2024). As adhesion was necessary for a sessile way of life, it was probably also a key mechanism for plant terrestrialization. Fasciclin-like AGPs are present in all investigated bryophyte genomes, and therefore they might have been important features for the ancestors of extant bryophytes to cope with life on land.

Besides adhesion, a huge challenge during terrestrialization was adaptation to water deficiency. In the clade of tracheophytes (lycophytes, ferns and seed plants), long-distance transport of water occurs in highly specialized tracheary elements. Historically, it was speculated that transporting tissue has been an innovation in tracheophytes. Today it is known that simple conductive tissue is widespread among setaphytes, although occurrence and morphology is variable (Woudenberg *et al*., 2022). An involvement of AGPs in development of water-transporting tissue seems probable, as antibodies directed against AGP glycan epitopes bound to xylem elements of angiosperms (Dolan *et al*., 1995; Goellner *et al*., 2013). Some of these antibodies also labelled water-conducting cells of different liverworts and mosses (Ligrone *et al.,* 2002). Especially the chimeric xylogen-like AGPs are candidates possibly involved in development of water conducting tissue. The chimeric AGP called “xylogen” is responsible for xylem development in cell cultures of the angiosperm *Zinnia* (Motose *et al*., 2004). Sequences of xylogen-like AGPs were more abundant in *Marchantia* and *Physcomitrium* compared to the hornworts, which corresponds to the morphological finding that conductive tissues like water-conducting cells are absent in hornworts (Ligrone *et al*., 2000; Woudenberg *et al*., 2022). On the other hand, our finding that sequences of xylogen-like AGPs appear in hornwort, liverwort and moss genomes supports the hypothesis, that an evolutionary continuity is more likely than a strict gap between water-conducting tissue of bryophytes and vascular plants (Woudenberg *et al*., 2022). In this study, we were able to further classify the type of nsLTP domains included in the bryophyte xylogen-like AGPs, which are highly congruent. Our findings suggest that xylogen-like AGP evolution and evolution of nsLTP domains in general are deeply interconnected. This becomes evident when comparing our findings with the published data of Edstam *et al*. (2011). These authors proposed an evolutionary history of nsLTPs with D and G type domains in the MRCA of all land plants while all streptophyte algae lack nsLTPs. We found 4 G-type nsLTP domains in xylogen-like AGPs in *Marchantia* corresponding to a total of 4 G-type nsLTP sequences in *Marchantia* expressed sequence tags (ESTs, Edstam *et al*., 2011). Therefore, we hypothesize that all G-type nsLTP sequences were initially chimeric AGP sequences and a following diversification led to all other (non-chimeric) nsLTPs. This is also supported when looking at all tracheophyte genomes investigated in our study: all chimeric xylogen-like AGPs and also the prototypical xylogen from the angiosperm *Zinnia* are classified either as G-type or ambiguous G/D-types. Remarkable was also the observation that all organisms living in an aquatic habitat (*Azolla*, *Salvinia*, *Isoetes*) have generally lower numbers of xylogen-like AGPs (Table S8).

### Detection of AGPs in plant tissue of *Anthoceros* by TEM

In some immunohistochemical studies, JIM13 has already been used to detect AGPs in bryophytes. JIM13 labels *Physcomitrium* protonemata (Lee *et al*., 2005), and in protoplasts of *Marchantia*, AGPs in cell walls and cell plates were detected by JIM13 (Shibaya & Sugawara, 2009). Berry *et al*. (2016) detected epitopes of JIM13 at *Physcomitrium* protoplasts and gametophores. In addition, JIM13 epitopes were identified in secondary wall layers of the hornwort *Megaceros* and in hyaline cells of the peat moss *Sphagnum* (Kremer *et al*., 2004). A study on *Physcomitrium* apical cell (derivatives) by TEM revealed occurrence of JIM13 positive AGPs over all developmental stages, with a cellular distribution pattern predominantly in an electron-lucent layer close to the plasma membrane and at the wall / plasma membrane interface. During early development of gametophores, JIM13 was also present in the cytoplasm, especially close to ribosomes, but also vacuolar and tonoplast labeling was observed (Mansouri, 2012). In general, it is well-established knowledge that AGPs are cell wall glycoproteins which can be plasma membrane bound or secreted into the extracellular matrix (Ma & Johnson, 2023), including mucilages (Plachno *et al*., 2020) or media of suspension cultures (Classen, 2007). In contrast, their subcellular distribution is largely unknown. Due to the detailed investigation on JIM13 positive AGPs in *Physcomitrium* (Mansouri, 2012), we focussed on mature thallus segments of *Anthoceros* in our study.

Comparable with the results of Mansouri (2012), our results demonstrated the presence of AGPs at the plasma membrane and at the electron-lucent wall layers close to it. As no classical GPI-anchored AGPs were identified in *Anthoceros* (see above), chimeric AGPs (which are GPI-anchored) are possible responsible for this labelling at the plasma membrane. The median, fibrillar wall layers were labelled in young, still developing cell walls, but not in mature walls, which indicates a developmental modification of at least the AGP distribution. Remarkably, in both developmental stages, no AGPs could be detected at plasmodesmata or at the surrounding wall area, although it has been shown for an *Arabidopsis thaliana* mutant defective in arabinogalactan chain biosynthesis that AGPs play an essential role in plasmodesmal biogenesis (Okawa *et al*., 2023). It is conceivable, however, that the AGPs required for this function cannot be detected by the JIM13 antibody or differ between *Arabidopsis* and *Anthoceros*. Moreover, the two species differ drastically with respect to the plasmodesmal development (Wegner & Ehlers 2024).

We also observed a very prominent labelling of the tonoplast and of vesicle(membrane)s. Occurrence of AGPs at the tonoplast has already been reported for some angiosperm plants (Šamaj *et al*., 2000; Sala *et al*., 2017; Olmos *et al*., 2017). A study on maize (*Zea mays*) seedlings revealed association of AGPs with endomembranes, for example with the secretory compartments (endoplasmic reticulum and Golgi apparatus; Šamaj *et al*., 2000). As AGP biosynthesis happens in the ER and Golgi apparatus (Silva *et al*., 2020), this has to be expected, especially in young developing cells. During xylem development in *Echinacea purpurea*, AGPs were detected in Golgi vesicles of young tracheary elements close to the cambium (Goellner *et al*., 2013). In adventitious roots of tomato, AGP epitopes were abundantly present in cytoplasmic compartments; and the JIM13 epitope was localized at the tonoplast (Sala *et al*., 2017). In tobacco cell cultures, JIM13 AGP epitopes were located at the plasma membrane and also in the cytoplasm associated with the endoplasmic reticulum, the Golgi apparatus, vesicles and the tonoplast (Olmos *et al*., 2017). The authors propose a role of AGPs as sodium carriers through vesicle trafficking from the plasma membrane to the tonoplast during salt stress. Herman and Lamb (1992) supported the hypothesis that AGPs can move from the plasma membrane to the tonoplast, since the AGP epitope was detected in the plasma membrane, in invaginations of the plasma membrane, in intracellular multivesicular bodies (= endosomes) and finally in partially degraded multivesicular bodies sequestered within the central vacuole. Our study on *Anthoceros* supports this view, since we also observed immunoreactivity of all membranes involved and of the dark material deposited in the vacuoles. There might be still unknown functions of AGPs, especially connected with localisation at the tonoplast. These need not necessarily depend on classical GPI-anchored AGPs which lack in *Anthoceros agrestis* according to our findings.

## CONCLUSION

To date, only a few AGPs have been isolated from moss and liverwort cell walls. In this study, AGPs of a hornwort have been characterized for the first time and bioinformatic search for enzymes involved in AGP biosynthesis and for AGP backbone proteins has been performed for 12 hornwort genomes in comparison to *Marchantia* and *Physcomitrium* genomes. There were striking differences between the hornworts and the setaphytes, as sequences for AGP protein backbones were less abundant in hornworts compared to *Marchantia*, *Physcomitrium* and also *Arabidopsis*. This was extreme for GPI-AGPs, which were completely missing in ten of the twelve hornwort genomes. Sequences encoding glycosyltransferases involved in AGP galactosylation were present in all bryophyte genomes, and the biochemical investigations also revealed presence of the typical AGP galactan framework in *Anthoceros* and *Physcomitrium*. This underlines that Hyp-rich proteins highly glycosylated by branched galactans already occurred in the MCRA of all land plants. They were probably necessary for adaptation to life on land. This is supported by the presence of sequences encoding fasciclin-like AGPs in all bryophyte genomes, which are described to play a role in adhesion, a fundamental prerequisite for plants conquering land. Furthermore, all bryophyte genomes except one contained sequences of xylogen-like AGPs, indicating that – although bryophytes contain no xylem – one of the basal requirements for development of water-conducting tissues is already present in all bryophyte taxa. This includes also hornworts which have not evolved morphologically differentiated conductive tissues. A special feature of bryophyte AGPs and highly abundant in *Physcomitrium* is terminal 3-*O*-methylrhamnose. This unusual monosaccharide also occurs in fern and some gymnosperm AGPs, but not in angiosperms. Identification of the corresponding rhamnosyl- and methytransferases is a challenging task for the future. In TEM, AGPs were detected in *Anthoceros* tissue at the plasma membrane/cell wall interface, as known for many angiosperms. Additionally, AGPs were present at the tonoplast and at vesicles, suggesting new functions of AGPs. Generation of AGP mutants in the model organisms *Marchantia* or *Physcomitrium* will help to elucidate functions of AGPs in bryophytes.

## MATERIAL AND METHODS

### Plant material and sampling

The hornwort *Anthoceros agrestis* PATON (*Anthoceros*) was cultivated on agar medium in the Pharmaceutical Institute of Kiel University, Germany (Figure S5). Spores and fresh gametophyte material (Saxony strain formerly known as “Bonn”) were kind gifts of Péter Szövényi (University of Zurich). For spore germination and gametophyte culture on BCD medium (Cove *et al*., 2009), the protocol of Szövényi *et al*. (2015) was used. *Anthoceros agrestis* suspension culture material was a kind gift of Maike Petersen (University of Marburg). The material was cultivated in her lab according to Petersen (2003). The moss *Physcomitrium patens* (HEDW.) BRUCH & SCHIMP (*Physcomitrium*) was grown in the greenhouse of the garden of the Pharmaceutical Institute. Fresh gametophyte material of *Physcomitrium* “Reute” strain were kind gifts of Stefan A. Rensing (University of Freiburg). All samples were cleaned with water and freeze-dried.

### Isolation of water-soluble cell wall fractions and arabinogalactan-proteins (AGPs)

After freeze-drying and grinding of the plant material, it was pre-extracted two times (2 h and 21 h) with acetone-water (70 %, *V/V*) in a ratio of 1:10 (*w/V*) for *Anthoceros* and 1:20 (*w/V*) for *Physcomitrium*. The air-dried plant residue was extracted with double-distilled water (ddH_2_O) under constant stirring for 21 h at 4 °C in an 1:50 (*w/V*) ratio for *Anthoceros* and 1:20 (*w/V*) for *Physcomitrium*. Afterwards, the aqueous extract was separated from the insoluble plant material in case of *Anthoceros* by centrifugation (19,000 *g*, 4 °C, 20 min) or through a tincture press for *Physcomitrium*. The insoluble plant pellet was freeze-dried and used for further extractions (see below).

Proteins in the aqueous extract were removed by heating to 90 – 95 °C for ten minutes with following centrifugation (20 min, 4 °C, ≥ 4,122 g). The aqueous extract was then concentrated to approx. one-tenth and precipitated with ethanol (final concentration: 80 %, *V*/*V*) at 4°C, overnight. The resulting precipitate was separated by centrifugation (4 °C, 19,000 *g*, 20 min) and freeze-dried afterwards (yielding AE).

AGPs were isolated from the AE with the β-Glc-Yariv reagent (βGlcY; 1 mg mL^-1^) The precipitated AGP-βGlcY-complex was separated and purified to gain the aqueous AGP fraction according to the procedure of Classen *et al*. (2005) and Mueller *et al*. (2023).

### Analysis of monosaccharides

The determination of the neutral monosaccharide composition was performed by acid hydrolysis (2 mol L^-1^ TFA, 1 h, 121 °C) with following derivatization to the acetylated alditols. These were measured *via* gas chromatography (GC) with flame ionization detection (FID) and mass spectrometry detection (MSD; GC + FID: 7890B; Agilent Technologies, Santa Clara, CA, USA; MSD: 5977B; Agilent Technologies; column: Optima-225; Macherey-Nagel, Düren, Germany; 25 m, 250 µm, 0.25 µm; helium flow rate: 1 ml min^-1^; split ratio 30:1. A temperature gradient was used for peak separation (initial temperature 200 °C, subsequent holding time of 3 min; final temperature 243 °C with a gradient of 2 °C min^-1^). The methylated monosaccharide 3-*O*-MeRha was identified by retention time and mass spectrum (see Happ & Classen, 2019).

The uronic acid content (UA) was performed according to published literature (Blakeney *et al*., 1983; Blumenkrantz & Asboe-Hansen, 1973) with modifications as described by Mueller *et al*. (2023). Therefore, the uronic acids were derivated with *m*-hydroxydiphenyl and qualified photometrically at 525 nm by using a linear calibration line of a 1:1 GalA and GlcA mixture.

### Reduction of uronic acids of AGPs

Following the workflow of Taylor & Conrad (1972) with modifications described in Mueller *et al*. (2023), the uronic acids in 10 – 20 mg AGP sample were carboxy-reduced with 216 mg of *N*-cyclohexyl-*N*-[2-(N-methylmorpholino)-ethyl]-carbodiimide-4-toluolsulfonate and deuterium-labelled with sodium borodeuteride solutions (2.0 mL of 1 mol L^-1^; 2.5 mL of 2 mol L^-1^; 2.5 mL of 4 mol L^-1^) to yield the AGP_UR_-fraction).

### Structural characterization of AGPs

Linkage types of monosaccharides were determined using the methodology of Mueller *et al*., 2023, which is a modified version of Harris *et al*. (1984).

To achieve permethylation, the samples were first treated alternately with potassium methylsulfinyl carbanion and iodomethane in dimethyl sulfoxide and subsequently derivatized to permethylated alditol acetates (PMAA). These PMAAs were analysed by gas-liquid chromatography-mass spectrometry (see “Analysis of monosaccharides”; column: Optima-1701, 25 m, 250 µm, 0.25 µm; helium flow rate: 1 mol L^-1^; initial temperature: 170 °C; hold time 2 min; rate 1°C min^-1^ until 210 °C; rate: 30 °C min^-1^ until 250 °C; final hold time 10 min).

### Determination of hydroxyproline and protein content

The protein content (nitrogen content x 6.25; Kjeldahl, 1883) was determined by elemental analyses in the Chemistry Department of Kiel University, Kiel, Germany (HEKAtech CHNS Analyzer, Wegberg, Germany). Quantification of Hyp was performed photometrically at 558 nm according to Stegemann & Stalder (1967) with slight modifications (see Mueller *et al*., 2023). After acid hydrolysis (6 mol L^-1^, 22 h, 110 °C), the Hyp content was coupled with 4-dimethylaminobenzaldehyde and determined by linear regression analysis (Standard: 4-hydroxy-L-proline).

### Indirect enzyme-linked immunosorbent assay (ELISA)

ELISA experiments were performed as described in Mueller *et al*. (2023). The aqueous AGP solutions were used in the following concentrations: 12.5 µg mL^-1^, 25 µg mL^-1^ and 50 µg mL^-1^). The primary antibodies (JIM13, KM1, LM2, LM6; Kerafast, Inc., Boston; USA) were diluted 1:20 (*V/V*). The secondary antibodies coupled with alkaline phosphatase (all from Sigma-Aldrich Chemie GmbH, Taufkirchen, Germany), against rat-IgG or mouse-IgG (only for KM1) were used in a dilution of 1:500 (*V/V*) in phosphate-buffered saline (PBS; pH 7.4). Epitopes of the primary antibodies and key references are listed in Table S3.

### Gel diffusion assay

Cavities were stamped into an agarose gel (Tris-HCl, 10 mmol L^-1^; CaCl_2_, 1 mmol L^-1^; NaCl, 0.9 % *w/V*; agarose, 1 % *w/V*) and loaded with 20 µl of a dilution of each AE sample (100 mg ml^-1^) or βGlcY (1 mg ml^-1^), respectively. Red precipitation bands appeared in positive samples after diffusion overnight in the dark.

### Plant culture and immunocytochemical transmission electron microscopy (TEM)

For TEM, *Anthoceros agrestis* (‘Saxony’ strain) plants were grown on soil in a green house. Mature gametophyte thallus fragments were cut and embedded in 2 % low-gelling Agarose (type VII, Sigma-Aldrich, Steinheim, Germany). After fixing for 2h each, at room temperature and on ice, in 1 % (*w/V*) paraformaldehyde and 0.25 % (*V/V*) glutaraldehyde in 50 mmol mL^-1^ sodium phosphate-buffer; pH 7.2), and afterwards washing in buffer and ddH_2_O, the samples were contrasted for 2 h on ice in 0.5 % uranyl acetate. Following dehydration in a graded ethanol series on ice (30, 50, 70, 90 %), tissues were transferred into LR White hard grade (Agar Scientific, Stansted, UK), embedded in closed gelatine capsules and polymerized for 24 h at 50 °C. Ultra-thin sections (80 nm thick) were cut on a Reichert Om U2 ultramicrotome (Leica Microsystems GmbH, Wetzlar, Germany) and mounted on formvar-coated single-slot gold grids.

For immunolabelling, the sections were blocked for 1 h with 5 % BSA in 100 mmol mL^-1^ PBS (pH 7.2), incubated with the primary antibody JIM13 (Table S1), diluted 1:20 in 0.5 % BSA in PBS for 3 h, and treated with goat-anti-rat IgG (H+L) – 10 nm gold conjugate (Cytodiagnostics, Burlington, Canada), diluted 1:50 in 0.5% BSA in PBS for 1 h. Washing with PBS was done after both antibody steps. Finally, the sections were treated with 3 % glutaraldehyde for 10 min, washed again with ddH_2_O, and contrasted for 12 min each with 2 % uranyl acetate and lead citrate (Reynolds, 1963). Analyses were performed with an EM912AB transmission electron microscope (Zeiss, Oberkochen, Germany) at 120 kV acceleration voltage under zero-loss energy filtering conditions. A 2 k × 2 k dual-speed slow-scan CCD camera (SharpEye, TRS, Moorenweis, Germany) and the iTEM software package (OSIS) were used for image recording. The micrographs were mounted to figure plates with Corel PHOTO-PAINT (2021, Version 23.1.0.389, Corel Ottawa Canada).

### Bioinformatic search for AGP backbone sequences and genes involved in AGP biosynthesis

Twelve hornwort (Schafran *et al*., 2025), one liverwort (Bowman *et al*., 2017) and one moss (Rensing *et al*., 2008; Lang *et al*., 2018) genomes were searched for AGP protein backbones with the workflow described in Mueller *et al*. (2023). In brief, protein sequences containing a putative N-terminal signal peptide were initially identified through querying the SignalP 5.0 webserver (Almagro Armenteros *et al*., 2019). Those were then classified through the R-implemented maab-pipeline (Johnson *et al*., 2017a; Dragićević *et al*., 2020). Small (<90 amino acids), highly similar (≥ 95%) and sequences containing any pfam domain (E-value <1e-5) were excluded from that analysis. GPI-anchor prediction was performed with bigPI (Eisenhaber *et al*., 2003) and in case of ambiguities additionally with NetGPI 1.1 (Gíslason *et al*., 2021) and PredGPI (Pierleoni *et al*., 2008).

Chimeric AGP sequences including minimum one FAS1 (pfam02469), copper-binding like (pfam02298) or nsLTP (pfam14368) domain were identified by querying the pfam database *via* the CDD webtool (Lu *et al*., 2020). To classify a sequence as chimeric AGP, minimum one region containing at least 3 dipeptides ([STAGV]P) separated by maximally 6 amino acids, should be present beside the known protein domain. After classification, the nsLTP domain was cut out with an addition of 5 amino acids at each end. Those trimmed regions were aligned using MAFFT in L-INS-i mode (version 7.490; Katoh & Standley, 2013) and manually inspected using Jalview (Waterhouse *et al*., 2009). Types of LTP domains based on spacing patterns between the eight conserved cysteins as well as presence of GPI-anchors were classified according to Edstam *et al*. (2011).

The workflow for annotation of carbohydrate-active enzymes (CAZy) was performed as described in Ali *et al*. (2024) were identified using the dbCAN2 pipeline (Zhang *et al*., 2018) annotating them with three tools (HMMer against the CAZyme domain database, DIAMOND for BLASTp against the CAZyme database and dbCAN-sub for HMMer detection of putative CAZy substrates). All sequences classified as one GT or GH family by minimum 2 tools were filtered and aligned using MAFFT (preferably L-INS-i mode, otherwise FFT-NS-I mode; Katoh & Standley, 2013) or FastTree2 (for GT31 family; Price *et al*., 2010). Sequences similar to DUF579 and P4Hs of *Arabidopsis* were identified using BLASTp (E-value of e^-7^) and aligned using MAFFT.

For comparison, genomes of two angiosperms (*Arabidopsis thaliana*, Lamesch *et al*., 2011; *Amborella trichopoda*, Albert *et al*., 2013), two gymnosperms (*Cycas panzhihuaensis*, Liu *et al*., 2022; *Picea abies*, Nystedt *et al*., 2013), two lycophytes (*Isoetes taiwaniensis*, Wickell *et al*., 2021; *Selaginella moellendorffii*, Banks *et al*., 2011) and three ferns (*Azolla filiculoides*, *Salvinia cucullata*, Li *et al*., 2018; *Ceratopteris richardii*, Marchant *et al*., 2022) were used. Phylogenetic trees were inferred with IQ-TREE (version 1.6.12; Nguyen *et al*., 2015) with built-in model finder and visualized with iTOL (Letunic & Bork, 2021)

### Data statement

All relevant experimental data can be found within the manuscript and its supporting materials.

## Acknowledgements

The authors thank Maike Petersen (University of Marburg) for *Anthoceros agrestis* suspension culture material, Péter Szövényi (University of Zurich) for spores and gametophytes of *Anthoceros agrestis*, and Stefan Rensing (University of Freiburg) for fresh gametophyte material of *Physcomitrium* “Reute” strain. We acknowledge the Imaging Unit of the JLU Giessen Germany for providing access to the TEM facilities.

## Authorship contribution statement

**Kim-Kristine Mueller:** Conceptualization, Data curation, Formal analysis, Investigation, Visualization, Writing – original draft, Writing – review and editing.

**Lukas Pfeifer:** Conceptualization, Data curation, Formal analysis, Resources, Investigation, Visualization, Writing – original draft, Writing – review and editing.

**Linus Wegner:** Data curation, Formal analysis, Investigation, Visualization, Writing – review and editing.

**Kathrin Ehlers:** Conceptualization, Funding acquisition, Project administration, Writing – review and editing.

**Birgit Classen:** Conceptualization, Methodology, Funding acquisition, Project administration, Writing – original draft, Writing – review and editing.

## Supporting information

Additional Supporting Information can be found in the online version of this article.

**Table S1.** Neutral monosaccharide composition of water-soluble macromolecules from *Anthoceros agrestis and Physcomitrium patens* in % (mol mol^-1^).

**Table S2.** Content of uronic acids in water-soluble polysaccharides and AGPs of *Anthoceros agrestis and Physcomitrium patens* in % (w w^-1^).

**Table S3.** Antibodies directed against AGP glycan motifs used in this study.

**Table S4.** Homolog numbers of characterized AGP-active enzymes as identified by phylogenetic genome analysis of 12 hornwort and 2 setaphyte genomes.

**Table S5.** Numbers of identified protein sequences for classical (+/- GPI-anchor) and hybrid arabinogalactan-proteins.

**Table S6.** Numbers of identified protein sequences for chimeric arabinogalactan-proteins.

**Table S7** Detailed analysis of sequence characteristics in non-specific lipid transfer protein domains within xylogen-like AGPs of bryophytes.

**Table S8** Detailed analysis of sequence characteristics in non-specific lipid transfer protein domains within xylogen-like AGPs of selected other embryophytes.

**Figure S1.** Gel diffusion assay of aqueous extracts from *Anthoceros agrestis*, *Marchantia polymorpha* and *Physcomitrium patens* with βGlcY.

**Figure S2.** Starch analysis in *Physcomitrium patens* AGP.

Figure S3. Immunolabeling of AGPs detected with antibody JIM13 in epidermal cells of *Anthoceros agrestis* observed by TEM.

**Figure S4.** Negative controls corresponding to the immunolabeling of AGPs detected with antibody JIM13 in epidermal cells of *Anthoceros agrestis* observed by TEM.

**Figure S5.** Culture of *Anthoceros agrestis* on agar plates.

**Data S1.** Phylogenetic tree for prolyl-4-hydroxylases (P4Hs).

**Data S2.** Phylogenetic tree for DUF579 homologs.

**Data S3.** Phylogenetic tree for glycosyltransferase 14 (GT14) family members.

**Data S4.** Phylogenetic tree for glycosyltransferase 29 (GT29) family members.

**Data S5.** Phylogenetic tree for glycosyltransferase 31 (GT31) family members.

**Data S6.** Phylogenetic tree for glycosyltransferase 37 (GT37) family members.

**Data S7.** Phylogenetic tree for glycosyltransferase 77 (GT77) family members.

**Data S8.** Phylogenetic tree for glycosylhydrolase 3 (GH3) family members.

**Data S9.** Phylogenetic tree for glycosylhydrolase 27 (GH27) family members.

**Data S10.** Phylogenetic tree for glycosylhydrolase 35 (GH35) family members.

**Data S11.** Phylogenetic tree for glycosylhydrolase 43 (GH43) family members.

**Data S12.** Phylogenetic tree for glycosylhydrolase 79 (GH79) family members.

## REFERENCES

Albert, V.A., Barbazuk, W.B., de Pamphilis, C.W., Der, J.P., Leebens-Mack, J., Ma, H., et al. (2013) The Amborella genome and the evolution of flowering plants. Science, 342, 1241089.

Almagro Armenteros, J.J., Tsirigos, K.D., Sønderby, C.K., Petersen, T.N., Winther, O., Brunak, S., et al. (2019) SignalP 5.0 improves signal peptide predictions using deep neural networks. Nature Biotechnology, 37, 420–423.

Akiyama, T., Tanaka, K., Yamamoto, S. (1987) An arabinogalactan-rich protein containing 3-*O*-methylrhamnose (Acofriose) in young plants of *Osmunda japonica*. Agricultural and Biological Chemistry, 51, 2599–2600.

Ali, Z.M., Tan, Q.W., Lim, P.K., Chen, H., Pfeifer, L., Julca, I. et al. (2024) Comparative transcriptomics in ferns reveals key innovations and divergent evolution of the secondary cell walls. bioRxiv, doi: 10.1101/2024.08.27.609851

Banks, J.A., Nishiyama, T., Hasebe, M., Bowman, J.L., Gribskov, M., de Pamphilis, C., et al. (2011) The Selaginella genome identifies genetic changes associated with the evolution of vascular plants. Science, 332, 960–963.

Bartels, D., Baumann, A., Maeder, M., Geske, T., Heise, E.M., von Schwartzenberg, K., Classen, B. (2017) Evolution of plant cell wall: Arabinogalactan-proteins from three moss genera show structural differences compared to seed plants. Carbohydrate Polymers, 163, 227–235.

Bartels, D. & Classen, B. (2017) Structural investigations on arabinogalactan-proteins from a lycophyte and different monilophytes (ferns) in the evolutionary context. Carbohydrate Polymers, 172, 342–351.

Baumann, A., Pfeifer, L., Classen, B. (2021) Arabinogalactan-proteins from non-coniferous gymnosperms have unusual structural features. Carbohydrate Polymers, 261, 11783.

Basile, D.V. & Basile, M.R. (1987) The occurrence of cell wall-associated arabinogalactan proteins in the Hepaticae. The Bryologist, 90, 401–404.

Basu, D., Tian, L., Wang, W., Bobbs, S., Herock, H., Travers, A., Showalter, A.M. (2015a) A small multigene hydroxyproline-O-galactosyltransferase family functions in arabinogalactan-protein glycosylation, growth and development in Arabidopsis. BMC Plant Biology, 15, 295.

Basu, D., Wang, W., Ma, S., DeBrosse, T., Poirier, E., Emch, K., Soukup, E., Tian, L., Showalter, A.M. (2015b) Two Hydroxyproline Galactosyltransferases, GALT5 and GALT2, Function in Arabinogalactan-Protein Glycosylation, Growth and Development in Arabidopsis. PloS ONE, 10, e0125624.

Bechteler, J., Peñaloza-Bojacá, G., Bell, D., et al. (2023) Comprehensive phylogenomic time tree of bryophytes reveals deep relationships and uncovers gene incongruences in the last 500 million years of diversification. American Journal of Botany, 110, e16249.

Berry, E.A., Tran, M.L., Dimos, C.S., Budziszek, Jr. M.J., Scavuzzo-Duggan, T.R., Roberts, A.W. (2016) Immuno and affinity cytochemical analysis of cell wall composition in the moss *Physcomitrella patens*. Frontiers in Plant Science, 7, 248.

Blakeney, A.B., Harris, P.J., Henry, R.J., Stone, B.A. (1983) A simple and rapid preparation of alditol acetates for monosaccharide analysis. Carbohydrate Research, 113, 291–299.

Blumenkrantz, N. & Asboe-Hansen, G. (1973) New method for quantitative determination of uronic acids. Analytical Biochemistry, 34, 484–489.

Bowles, A.M.C., Paps, J., Bechtold, U. (2021) Evolutionary origins of drought tolerance in spermatophytes. Frontiers in Plant Science, 12, article655924.

Bowman, J.L., Kohchi, T., Yamato, K.T., et al. (2017). Insights into land plant evolution garnered from the *Marchantia polymorpha* genome. Cell, 171, 287–304.

Carafa, A., Duckett, J.G., Knox, J.P., Ligrone, R. (2005) Distribution of cell-wall xylans in bryophytes and tracheophytes: new insights into basal interrelationships of land plants. New Phytologist, 168, 231–240.

Clarke, A.E., Gleeson, P.A., Jermyn, M.A., Knox, R.B. (1978) Characterization and localization of β-lectins in lower and higher plants. Australian Journal of Plant Physiology, 5, 707–722.

Clarke, A.E., Anderson, R.L., Stone, B.A. (1979) Form and function of arabinogalactans and arabinogalactan-proteins. Phytochemistry, 18, 521 –540.

Classen, B., Witthohn, K., Blascheck, W. (2000) Characterization of an arabinogalactan-protein isolated from pressed juice of *Echinacea purpurea* by precipitation with the β-glucosyl Yariv reagent. Carbohydrate Research, 327, 497–504.

Classen, B., Csávás, M., Borbás, A., Dingermann, T., Zündorf, I. (2004) Monoclonal antibodies against an arabinogalactan-protein from pressed juice of *Echinacea purpurea*. Planta Medica, 70, 861–865.

Classen, B., Mau, S.-L. & Bacic, A. (2005) The arabinogalactan-proteins from pressed juice of *Echinacea purpurea* belong to the hybrid class of hydroxyproline-rich glycoproteins. Planta Medica, 71, 59–66.

Classen, B. (2007) Characterization of an arabinogalactan-protein from suspension culture of *Echinacea purpurea*. Plant Cell Tissue and Organ Culture, 88: 267 – 275.

Cove, D. (2001) The moss, *Physcomitrella patents*. Journal of Plant Growth Regulation, 19, 275–283.

Cui, S., Hu, J., Guo, S., Wang, J., Cheng, Y., Dang, X., Wu, L., He, Y. (2012) Proteome analysis of *Physcomitrella patens* exposed to progressive dehydration and rehydration. Journal of Experimental Botany, 63, 711–726.

Degola, F., Di Toppi, L.S., Petraglia, A. (2022) Bryophytes: how to conquer an alien planet and live happily (ever after). Journal of Experimental Botany, 73 (13), 4267–4272.

de Vries, J. & Archibald, J.M. (2018) Plant evolution: Landmarks on the path to terrestrial life. New Phytologist, 217, 1428–1434.

Dilokpimol, A. & Geshi, N. (2014) *Arabidopsis thaliana* glucuronosyltransferase in family GT14. Plant Signaling & Behavior. 9, e28891.

Dolan, L., Linstead, P., Roberts, K. (1995) An AGP epitope distinguishes a central metaxylem initial from other vascular initials in the *Arabidopsis* root. Protoplasma, 189, 149–155.

Domozych, D.S. & Bagdan, K. (2022) The cell biology of charophytes: exploring the past and models for the future. Plant Physiology, 190, 1588–1608.

Donoghue, P.C.J., Harrison, C.J., Paps, J., Schneider, H. (2021) The evolutionary emergence of land plants. Current Biology, 31, 1281–1298.

Dragićević, M.B, Paunović, D.M., Bogdanović, M.D., Todorović, S.I., & Simonović, A.D. (2020) Ragp: pipeline for mining of plant hydroxyproline-rich glycoproteins with implementation in R. Glycobiology, 30, 19–35.

Edstam, M.M., Viitanen, L., Salminen, T.A., Edqvist, J. (2011) Evolutionary history of the non-specific lipid transfer proteins. Molecular Plant, 4, 947–964.

Edstam, M.M. & Edqvist, J. (2014) Involvement of GPI-anchored lipid transfer proteins in the development of seed coats and pollen in *Arabidopsis thaliana*. Physiologia Plantarum, 152, 32–42.

Eisenhaber, B., Wildpaner, M., Schultz, C.J., Borner, G.H.H., Dupree, P. & Eisenhaber, F. (2003) Glycosylphosphatidylinositol lipid anchoring of plant proteins. Sensitive prediction from sequence- and genome-wide studies for Arabidopsis and Rice. Plant Physiology, 153, 403–419.

Fu, H., Yadav, M.P., Nothnagel, E.A. (2007) *Physcomitrella patens* arabinogalactan proteins contain abundant terminal 3-*O*-methyl-L-rhamnosyl residues not found in angiosperms. Planta, 226, 1511–1524.

Fürst-Jansen, J.M.R., de Vries, S., de Vries, J. (2020) Evo-physio: on stress responses and the earliest land plants. Journal of Experimental Botany, 71, 3254–3269.

Geddes, D.S. & Wilkie, K.C.B. (1971) Hemicelluloses from the stem tissues of the aquatic moss *Fontinalis antipyretica*. Carbohydrate Research, 18, 333–335.

Gille, S., Sharma, V., Baidoo, E.E.K., Keasling, J.D., Scheller, H.V., Pauly, M. (2013) Arabinosylation of a Yariv-precipitable cell wall polymer impacts plant growth as exemplified by the *Arabidopsis* glycosyltransferase mutant ray1. Molecular Plant, 6, 1369–1372.

Gíslason, M.H., Nielsen, H., Almagro Armenteros, J.J., & Johansen, A.R. (2021) Prediction of GPI-anchored proteins with pointer neural networks. Current Research in Biotechnology, 3, 6–13.

Goellner, E.M., Ichinose, H., Kaneko, S., Blaschek, W., Classen, B. (2011) An arabinogalactan-protein from whole grain of *Avena sativa* L. belongs to the wattle-blossom type of arabinogalactan-proteins. Journal of Cereal Science, 53, 244–249.

Goellner, E.M., Gramann, J.C., Classen, B. (2013) Antibodies against Yariv’s Reagent for Immunolocalization of Arabinogalactan-Proteins in Aerial Parts of *Echinacea purpurea*. Planta Medica, 79, 175 – 180.

Happ, K. & Classen, B. (2019) Arabinogalactan-proteins from the liverwort *Marchantia polymorpha* L., a member of a basal land plant lineage, are structurally different to those of angiosperm. Plants, 8, 460.

Harholt, J., Moestrup, Ø., Ulvskov, P. (2016) Why plants were terrestrial from the beginning. Trends in Plant Science, 21, 96–101.

Harris, P.J., Henry, R.J., Blakeney, A.B., Stone, B.A. (1984) An improved procedure for the methylation analysis of oligosaccharides and polysaccharides. Carbohydrate Research, 127, 59–73.

Herman, E.M. & Lamb C.J. (1992) Arabinogalactan-rich glycoproteins are localized on the cell surface and in intravacuolar multivesicular bodies. Plant Physiology, 98, 264–272

Hodgetts, N., Cálix, M., Englefield, E. et al. (2019) A miniature world in decline: European Red List of Mosses, Liverworts and Hornworts. Brussels, Belgium: IUCN.

Johnson, K.L., Andrew M. Cassin, A.M., Lonsdale, A., Bacic, A., Doblin, M.S., Schultz, C.J. (2017a) A motif and amino acid bias bioinformatics pipeline to identify hydroxyproline-rich glycoproteins. Plant Physiology, 174, 886–903.

Johnson, K.L., Cassin, A.M., Lonsdale, A., Ka-Shu Wong, G., Soltis, D.E., Miles, N.W. et al. (2017b) Insights into the evolution of hydroxyproline-rich glycoproteins from 1000 plant transcriptomes. Plant Physiology, 174, 904–921.

Johnson, K.L., Jones, B.J., Schultz, C.J., Bacic, A. (2018) Non-enzymic cell wall (glyco)proteins. In: Annual Plant Reviews online, J.A. Roberts (Ed.). doi: 10.1002/9781119312994.apr0070.

Katoh, K. & Standley, D.M. (2013) MAFFT multiple sequence alignment software version 7: improvements in performance and usability. Molecular Biology and Evolution, 30, 772–780.

Kitazawa, K., Tryfona, T., Yoshimi, Y. et al. (2013) β-galactosyl Yariv reagent binds to the β-1,3-galactan of arabinogalactan proteins. Plant Physiology, 161, 1117–1126.

Kjeldahl, J. (1883) Neue Methode zur Bestimmung des Stickstoffs in organischen Körpern. Zeitschrift für analytische Chemie, 22, 366–382.

Knoch, E., Dilokpimol, A., Tryfona, T., Poulsen, C.P., Xiong, G., Harholt, J., Petersen, B.L., Ulvskov, P., Hadi, M.Z., Kotake, T., Tsumuraya, Y., Pauly, M., Dupree, P., Geshi, N. (2013) A β-glucuronosyltransferase from *Arabidopsis thaliana* involved in biosynthesis of type II arabinogalactan has a role in cell elongation during seedling growth. The Plant Journal, 76, 1016–1029.

Knox, J.P., Linstead, P.J., Cooper, J.P.C., Roberts, K. (1991) Developmentally regulated epitopes of cell surface arabinogalactan proteins and their relation to root tissue pattern formation. The Plant Journal, 1, 317–326.

Kobayashi, Y., Motose, H., Iwamoto, K., & Fukuda, H. (2011) Expression and genome-wide analysis of the xylogen-type gene family. Plant Cell Physiology, 52, 1095–1106.

Konrad, S.S.A. & Ott, T. (2015) Molecular principles of membrane microdomain targeting in plants. Trends in Plant Science, 20, 351–361.

Kremer, C., Pettolino, F., Bacic, A., Drinnan, A. (2004) Distribution of cell wall components in *Sphagnum* hyaline cells and in liverwort and hornwort elaters. Planta, 219, 1023–1035.

Lamesch, P., Berardini, T.Z., Li, D., Swarbreck, D., Wilks, C., Sasidharan, R., et al. (2011). The Arabidopsis Information Resource (TAIR): improved gene annotation and new tools. Nucleic Acids Research, 40, D1202–D1210.

Lang, D., Ullrich, K.K., Murat, F., Fuchs, J., Jenkins, J., Haas, F.B., et al. (2018) The *Physcomitrella patens* chromosome[scale assembly reveals moss genome structure and evolution. The Plant Journal, 93, 515–533.

Lee, K. J.D., Sakata, Y., Mau, S.L., Pettolino, F., Bacic, A., Quatrano, R.S., Knight, C.D., Knox, J.P. (2005) Arabinogalactan proteins are required for apical cell extension in the moss *Physcomitrella patens*. The Plant Cell, 17, 3051–3065.

Letunic, I. & Bork, P. (2021) Interactive tree of life (iTOL) v5: an online tool for phylogenetic tree display and annotation. Nucleic Acids Research, 49, W293–W296.

Li, F.W., Brouwer, P., Carretero-Paulet, L., Cheng, S., de Vries, J., Delaux, P.-M., et al. (2018) Fern genomes elucidate land plant evolution and cyanobacterial symbioses. Nature Plants, 4, 460–472.

Li, J., Nishiyama T., Waller, M. et al. (2020) *Anthoceros* genomes illuminate the origin of land plants and the unique biology of hornworts. Nature Plants, 6, 259–272.

Liu, F., Zhang, X., Lu, C., Zeng, X., Li, Y., Fu, D., Wu, G. (2015) Non-specific lipid transfer proteins in plants: presenting new advances and an integrated functional analysis. Journal of Experimental Botany, 66, 5663–5681.

Liu, Y., Wang, S., Li, L., Yang, T., Dong, S., Wei, T. et al. (2022) The *Cycas* genome and the early evolution of seed plants. Nature Plants, 8, 389–401.

Ligrone, R., Duckett, J.G., Renzaglia, K.S. (2000) Conducting tissues and phyletic relationships of bryophytes. Philosophical Transactions of the Royal Society B, 355, 795–813.

Ligrone, R., Vaughn, K.C., Renzaglia, K.S., Knox, J.P., Duckett, J.G. (2002) Diversity in the distribution of polysaccharide and glycoprotein epitopes in the cell walls of bryophytes: new evidence for the multiple evolution of water-conducting cells. New Phytologist, 156, 491–508.

LoRicco, J.G., Sun, L., Bauer, L., Sgambettera, G., Epstein, R., Bagdan, K., Winegrad, A., et al. (2024) The multifunctional roles of the extracellular matrix in the sessile life of the zygnematophyte *Penium margaritaceum*: stick, glide and cluster. (Physiologia Plantarum, 176, e14520.

Lu, S., Wang, J., Chitsaz, F., Derbyshire, M.K., Geer, R.C., Gonzales, N.R., et al. (2020) CDD/SPARCLE: the conserved domain database in 2020. Nucleic Acids Research, 48, D265–D268.

Ma, Y., Yan, C., Li, H., Wu, W., Liu, Y., Wang, Y. et al. (2017) Bioinformatics prediction and evolution analysis of arabinogalactan proteins in the plant kingdom. Frontiers in Plant Science, 8, 66.

Ma, Y., Zeng, W., Bacic, A., Johnson, K. (2018) AGPs through time and space. In: Annual Plant Reviews online, J.A. Roberts (Ed.). doi: 10.1002/9781119312994.apr0608

Ma, Y. & Johnson, K. (2023) Arabinogalactan proteins – Multifunctional glycoproteins of the plant cell wall. The Cell Surface, 9, 100102, doi: 10.1016/j.tcsw.2023.100102

Marchant, D.B., Chen, G., Cai, S., Chen, F., Schafran, P., Jenkins, J., et al. (2022) Dynamic genome evolution in a model fern. Nature Plants, 8, 1038–1051.

Mareri, L., Romi, M., Cai, G. (2018) Arabinogalactan proteins: actors or spectators during abiotic and biotic stress in plants? Plant Biosystems, 153, 173–185.

Mansouri, K. (2012) Comparative ultrastructure of apical cells and derivatives in bryophytes, with special reference to plasmodesmata (Ph.D. thesis). Southern Illinois University Carbondale, Ann Arbor, United States

McCourt, R.M., Lewis, L.A., Strother, P.K., Delwiche, C.F., Wickett, N.J., de Vries, J., Bowman, J.L. (2023) Green land: multiple perspectives on green algal evolution and the earliest land plants. American Journal of Botany, 110, e16175.

Morris, J.L., Puttick, M.N., Clark, J.W., Edwards, D., Kenrick, P., Pressel, S., Wellman, C.H., Yang, Z., Schneider, H., Donoghue, P.C.J. (2018) The timescale of early land plant evolution. Proceedings of the National Academy of Sciences, 115, E2274–E2283. doi: 10.1073/pnas.1719588115.

Motose, H., Sugiyama, M., Fukuda, H. (2004) A proteoglycan mediates inductive interaction during plant vascular development. Nature, 429, 873–878.

Mueller, K.-K., Pfeifer, L., Schuldt, L., Szövényi, P., de Vries, S., de Vries, J., Johnson, K.L., Classen, B. (2023) Fern cell walls and the evolution of arabinogalactan proteins in streptophytes. The Plant Journal, 114, 875–894.

Nguyen, L.T., Smidt, H.A., von Haeseler, A. & Minh, B.Q. (2015) IQ-TREE: A fast and effective stochastic algorithm for estimating maximum-likelihood phylogenies. Molecular Biology and Evolution, 32, 268–274.

Nystedt, B., Street, N.R., Wetterbom, A., Zuccolo, A., Lin, Y.-C., Scofield, D.G., et al. (2013). The Norway spruce genome sequence and conifer genome evolution. Nature, 497, 579–584.

Ogawa, K., Yamaura, M., Maruyama, I. (1997) Isolation and identification of 2-*O*-methyl-L-rhamnose and 3-*O*-methyl-L-rhamnose as constituents of an acidic polysaccharide of *Chlorella vulgaris*. Bioscience, Biotechnology, and Biochemistry, 61, 539–540.

Ogawa-Ohnishi, M. & Matsubayashi, Y. (2015) Identification of three potent hydroxyproline *O*-galactosyltransferases in *Arabidopsis*. The Plant Journal, 81, 736–746.

Okawa, R., Hayashi, Y., Yamashita, Y., Matsubayashi, Y., Ogawa-Ohnishi, M. (2023) Arabinogalactan protein polysaccharide chains are required for normal biogenesis of plasmodesmata. The Plant Journal, 113, 493–503

Olmos, E., de la Garma, J. G., Gomez-Jimenez, M. C., Fernandez-Garcia, N. (2017) Arabinogalactan proteins are involved in salt-adaptation and vesicle trafficking in Tobacco BY-2 cell cultures. Frontiers in Plant Science, 8, 1092; doi: 10.3389/fpls.2017.01092

Palacio-López, K., Tinaz, B., Holzinger, A., Domozych, D. S. (2019) Arabinogalactan proteins and the extracellular matrix of charophytes: a sticky business. Frontiers in Plant Science, 10, 447.

Paulsen, B.S., Craik, D. J., Dunstan, D. E., Stone, B. A., Bacic, A. (2014) The Yariv reagent: behaviour in different solvents and interaction with a gum arabic arabinogalactan-protein. Carbohydrate Polymers, 106, 460–468.

Peña, M.J., Darvill, S.G., Eberhard, S., York, W.S., O’Neill, M.A. (2008) Moss and liverwort xyloglucans contain galacturonic acid and are structurally distinct from the xyloglucans synthesized by hornworts and vascular plants. Glycobiology, 18, 891–904.

Petersen, M. (2003) Cinnamic acid 4-hydroxylase from cell cultures of the hornwort *Anthoceros agrestis*. Planta, 217, 96–101

Pfeifer, L., Mueller, K.-K., Classen, B. (2022a) The cell wall of hornworts and liverworts: innovations in early land plant evolution? Journal of Experimental Botany, 73, 4454– 4472.

Pfeifer, L., Utermöhlen, J., Happ, K., Permann, C., Holzinger, A., von Schwartzenberg, K., Classen, B. (2022b) Search for evolutionary roots of land plant arabinogalactan-proteins in charophytes: presence of a rhamnogalactan-protein in *Spirogyra pratensis* (Zygnematophyceae). The Plant Journal, 109, 568–584.

Pierleoni, A., Martelli, P.L. & Casadio, R. (2008) PredGPI: a GPI-anchor predictor. BMC Bioinformatics, 9, 392.

Płachno, B.J., Kapusta, M., Swiatek, P., Stolarczyk, P., Kocki, J. (2020) Immunodetection of Pectic Epitopes, Arabinogalactan Proteins, and Extensins in Mucilage Cells from the Ovules of *Pilosella officinarum* Vaill. and *Taraxacum officinale* Agg. (Asteraceae). International Journal of Molecular Sciences, 21, 9642, doi: 10.3390/ijms21249642

Popper, Z.A. and Fry, S.C. (2003) Primary cell wall composition of bryophytes and charophytes. Annals of Botany, 91, 1–12.

Price, M.N., Dehal, P.S., & Arkin, A.P. (2010) FastTree 2 – Approximately-maximum likelihood trees for large alignments. PLOS One, 5, e9490.

Puttick, M.N., Morris, J.L., Williams, T.A., Cox, C.J., Edwards, D., Kenrick, P. et al. (2018) The interrelationship of land plants and the nature of the ancestral embryophyte. Current Biology, 28, 733–745.

Rensing, S.A., Lang, D., Zimmer, A.D. et al. (2008) The *Physcomitrella* genome reveals evolutionary insights into the conquest of land by plants. Science, 319, 64–69.

Reynolds, E. S. (1963) The use of lead citrate at high pH as an electron-opaque stain in electron microscopy. The Journal of Cell Biology, 17, 208–212.

Roberts A.W., Lahnstein, J., Hsieh, Y.S.Y., et al. (2018) Functional characterization of a glycosyltransferase from the moss *Physcomitrella patens* involved in the biosynthesis of a novel cell wall arabinoglucan. The Plant Cell, 30, 1293–1308.

Ruprecht, C., Bartetzko, M.P., Senf, D., et al. (2017) A synthetic glycan microarray enables epitope mapping of plant cell wall glycan-directed antibodies. Plant Physiology, 175, 1094–1104.

Sala, K., Malarz, K., Barlow, P.W., Kurczyńska, E.U. (2017) Distribution of some pectic and arabinogalactan protein epitopes during Solanum lycopersicum (L.) adventitious root development. BMC Plant Biology, 17, 25; doi: 10.1186/s12870-016-0949-3

Šamaj, J., Šamajová, O., Peters, M., Baluška, F., Lichtscheidl, I., Knox, J.P., Volkmann, D. (2000) Immunolocalization of LM2 arabinogalactan protein epitope associated with endomembranes of plant cells. Protoplasma, 212, 186–196

Schafran, P. Hauser, D.A., Nelson, J.M., Xu, X., Mueller, L.A., Kulshrestha, S., Smalley, I., et al. (2025) Pan-phylum genomes of hornworts reveal conserved autosomes but dynamic accessory and sex chromosomes. Nature Plants, 11, 49–62.

Seifert, G.J. (2021) The FLA4-FEI pathway: a unique and mysterious signaling module related to cell wall structure and stress signaling. Genes, 12, 145.

Seifert, G.J. & Roberts, K. (2007) The biology of arabinogalactan proteins. Annual Review of Plant Biology, 58, 137–161.

Showalter, A.M. & Basu, D. (2016) Glycosylation of arabinogalactan-proteins essential for development in *Arabidopsis*. Communicative and Integrative Biology, 5, e1177687.

Shibaya, T., & Sugawara, Y. (2009) Induction of multinucleation by β-glucosyl Yariv reagent in regenerated cells from Marchantia polymorpha protoplasts and involvement of arabinogalactan proteins in cell plate formation. Planta, 2030, 581–588.

Silva, J., Ferraz, R., Dupree, P., Showalter, A.M., Coimbra, S. (2020) Three Decades of Advances in Arabinogalactan-Protein Biosynthesis. Frontiers in Plant Science, 11, 610377, doi: 10.3389/fpls.2020.610377

Smallwood, M., Yates, E.A., Willats, W.G.T., Martin, H., Knox, J. P. (1996) Immunochemical comparison of membrane-associated and secreted arabinogalactan-proteins in rice and carrot. Planta, 198, 452–459.

Stegemann, H. & Stalder, K. (1967) Determination of hydroxyproline. Clinica Chimica Acta, 18, 267–273.

Strasser, R., Seifert, G., Doblin, M.S., Johnson, K.L., Ruprecht, C., Pfrengle, F., Bacic, A., Estevez, J.M. (2021) Cracking the “sugar code”: a snapshot of *N*- and *O*-glycosylation pathways and functions in plants cells. Frontiers in Plant Science, 12, 640919, 10.3389/fpls.2021.640919

Su, D., Yang, L., Shi, X., Ma, X., Zhou, X., Hedges, S.B., Zhong, B. (2021) Large-scale phylogenomic analyses reveal the monophyly of bryophytes and neoproterozoic origin of land plants. Molecular Biology and Evolution, 38, 3332–3344.

Szövényi, P., Frangedakis, E., Ricca, M., Quandt, D., Wicke, S., Langdale, J.A. (2015) Establishment of *Anthoceros agrestis* as a model species for studying the biology of hornworts. BMC Plant Biology, 15, 98; doi: 10.1186/s12870-015-0481-x

Taylor, R.L. & Conrad, H.E. (1972) Stoichiometric depolymerization of polyuronides and glycosaminoglycuronans to monosaccharides following reduction of their carbodiimide-activated carboxyl groups. Biochemistry 11, 1383–1388.

Temple, H., Mortimer, J.C., Tryfona, T., Yu, X., Lopez-Hernandez, F., Sorieul, M., Anders, N., Dupree, P. (2019) Two members of the DUF579 family are responsible for arabinogalactan methylation in Arabidopsis. Plant Direct, 3, e00117.

Thude, S. & Classen, B. (2005) High molecular weight constituents from roots of *Echinacea pallida*: An arabinogalactan-protein and an arabinan. Phytochemistry, 66, 1026–1032.

Tryfona, T., Liang, H.-C., Kotake, T., Tsumuraya, Y., Stephens, E., Dupree, P. (2012) Structural characterization of *Arabidopsis* leaf arabinogalactan polysaccharides. Plant Physiology, 160, 653–666.

Wang, Q.-H., Zhang, J., Liu, Y., Jia, Y., Jiao, Y.-N., Xu, B., Chen, Z.-D. (2022) Diversity, phylogeny, and adaptation of bryophytes: insights from genomic and transcriptomic data. Journal of Experimental Botany, 73, 4306–4322.

Waterhouse, A.M., Procter, J.B., Martin, D.M.A., Clamp, M. & Barton, G.J. (2009) Jalview Version 2 – a multiple sequence alignment editor and analysis workbench. Bioinformatics, 25, 1189–1191.

Verhertbruggen, Y., Marcus, S.E., Haeger, A., Verhoef, R., Schols, H.A., McCleary, B.V. McKee, L., Gilbert, H. J., Knox, J. P. (2009) Developmental complexity of arabinan polysaccharides and their processing in plant cell walls. The Plant Journal, 59, 413–425.

Wack, M., Classen, B., Blaschek, W. (2005) An acidic arabinogalactan-protein from the roots of *Baptisia tinctoria* (L.) R. Brown. Planta Medica, 71, 814–818.

Wegner, L. & Ehlers, K. (2024) Plasmodesmata dynamics in bryophyte model organisms: secondary formation and developmental modifications of structure and function. Planta, 260, 45, doi: 10.1007/s00425-024-04476-1

Wickell, D., Kuo, L.Y., Yang, H.P., Ashok, A.D., Irisarri, I., Dadras, A. (2021) Underwater CAM photosynthesis elucidated by *Isoetes* genome. Nature Communications, 12, 6348.

Woudenberg, S., Renema, J., Tomescu, A.M.F., De Rybel, B., Weijers, D. (2022) Deep origin and gradual evolution of transporting tissues: perspectives from across the land plants. *Plant Physiology*, kiac304, doi: 10.1093/plphys/kiac304.

Yadav, S., Basu, S., Srivastava, A., Biswas, S., Mondal, R., Jha, V.K., Singh, S.K., Mishra, Y. (2023) Bryophytes as modern model plants: an overview of their development, contributions, and future prospects. Journal of Plant Growth Regulation, 42, 6933–6950.

Yariv, J., Rapport, M.M., Graf, L. (1962) The interaction of glycosides and saccharides with antibody to the corresponding phenylazo glycosides. Biochemical Journal, 85, 383– 388.

Yates, E.A., Valdor, J.F., Haslam, S.M., Morris, H.R., Dell, A., Mackie, W., Knox, J.P. (1996) Characterization of carbohydrate structural features recognized by anti-arabinogalactan-protein monoclonal antibodies. Glycobiology, 6, 131–139.

Ye Z.-H. & Zhong, R. (2022) Cell wall biology of the moss *Physcomitrium patens*. Journal of Experimental Botany, 73, 4440–4453

Zhang, H., Yohe, T., Huang, L., Entwistle, S., Wu, P., Yang, Z. et al. (2018) dbCAN2: a meta server for automated carbohydrate-active enzyme annotation. Nucleic Acids Research, 46, W95–W101.

Zhang, J., Fu, X.-X., Li, R.-Q., et al. (2020) The hornwort genome and early land plant evolution. Nature Plants, 6, 107–118.

