## Supplementary Material for "New insights into bryophyte arabinogalactan-proteins from a hornwort and a moss model organism"

**Table S1** Neutral monosaccharide composition of high molecular weight, water-soluble fractions (AE) from *Anthoceros agrestis* (plants and cell cultures) and *Physcomitrium patens* in % (mol $mol^{-1}$; n=3; tr: trace value < 1 %).

| Neutral monosaccharide | *A. agrestis* plants | *A. agrestis* cell culture | *P. patens* plants |
|---|---|---|---|
| 3-*O*-MeRha | 1.3 ± 0.1 | 2.8 ± 0.0 | 2.7 ± 0.0 |
| Rha | 2.2 ± 0.1 | 1.7 ± 0.5 | 4.2 ± 0.0 |
| Fuc | 1.4 ± 0.1 | 5.2 ± 0.3 | 4.0 ± 0.1 |
| Rib | tr | 9.4 ± 0.8 | 2.1 ± 0.2 |
| Ara | 8.4 ± 0.1 | 28.5 ± 2.6 | 11.5 ± 0.2 |
| Xyl | 23.8 ± 0.1 | 11.1 ± 0.8 | 7.7 ± 0.1 |
| Man | 1.2 ± 0.0 | 7.1 ± 1.0 | 6.0 ± 0.2 |
| Gal | 42.2 ± 0.3 | 26.0 ± 0.5 | 14.2 ± 0.3 |
| Glc | 19.5 ± 0.2 | 8.2 ± 3.2 | 47.6 ± 0.8 |

**Table S2** Content of uronic acids in high molecular weight, water-soluble fractions (AE) and AGPs from *Anthoceros agrestis* plants and cell cultures and *Physcomitrium patens*, in % (w $w^{-1}$).

| | *A. agrestis* plants | *A. agrestis* cell culture | *P. patens* plants |
|---|---|---|---|
| AE | 11.1 | 2.2 | 3.1 |
| AGP | 7.3 | 4.8 | 6.5 |

**Table S3** Antibodies directed against AGP glycan motifs used in this study.

| Antibody | Epitope | Key References |
|---|---|---|
| JIM13 | AGP glycan, e.g. β-D-Glc*p*A-(1→3)-α-D-Gal*p*A-(1→2)-α-L-Rha | Pfeifer *et al.* (2022b); Yates *et al.* (1996) |
| KM1 | (1→6)-β-D-Gal*p* units in AGs type II | Classen *et al.* (2004); Ruprecht *et al.* (2017) |
| LM2 | (1→6)-β-D-Gal*p* units with terminal ß-D-Glc*p*A in AGPs | Ruprecht *et al.* (2017); Smallwood *et al.* (1996); |
| LM6 | (1→5)-α-L-Ara*f* oligomers in arabinan or AGPs | Verhertbruggen *et al.* (2009) |

**Table S4** Homolog numbers of characterized AGP-active enzymes as identified by phylogenetic genome analysis of 12 hornwort and 2 setaphyte genomes.

| **species** | **P4H1** | **P4H5** | **P4H13** | **GALT29A** | **GALT31A/KNS4 /UPEX1** | **GALT2-6** | **HPGT1-3** | **GALT9** | **GlcAT14A-E** | **FUTs (GT37)** | **RAY1 (GT77)** | **AGM1+2** | **Rs*Araf*** | **AGAL2-3** | **AtBGAL8** | **GH43** | **AtGUS2** |
|---|---|---|---|---|---|---|---|---|---|---|---|---|---|---|---|---|---|
| *Leicosporoceros dussii* | 1 | 1 | 1 | 2 | 1 | 3 | 1 | 2 | 1 | 5 | 1 | 0 | 2 | 1 | 1 | 1 | 1 |
| *Anthoceros agrestis 'Bonn'* | 3 | 1 | 1 | 1 | 1 | 4 | 1 | 1 | 3 | 3 | 1 | 0 | 3 | 1 | 1 | 0 | 1 |
| *Anthoceros agrestis 'Oxford'* | 3 | 1 | 1 | 2 | 1 | 4 | 1 | 2 | 3 | 4 | 1 | 0 | 3 | 1 | 2 | 1 | 1 |
| *Anthoceros punctatus* | 1 | 1 | 1 | 2 | 1 | 3 | 1 | 2 | 3 | 4 | 1 | 0 | 3 | 1 | 2 | 1 | 1 |
| *Anthoceros fusiformis* | 1 | 1 | 1 | 2 | 1 | 2 | 1 | 2 | 3 | 4 | 1 | 0 | 3 | 1 | 2 | 2 | 1 |
| *Notothylas orbicularis* | 1 | 1 | 1 | 3 | 1 | 2 | 1 | 2 | 1 | 3 | 1 | 0 | 2 | 1 | 1 | 1 | 1 |
| *Paraphymatoceros pearsonii* | 1 | 1 | 2 | 3 | 1 | 3 | 1 | 2 | 1 | 3 | 1 | 0 | 3 | 1 | 2 | 1 | 1 |
| *Phaeoceros carolinianus* | 1 | 1 | 1 | 3 | 1 | 3 | 1 | 4 | 1 | 3 | 1 | 0 | 5 | 2 | 2 | 1 | 1 |
| *Phaeoceros sp.* | 1 | 1 | 1 | 4 | 1 | 3 | 1 | 2 | 1 | 3 | 1 | 0 | 6 | 3 | 2 | 1 | 1 |
| *Megaceros flagellaris* | 1 | 1 | 1 | 4 | 1 | 2 | 1 | 2 | 1 | 4 | 1 | 0 | 3 | 3 | 2 | 1 | 1 |
| *Phaeomegaceros chiloensis* | 1 | 1 | 1 | 3 | 1 | 2 | 1 | 2 | 1 | 4 | 1 | 0 | 5 | 3 | 0 | 1 | 1 |
| *Phymatoceros phymatodes* | 1 | 1 | 1 | 3 | 1 | 2 | 1 | 2 | 1 | 3 | 1 | 0 | 5 | 3 | 1 | 1 | 1 |
| *Marchantia polymorpha* | 0 | 1 | 1 | 4 | 2 | 2 | 1 | 1 | 1 | 3 | 1 | 0 | 5 | 1 | 3 | 1 | 3 |
| *Physcomitrium patens* | 2 | 2 | 2 | 2 | 4 | 3 | 2 | 2 | 7 | 6 | 1 | 0 | 2 | 1 | 4 | 2 | 4 |
| **average hornworts** | **1.3** | **1.0** | **1.1** | **2.7** | **1.0** | **2.8** | **1.0** | **2.1** | **1.7** | **3.6** | **1.0** | **0.0** | **3.6** | **1.8** | **1.5** | **1.0** | **1.0** |

**Table S5** Numbers of identified protein sequences for classical (+/- GPI-anchor) and hybrid arabinogalactan-proteins. We used the motif and amino acid bias (MAAB) classification system of Johnson *et al*., 2017.

| species | GPI-AGP (class 1) | CL-EXT (class 2) | PRP (class 3) | Non-GPI-AGP (class 4) | Hybrid AGP (class 5-8) | GPI-EXT (class 9) | Hybrid EXT (class 10-14) | Hybrid PRP (class 15-18) | Shared bias (class 19-23) | Non-HRGPs (class 24) |
|---|---|---|---|---|---|---|---|---|---|---|
| *Leicosporoceros dussii* | 1.0 | 2.0 | - | - | - | - | 1.0 | 2.0 | - | 2.0 |
| *Anthoceros agrestis 'Bonn'* | - | - | - | 2.0 | - | - | - | - | - | 4.0 |
| *Anthoceros agrestis 'Oxford'* | - | 1.0 | - | 2.0 | - | - | 1.0 | - | - | 6.0 |
| *Anthoceros punctatus* | 1.0 | 1.0 | - | 1.0 | - | - | 1.0 | - | - | 5.0 |
| *Anthoceros fusiformis* | - | - | - | 1.0 | - | - | 1.0 | 1.0 | - | 3.0 |
| *Notothylas orbicularis* | - | 2.0 | - | 3.0 | 1.0 | - | - | - | 3.0 | 4.0 |
| *Paraphymatoceros pearsonii* | - | 1.0 | - | 4.0 | - | - | - | - | 1.0 | 1.0 |
| *Phaeoceros carolinianus* | - | 2.0 | - | 6.0 | 1.0 | - | 2.0 | - | 4.0 | 6.0 |
| *Phaeoceros sp.* | - | 4.0 | - | 7.0 | 2.0 | - | 2.0 | 1.0 | 3.0 | 8.0 |
| *Megaceros flagellaris* | - | 3.0 | - | 4.0 | - | - | - | 1.0 | 2.0 | - |
| *Phaeomegaceros chiloensis* | - | 3.0 | - | 2.0 | - | - | 1.0 | - | 1.0 | 1.0 |
| *Phymatoceros phymatodes* | - | 5.0 | - | 3.0 | - | - | 1.0 | 2.0 | - | 5.0 |
| *Marchantia polymorpha* | 20.0 | 1.0 | - | 9.0 | 1.0 | - | 3.0 | 1.0 | 2.0 | 13.0 |
| *Physcomitrium patens* | 16.0 | - | - | 18.0 | 2.0 | - | - | - | 3.0 | 1.0 |
| **average hornworts** | **0.2** | **2.0** | **-** | **2.9** | **0.3** | **-** | **0.8** | **0.6** | **1.2** | **3.8** |

**Table S6** Numbers of identified protein sequences for chimeric arabinogalactan-proteins. We focused here on fasciclin-like, plastocyanin-like and xylogen-like AGPs.

| species | fasciclin-like AGPs (pfam02469) | plastocyanin-like AGPs (pfam02298) | xylogen-like AGPs (pfam14368) |
|---|---|---|---|
| *Leicosporoceros dussii* | 3 | 1 | 0 |
| *Anthoceros agrestis 'Bonn'* | 4 | 19 | 2 |
| *Anthoceros agrestis 'Oxford'* | 4 | 21 | 2 |
| *Anthoceros punctatus* | 2 | 12 | 2 |
| *Anthoceros fusiformis* | 2 | 5 | 2 |
| *Notothylas orbicularis* | 6 | 7 | 1 |
| *Paraphymatoceros pearsonii* | 3 | 9 | 1 |
| *Phaeoceros carolinianus* | 1 | 10 | 1 |
| *Phaeoceros sp.* | 3 | 23 | 1 |
| *Megaceros flagellaris* | 4 | 7 | 1 |
| *Phaeomegaceros chiloensis* | 2 | 12 | 1 |
| *Phymatoceros phymatodes* | 7 | 8 | 1 |
| *Marchantia polymorpha* | 9 | 21 | 3 |
| *Physcomitrium patens* | 5 | 13 | 6 |
| **average hornworts** | **3.4** | **11.2** | **1.3** |

**Table S7** Detailed analysis of sequence characteristics in non-specific lipid transfer protein domains within xylogen-like AGPs of bryophytes. 12 hornwort genomes and two setaphyte genomes (*Physcomitrium patens*, Pp; *Marchantia polymorpha*, Mp) were searched. The column "type" uses the classification system of Edstam *et al*. (2011). For the two setaphytes some sequences contained two predicted domains which are here mentioned as "$1^{st}$ domain" and "$2^{nd}$ domain".

| id | spacing patterns | | | | | | | | | | | GPI | type |
|---|---|---|---|---|---|---|---|---|---|---|---|---|---|
| AagrBONN_evm.model.Sc2ySwM_344.1345.2 | C | 9 | C | 14 | CC | 12 | C-1-C | 21 | C | 11 | C | - | D/G |
| AagrBONN_evm.model.Sc2ySwM_368.1844.1 | C | 9 | C | 14 | CC | 12 | C-1-C | 22 | C | 8 | C | ✓ | G |
| AnagrOXF.S3G187000.t1 | C | 9 | C | 14 | CC | 12 | C-1-C | 22 | C | 8 | C | ✓ | G |
| AnagrOXF.S5G197600.t1 | C | 9 | C | 14 | CC | 12 | C-1-C | 21 | C | 11 | C | ✓ | G |
| Anfus.S1G112900.t1 | C | 9 | C | 14 | CC | 12 | C-1-C | 21 | C | 9 | C | - | D/G |
| Anfus.S3G155900.t1 | C | 9 | C | 14 | CC | 12 | C-1-C | 22 | C | 8 | C | ✓ | G |
| Anpun.S1G391000.t1 | C | 9 | C | 14 | CC | 12 | C-1-C | 22 | C | 8 | C | ✓ | G |
| Anpun.S3G522700.t1 | C | 9 | C | 14 | CC | 12 | C-1-C | 21 | C | 11 | C | ✓ | G |
| Mefla.S3G388200.t1 | C | 9 | C | 14 | CC | 12 | C-1-C | 24 | C | 9 | C | ✓ | G |
| Noorb.S3G288500.t1 | C | 9 | C | 14 | CC | 12 | C-1-C | 25 | C | 9 | C | ✓ | G |
| Phcar.S4G164600.t1 | C | 9 | C | 14 | CC | 12 | C-1-C | 25 | C | 9 | C | ✓ | G |
| Phchi.S1G653500.t1 | C | 9 | C | 14 | CC | 12 | C-1-C | 22 | C | 8 | C | ✓ | G |
| Papea.S4G078400.t1 | C | 9 | C | 14 | CC | 12 | C-1-C | 22 | C | 9 | C | - | D/G |
| Phphy.S3G353100.t1 | C | 9 | C | 14 | CC | 12 | C-1-C | 27 | C | 9 | C | ✓ | G |
| Phsp.C5G108400.t1 | C | 9 | C | 14 | CC | 12 | C-1-C | 25 | C | 9 | C | ✓ | G |
| Mp1g21600.1 | C | 9 | C | 14 | CC | 12 | C-1-C | 29 | C | 12 | C | ✓ | G |
| Mp1g22240.1 | C | 9 | C | 14 | CC | 12 | C-1-C | 24 | C | 8 | C | ✓ | G |
| Mp8g18180.1 / $1^{st}$ domain | C | 9 | C | 14 | CC | 12 | C-1-C | 27 | C | 8 | C | ✓ | G |
| Mp8g18180.1 / $2^{nd}$ domain | C | 9 | C | 14 | CC | 12 | C-1-C | 21 | C | 8 | C | ✓ | G |
| Pp3c11_11470V3.1.p | C | 9 | C | 16 | CC | 12 | C-1-C | 23 | C | 8 | C | ✓ | G |
| Pp3c11_8360V3.1.p | C | 9 | C | 14 | CC | 12 | C-1-C | 23 | C | 8 | C | ✓ | G |
| Pp3c14_3790V3.1.p | C | 9 | C | 14 | CC | 12 | C-1-C | 27 | C | 8 | C | ✓ | G |
| Pp3c1_31020V3.1.p / $1^{st}$ domain | C | 9 | C | 14 | CC | 12 | C-1-C | 27 | C | 8 | C | ✓ | G |
| Pp3c1_31020V3.1.p / $2^{nd}$ domain | C | 9 | C | 15 | CC | 12 | C-1-C | 25 | C | 8 | C | ✓ | G |
| Pp3c2_6280V3.1.p / $1^{st}$ domain | C | 9 | C | 15 | CC | 12 | C-1-C | 25 | C | 8 | C | ✓ | G |
| Pp3c2_6280V3.1.p / $2^{nd}$ domain | C | 9 | C | 14 | CC | 12 | C-1-C | 27 | C | 8 | C | ✓ | G |
| Pp3c7_20230V3.1.p | C | 9 | C | 14 | CC | 12 | C-1-C | 23 | C | 8 | C | ✓ | G |

**Table S8** Detailed analysis of sequence characteristics in non-specific lipid transfer protein domains within xylogen-like AGPs of selected other embryophytes. Two angiosperms (*Amborella trichopoda*, AMTR; *Arabidopsis thaliana*, AT), two gymnosperms (*Picea abies*, MA, *Cycas panzhihuaensis*, CYCAS), two lycophytes (*Isoetes taiwaniensis*, Itaiw; *Selaginella moellendorfii*, Smo) and three ferns (*Ceratopteris richardii*, Ceric; *Azolla filiculoides*, Azfi; *Salvinia cucullata*, Sacu) were analyzed. The column "type" uses the classification system of Edstam *et al*. (2011). Some sequences contained two predicted domains which are here mentioned as "1st domain" and "2nd domain".

| id | spacing patterns | | | | | | | | | | | GPI | type |
|---|---|---|---|---|---|---|---|---|---|---|---|---|---|
| AMTR_s00002p00272100 | C | 9 | C | 20 | CC | 12 | C-1-C | 24 | C | 6 | C | ✓ | G |
| AMTR_s00002p00272110 | C | 10 | C | 17 | CC | 12 | C-1-C | 24 | C | 8 | C | ✓ | G |
| AMTR_s00002p00272120 | C | 9 | C | 14 | CC | 12 | C-1-C | 24 | C | 9 | C | ✓ | G |
| AMTR_s00010p00204620 | C | 9 | C | 16 | CC | 12 | C-1-C | 24 | C | 9 | C | ✓ | G |
| AT1G03103 | C | 9 | C | 14 | CC | 12 | C-1-C | 26 | C | 9 | C | ✓ | G |
| AT1G05450 | C | 10 | C | 14 | CC | 12 | C-1-C | 24 | C | 8 | C | ✓ | G |
| AT1G18280 | C | 6 | C | 13 | CC | 12 | C-1-C | 25 | C | 8 | C | ✓ | G |
| AT1G27950 | C | 9 | C | 14 | CC | 12 | C-1-C | 29 | C | 9 | C | ✓ | G |
| AT1G32280 | C | 10 | C | 17 | CC | 9 | C-1-C | 22 | C | 9 | C | - | G/D |
| AT1G36150 (AtXYLP5) | C | 9 | C | 16 | CC | 12 | C-1-C | 24 | C | 7 | C | ✓ | G |
| AT1G73560 | C | 6 | C | 14 | CC | 12 | C-1-C | 25 | C | 8 | C | ✓ | G |
| AT1G73890 | C | 9 | C | 14 | CC | 12 | C-1-C | 26 | C | 8 | C | ✓ | G |
| AT2G13820 (AtXYP2) | C | 9 | C | 16 | CC | 12 | C-1-C | 24 | C | 9 | C | ✓ | G |
| AT2G44290 (AtXYLP9) | C | 9 | C | 14 | CC | 12 | C-1-C | 26 | C | 8 | C | ✓ | G |
| AT2G48130 (AtXYLP11) | C | 9 | C | 14 | CC | 12 | C-1-C | 25 | C | 9 | C | ✓ | G |
| AT2G48140 | C | 10 | C | 17 | CC | 12 | C-1-C | 25 | C | 8 | C | ✓ | G |
| AT3G22600 (AtXYLP12) | C | 9 | C | 14 | CC | 12 | C-1-C | 25 | C | 9 | C | ✓ | G |
| AT3G22620 | C | 10 | C | 17 | CC | 12 | C-1-C | 24 | C | 8 | C | ✓ | G |
| AT3G43720 (AtXYLP10) | C | 9 | C | 18 | CC | 12 | C-1-C | 26 | C | 9 | C | ✓ | G |
| AT4G08670 (AtXYLP3) | C | 9 | C | 16 | CC | 12 | C-1-C | 24 | C | 8 | C | ✓ | G |
| AT4G14805 | C | 9 | C | 17 | CC | 13 | C-1-C | 24 | C | 12 | C | ✓ | G |
| AT5G09370 (AtXYLP4) | C | 9 | C | 16 | CC | 12 | C-1-C | 23 | C | 9 | C | ✓ | G |
| AT5G48490 | C | 9 | C | 15 | CC | 9 | C-1-C | 24 | C | 7 | C | - | G/D |
| AT5G64080 (AtXYP1) | C | 9 | C | 16 | CC | 12 | C-1-C | 24 | C | 9 | C | ✓ | G |
| Azfi_s0011.g012661 | C | 9 | C | 14 | CC | 12 | C-1-C | 28 | C | 8 | C | ✓ | G |
| Azfi_s0803.g087563 | C | 9 | C | 15 | CC | 12 | C-1-C | 26 | C | 9 | C | ✓ | G |
| Azfi_s0803.g087563 | C | 9 | C | 15 | CC | 12 | C-1-C | 26 | C | 9 | C | ✓ | G |
| Azfi_s0803.g087563 | C | 9 | C | 15 | CC | 12 | C-1-C | 26 | C | 9 | C | ✓ | G |
| Ceric.01G082200.1.p | C | 9 | C | 14 | CC | 12 | C-1-C | 27 | C | 8 | C | ✓ | G |
| Ceric.04G032800.1.p | C | 9 | C | 15 | CC | 14 | C-1-C | 22 | C | 10 | C | ✓ | G |
| Ceric.05G095500.1.p / 1st domain | C | 9 | C | 14 | CC | 12 | C-1-C | 24 | C | 9 | C | ✓ | G |
| Ceric.05G095500.1.p / 2nd domain | C | 9 | C | 14 | CC | 12 | C-1-C | 24 | C | 9 | C | ✓ | G |
| Ceric.07G064700.1.p | C | 10 | C | 17 | CC | 9 | C-1-C | 22 | C | 7 | C | - | G/D |
| Ceric.12G092700.1.p | C | 9 | C | 15 | CC | 14 | C-1-C | 24 | C | 10 | C | ✓ | G |
| Ceric.14G098300.1.p | C | 9 | C | 15 | CC | 14 | C-1-C | 24 | C | 10 | C | ✓ | G |
| Ceric.32G069900.1.p | C | 9 | C | 15 | CC | 14 | C-1-C | 24 | C | 9 | C | ✓ | G |
| Ceric.35G030300.1.p | C | 12 | C | 14 | CC | 12 | C-1-C | 24 | C | 9 | C | ✓ | G |
| CYCAS_013046 | C | 9 | C | 14 | CC | 18 | C-1-C | 21 | C | 10 | C | - | G/D |
| CYCAS_024350 | C | 9 | C | 11 | CC | 12 | C-1-C | 24 | C | 9 | C | - | G/D |
| CYCAS_024351 | C | 9 | C | 16 | CC | 12 | C-1-C | 24 | C | 9 | C | ✓ | G |
| CYCAS_024351 | C | 9 | C | 16 | CC | 12 | C-1-C | 25 | C | 9 | C | ✓ | G |

| | | | | | | | | | | | | | |
|---|---|---|---|---|---|---|---|---|---|---|---|---|---|
| CYCAS_024353 | C | 9 | C | 16 | CC | 12 | C-1-C | 24 | C | 9 | C | - | G/D |
| CYCAS_024354 | C | 9 | C | 18 | CC | 12 | C-1-C | 24 | C | 9 | C | ✓ | G |
| CYCAS_024355 | C | 9 | C | 19 | CC | 12 | C-1-C | 24 | C | 9 | C | - | G/D |
| CYCAS_024356 | C | 9 | C | 17 | CC | 12 | C-1-C | 24 | C | 9 | C | ✓ | G |
| MA_10069449g0010 | C | 9 | C | 14 | CC | 12 | C-1-C | 26 | C | 8 | C | - | G/D |
| MA_10432091g0010 | C | 9 | C | 15 | CC | 12 | C-1-C | 25 | C | 9 | C | - | G/D |
| MA_113140g0010 | C | 9 | C | 19 | CC | 12 | C-1-C | 23 | C | 9 | C | - | G/D |
| MA_138772g0010 | C | 9 | C | 16 | CC | 12 | C-1-C | 25 | C | 9 | C | - | G/D |
| MA_169045g0010 | C | 9 | C | 16 | CC | 12 | C-1-C | 34 | C | 9 | C | ✓ | G |
| MA_494876g0010 | C | 9 | C | 14 | CC | 12 | C-1-C | 24 | C | 9 | C | - | G/D |
| MA_50191g0010 | C | 9 | C | 15 | CC | 12 | C-1-C | 24 | C | 9 | C | ✓ | G |
| MA_697637g0010 | C | 9 | C | 17 | CC | 12 | C-1-C | 24 | C | 8 | C | - | G/D |
| MA_71069g0010 | C | 9 | C | 14 | CC | 12 | C-1-C | 24 | C | 9 | C | ✓ | G |
| MA_76307g0020 | C | 9 | C | 14 | CC | 12 | C-1-C | 24 | C | 9 | C | ✓ | G |
| MA_8892965g0010 | C | 9 | C | 16 | CC | 12 | C-1-C | 24 | C | 9 | C | - | G/D |
| Itaiw_v1_scaffold_52_t29397-RA | C | 9 | C | 14 | CC | 12 | C-1-C | 26 | C | 8 | C | ✓ | G |
| Sacu_v1.1_s0056.g014600 | C | 9 | C | 14 | CC | 12 | C-1-C | 28 | C | 8 | C | ✓ | G |
| Sacu_v1.1_s0073.g017064 / 1$^{st}$ domain | C | 9 | C | 15 | CC | 12 | C-1-C | 26 | C | 9 | C | ✓ | G |
| Sacu_v1.1_s0073.g017064 / 2$^{nd}$ domain | C | 9 | C | 15 | CC | 12 | C-1-C | 26 | C | 9 | C | ✓ | G |
| Smo445118 | C | 10 | C | 13 | CC | 9 | C-1-C | 22 | C | 7 | C | - | G/D |

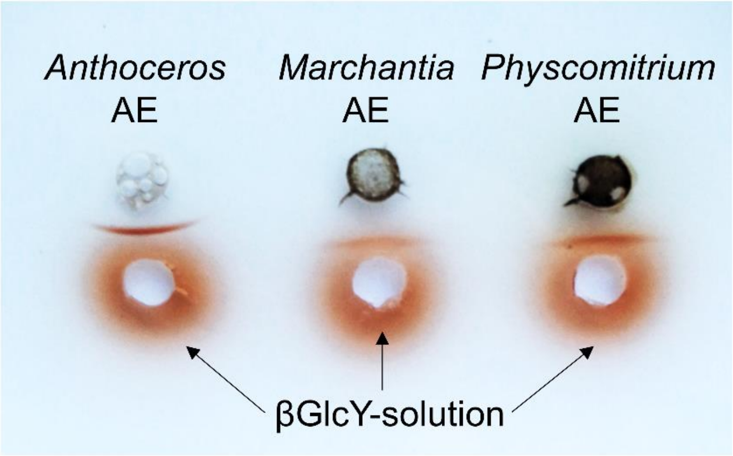

**Figure S1.** Gel diffusion assay of aqueous extracts from *Anthoceros agrestis*, *Marchantia polymorpha* and *Physcomitrium patens* (100 mg $mL^{-1}$) with βGlcY (1 mg $mL^{-1}$). The red precipitation line indicates presence of AGPs.

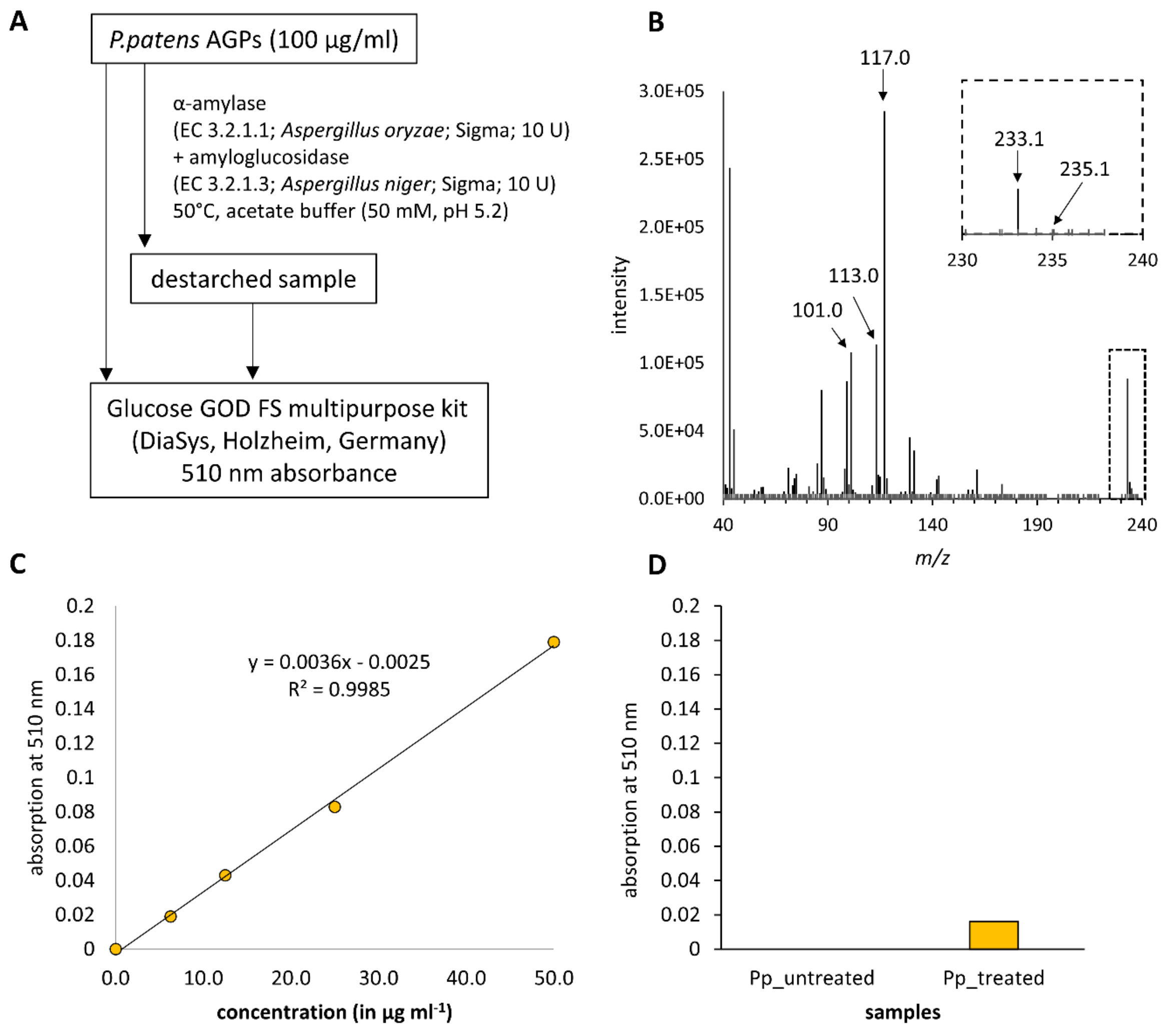

**Figure S2.** Starch analysis in *Physcomitrium patens* AGP. **A**: Workflow for enzymatic digestion and quantification of released glucose by a colorimetric glucose oxidase kit. **B**: Fragmentation pattern of partially methylated and acetylated 1,4-linked glucose in the mass spectrum of the uronic acid reduced sample of *P. patens*. The box highlights the region in which the two diagnostic primary ions m/z = 233 (1,4-linked Glc*p*) and m/z = 235 (1,4-linked Glc*p*A) are found. The extreme overrepresentation of m/z = 233.1 supports the presence of starch. **C**: calibration line for glucose determined with the glucose oxidase kit. **D**: Absorption of *P. patens* AGP before (Pp_untreated) and after (Pp_treated) digestion with α-amylase and amyloglucosidase. The calculated value for released glucose corresponds to approximately 11.2 % (w/w) in the AGP solution.

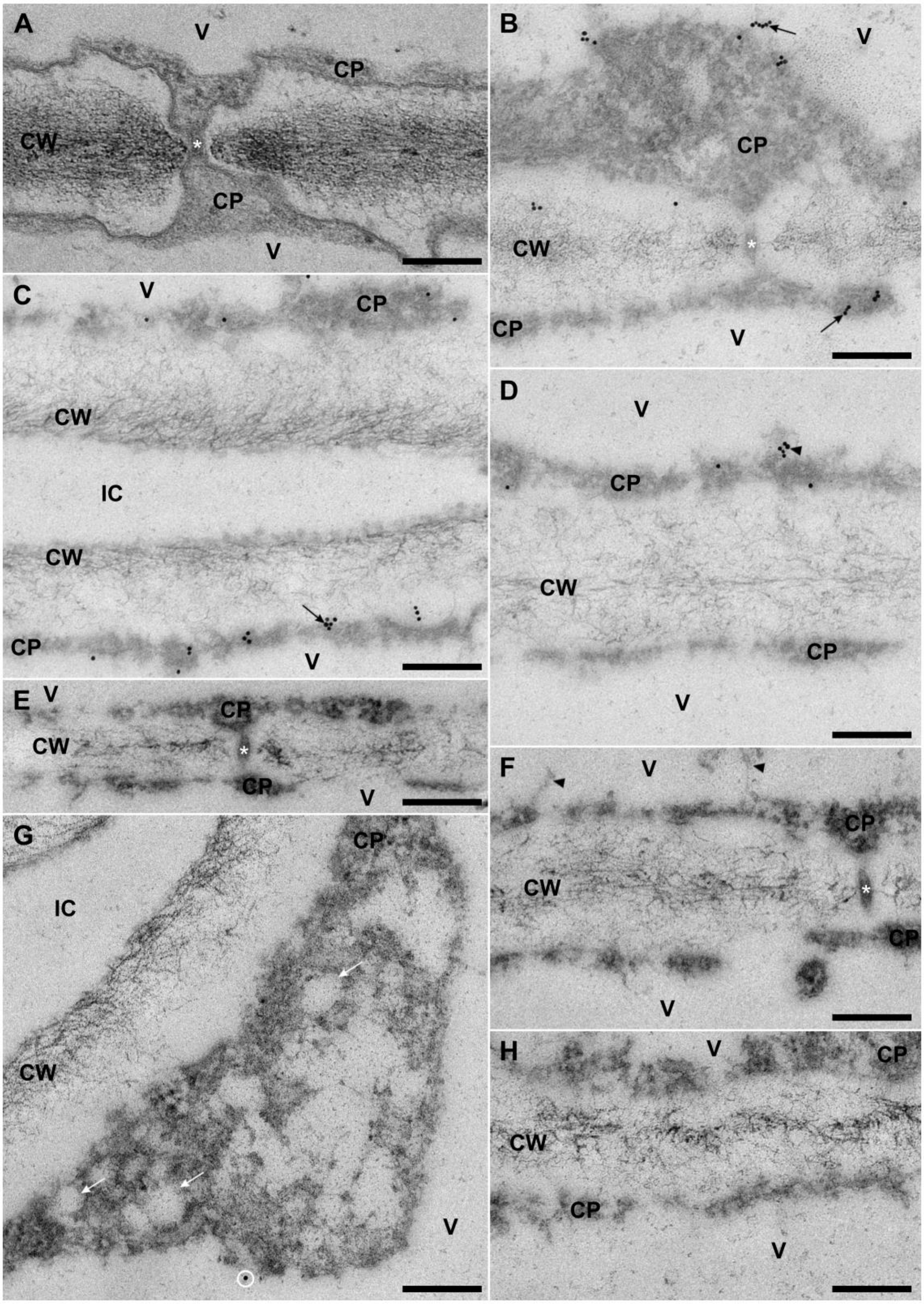

**Figure S3.** Immunolabeling of AGPs detected with JIM13 primary antibodies and 10 nm gold-conjugated secondary antibodies in epidermal cells of *Anthoceros agrestis* gametophyte thalli observed by transmission electron microscopy (TEM). **A:** TEM micrograph of a plasmodesmos (asterisk) in a sample conventionally fixed with glutaraldehyde and osmium tetroxide and embedded in Spurr's Resin. **B-H**: Detail images of samples fixed for immunocytochemistry. **B-D**: With full AGP immunolabeling including the primary antibody. **E-H**: Corresponding control sections treated with the secondary antibody, only. **B**: Plasmodesmata (asterisk) and their directly adjacent cell wall-areas were never labeled, although (clusters of) gold particles occurred at the youngest cell wall-layers, at the plasma membranes and the tonoplasts (black arrows). **C**: Two cells adjacent to a narrow intercellular space showed characteristic labeling of the plasma membrane (e.g. black arrow) and the tonoplast. **D**: Intercellular wall with gold labels at the plasma membrane and the tonoplast, as well as at the dark proteinaceous material in the vacuole (arrowhead) of the adjacent cells. **E**, **F**: Plasmodesmata (asterisks) and the surrounding cell wall material showed no non-specific immunolabels in the control sections. **E-H**: Gold labels also lack on the other regions of the cell walls, at the plasma membranes and tonoplasts, at the proteinaceous material of the vacuole (arrowheads in **F**), as well as at vesicles in the cytoplasm (white arrows in **G**), except for some rare, randomly distributed background labels without a specific association (white circle in **G**). Cell walls adjacent to an intercellular space are shown in **G**, while **E, F** and **H** show intercellular contact walls. Scale bars: 200 nm, CW: cell wall, CP: cytoplasm, IC: intercellular space, V: vacuole. Differences in contrast between samples and controls result from the auto-contrast function of the camera which refers to the black gold particles to adjust the contrast.

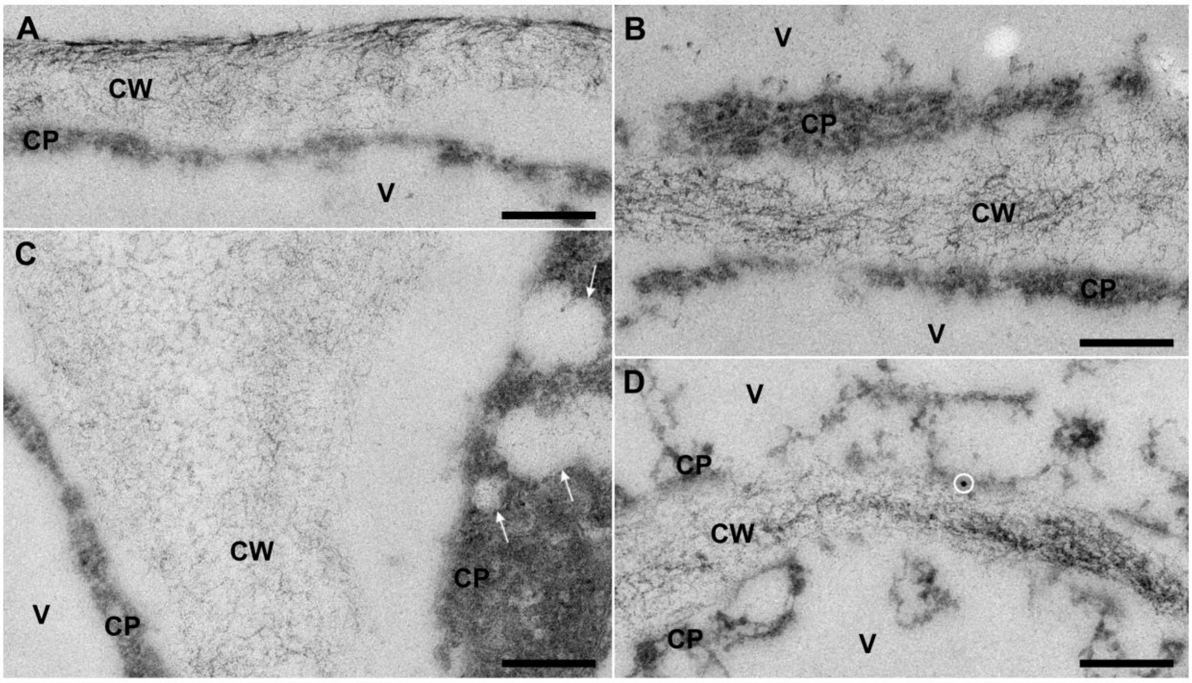

**Figure S4.** Negative controls corresponding to the immunolabeling of AGPs detected with JIM13 primary antibodies and 10 nm gold-conjugated secondary antibodies in epidermal cells of *Anthoceros agrestis* gametophyte thalli observed by transmission electron microscopy (TEM). TEM immunocytochemistry images are comparable to those in Fig. 4, but lack JIM13 primary antibody treatment. (**A**) Externally oriented epidermis wall and (**B**) intercellular contact wall without any non-specific immunolabels at the cell walls, at the cytoplasm, and at the vacuoles. (**C**) No gold particles occurred at the wedge-shaped cell wall-area that borders on an intercellular space. Multiple vesicles (white arrows) in the right cell without label. (**D**) Even the young cell walls only very rarely showed non-specific labeling (white circle). Scale bars: 200 nm, CW: cell wall, CP: cytoplasm, V: vacuole

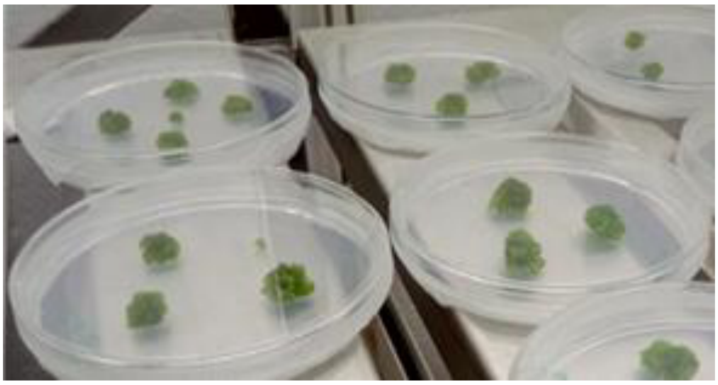

**Figure S5.** Culture of *Anthoceros agrestis* on agar in the Pharmaceutical Institute of Kiel University, Germany.

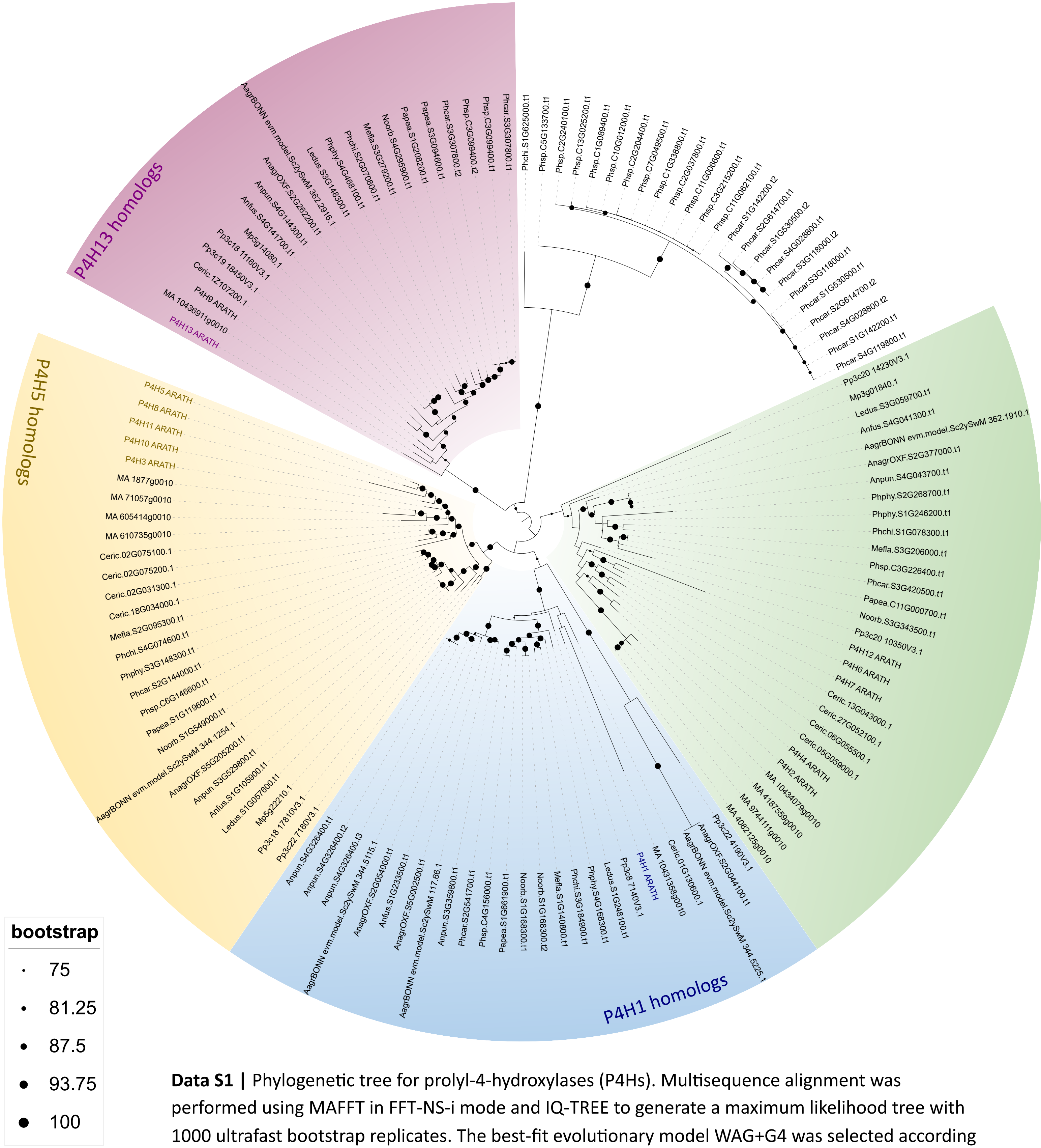

**Data S1 |** Phylogenetic tree for prolyl-4-hydroxylases (P4Hs). Multisequence alignment was performed using MAFFT in FFT-NS-i mode and IQ-TREE to generate a maximum likelihood tree with 1000 ultrafast bootstrap replicates. The best-fit evolutionary model WAG+G4 was selected according to Bayesian Information Criterion.

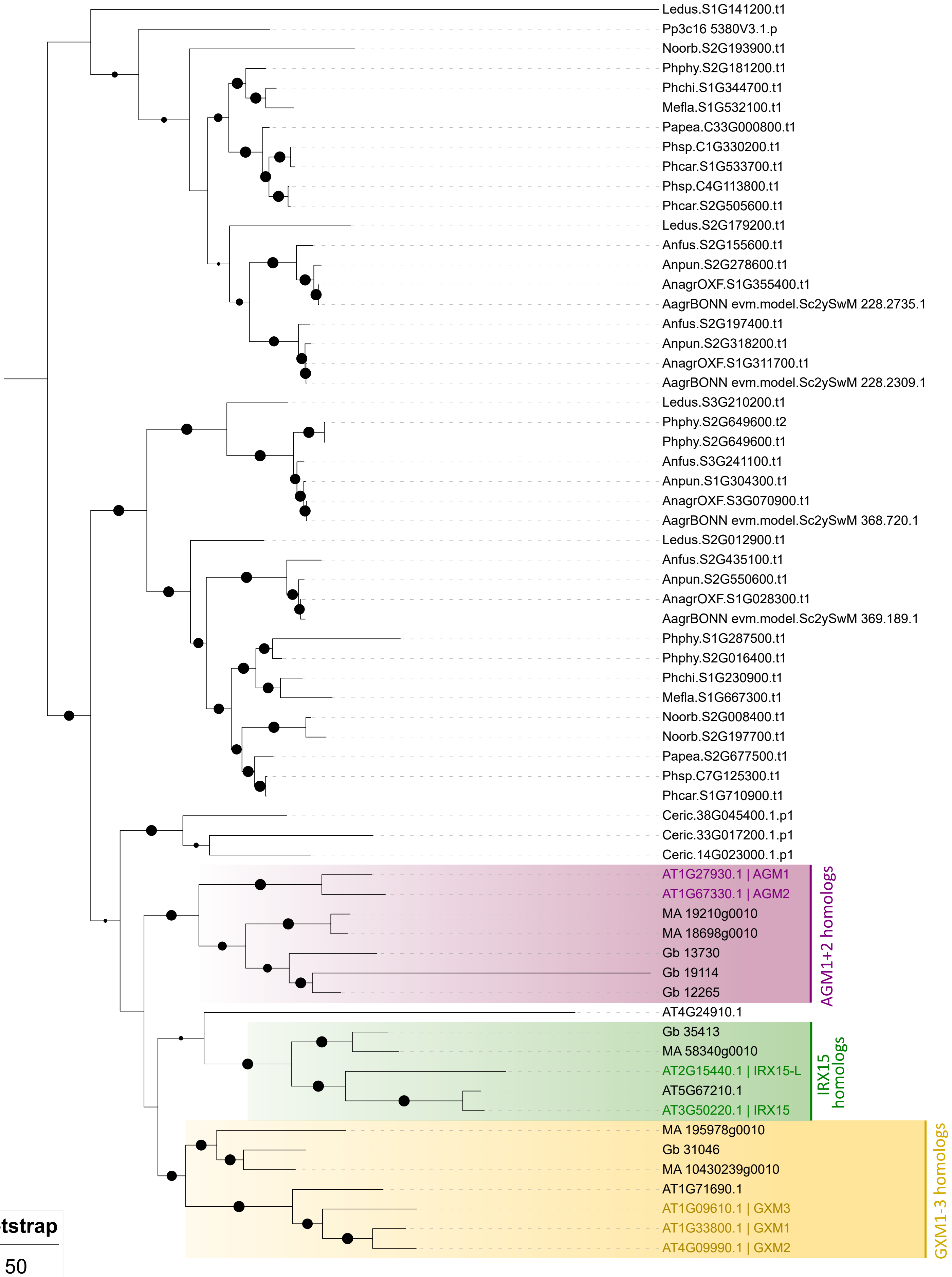

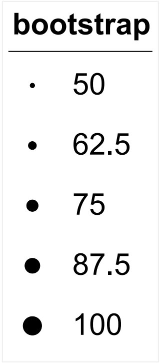

**Data S2 |** Phylogenetic tree for DUF579 homologs. Multisequence alignment was performed using MAFFT in L-INS-i mode and IQ-TREE to generate a maximum likelihood tree with 1000 ultrafast bootstrap replicates. The best-fit evolutionary model WAG+G4 was selected according to Bayesian Information Criterion.

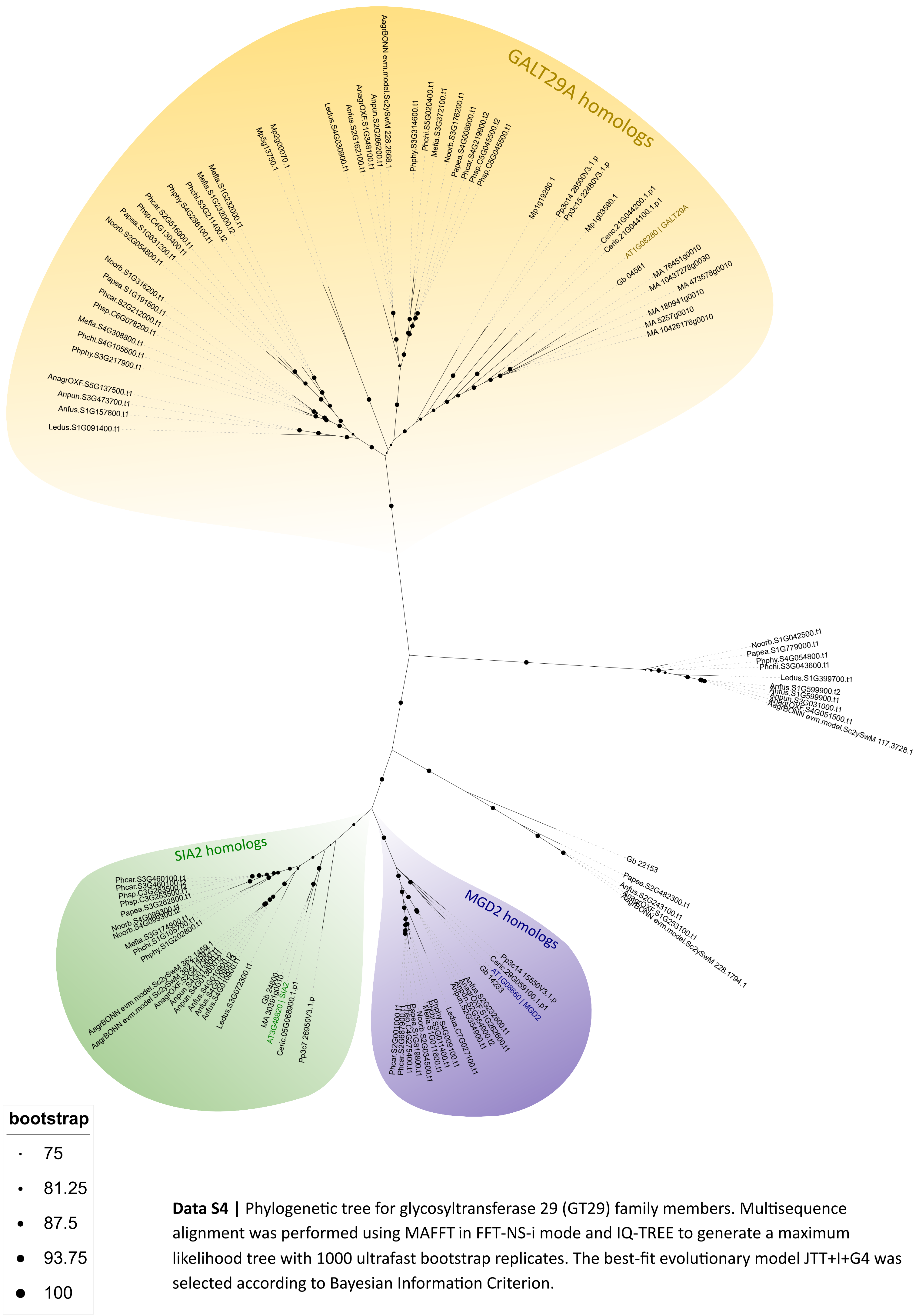

**Data S4 | Phylogenetic tree for glycosyltransferase 29 (GT29) family members.** Multisequence alignment was performed using MAFFT in FFT-NS-i mode and IQ-TREE to generate a maximum likelihood tree with 1000 ultrafast bootstrap replicates. The best-fit evolutionary model JTT+I+G4 was selected according to Bayesian Information Criterion.

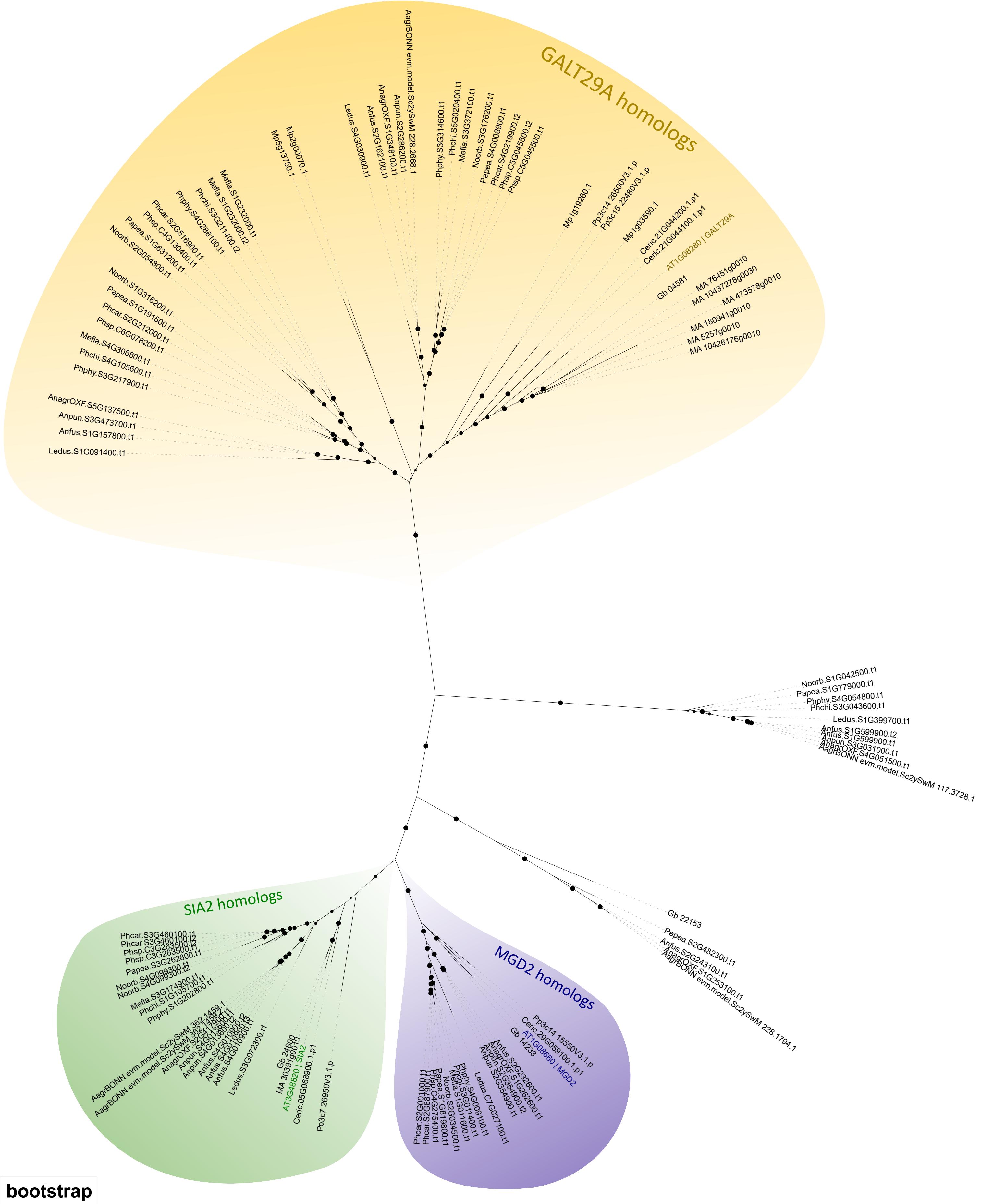

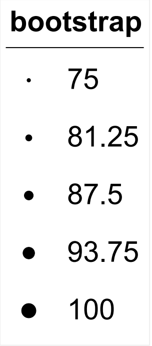

**Data S4 |** Phylogenetic tree for glycosyltransferase 29 (GT29) family members. Multisequence alignment was performed using MAFFT in FFT-NS-i mode and IQ-TREE to generate a maximum likelihood tree with 1000 ultrafast bootstrap replicates. The best-fit evolutionary model JTT+I+G4 was selected according to Bayesian Information Criterion.

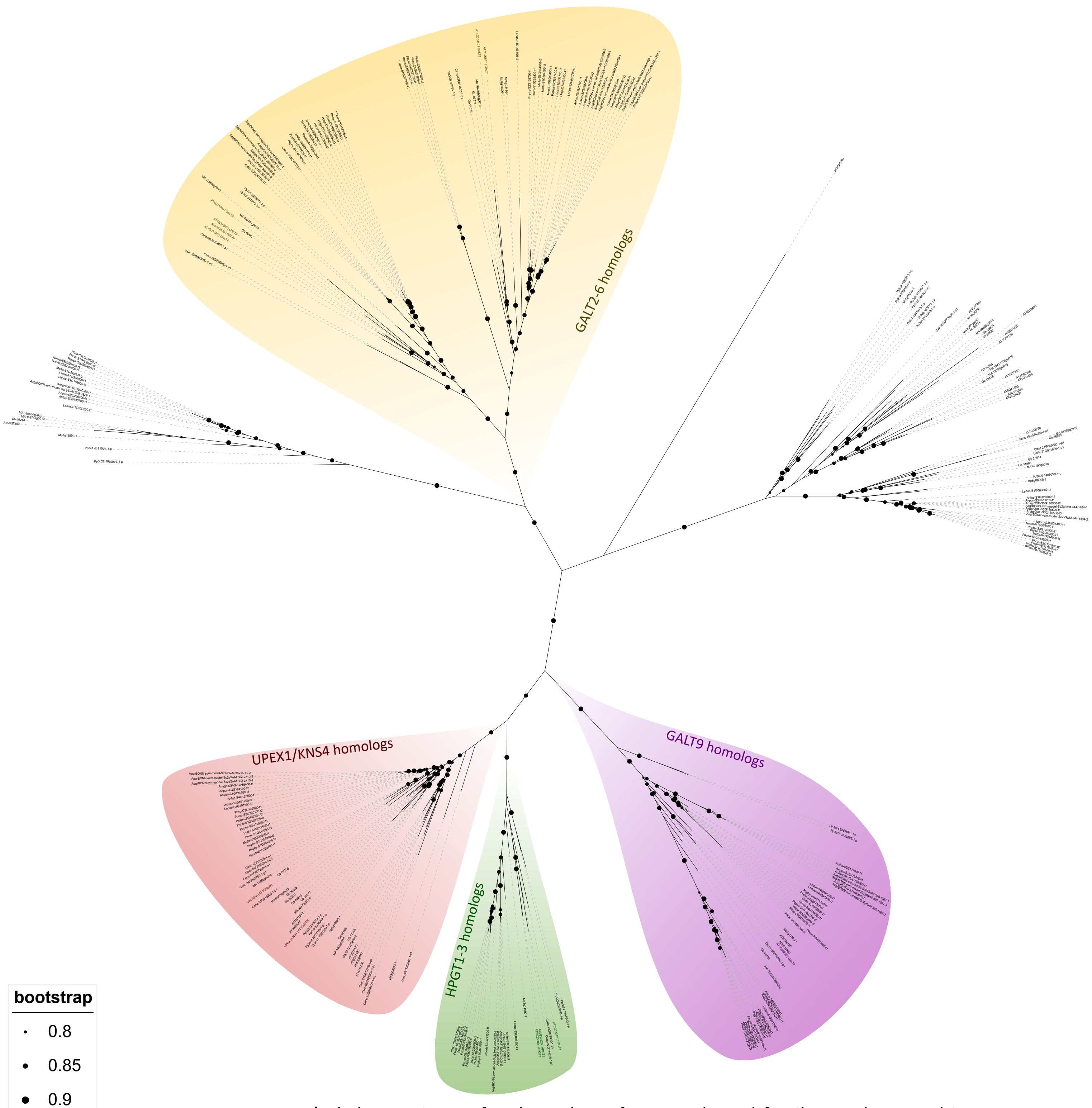

**Data S5 |** Phylogenetic tree for glycosyltransferase 31 (GT31) family members. Multisequence alignment was performed using MAFFT in FFT-NS-i mode and FastTree 2 to generate an approximately maximum likelihood tree. The evolutionary model JTT-CAT was chosen.

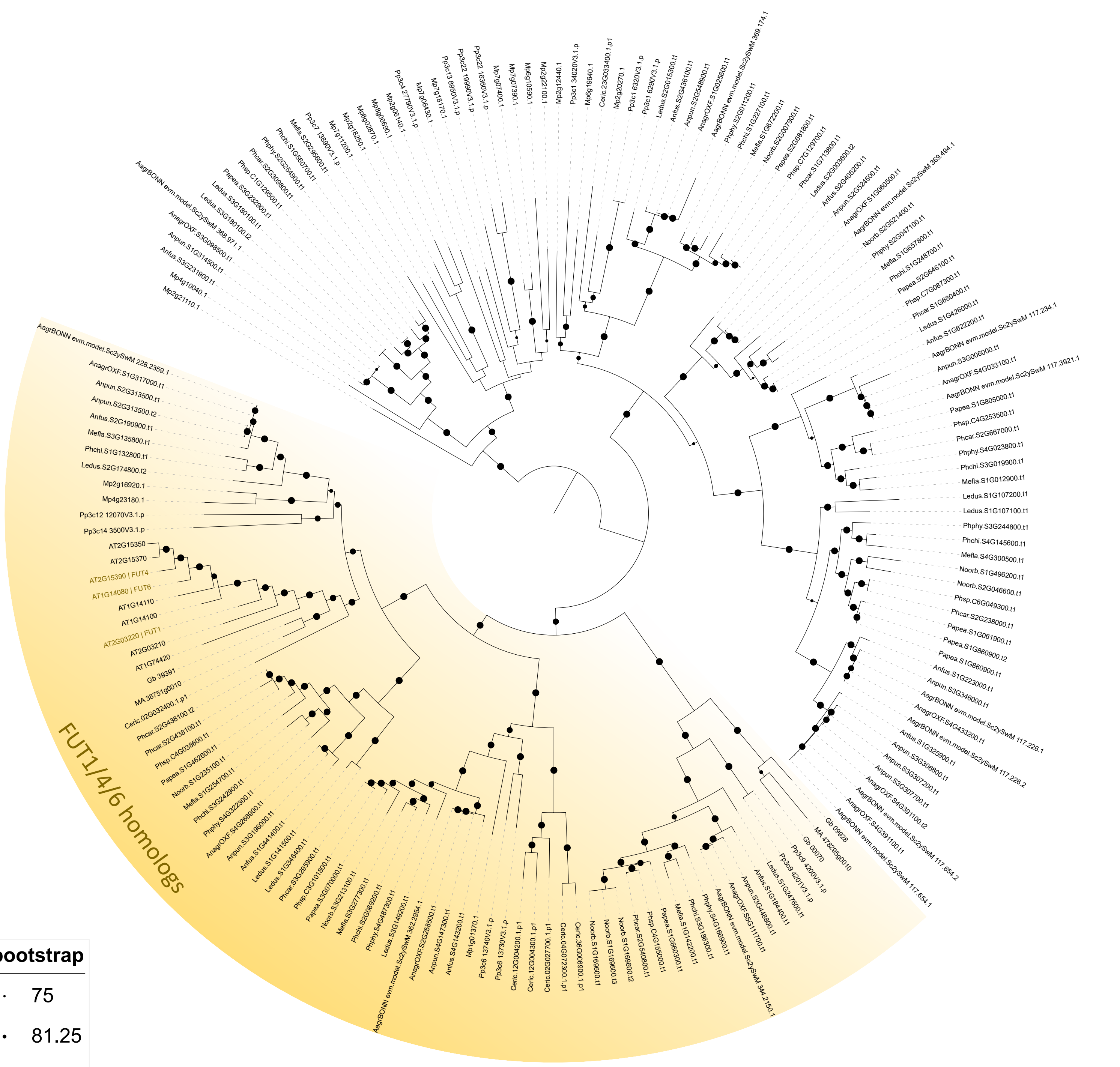

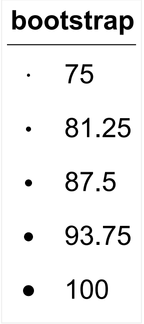

**Data S6 |** Phylogenetic tree for glycosyltransferase 37 (GT37) family members. Multisequence alignment was performed using MAFFT in FFT-NS-i mode and IQ-TREE to generate a maximum likelihood tree with 1000 ultrafast bootstrap replicates. The best-fit evolutionary model WAG+F+I+G4 was selected according to Bayesian Information Criterion.

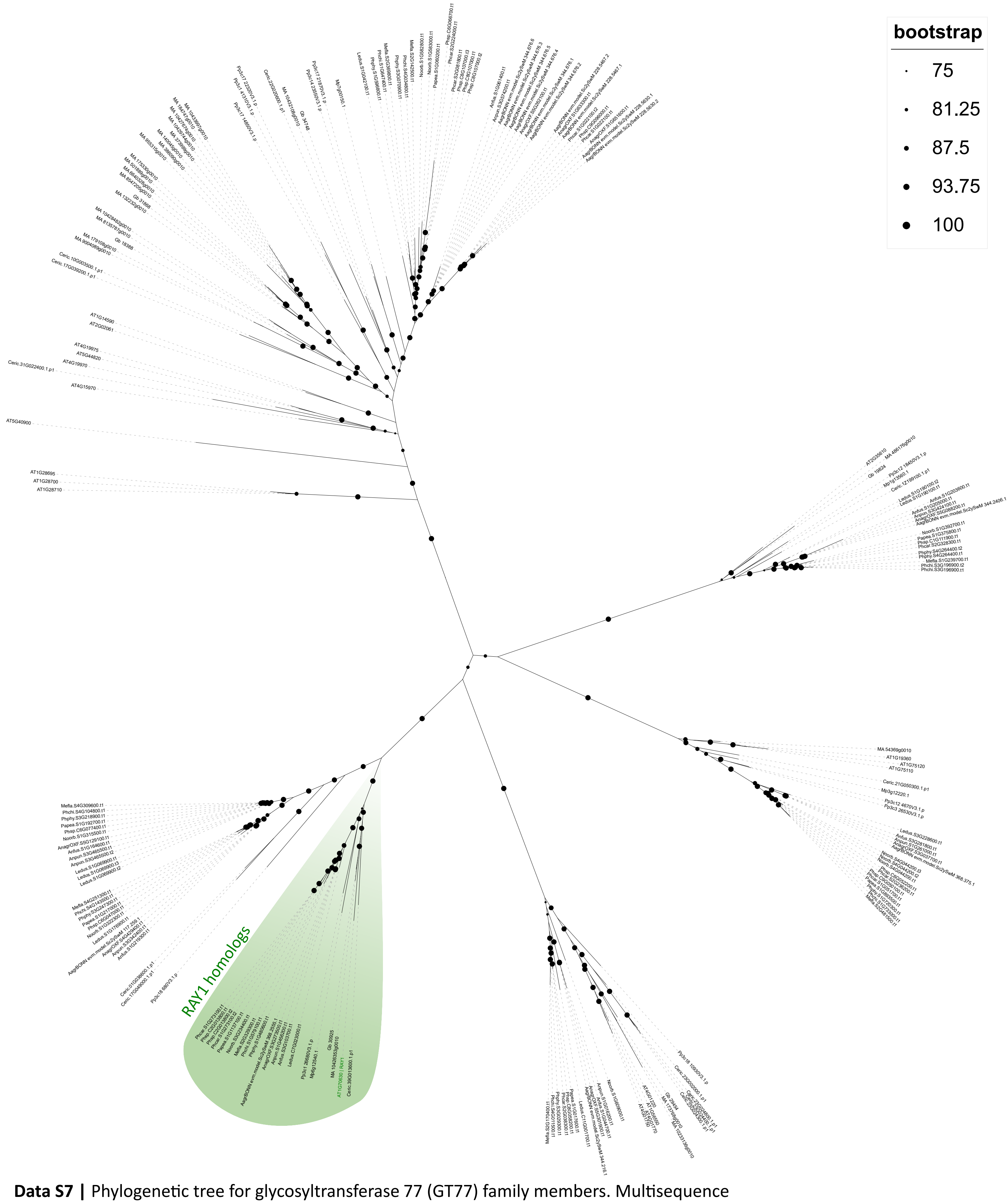

**Data S7 |** Phylogenetic tree for glycosyltransferase 77 (GT77) family members. Multisequence alignment was performed using MAFFT in L-INS-i mode and IQ-TREE to generate a maximum likelihood tree with 1000 ultrafast bootstrap replicates. The best-fit evolutionary model WAG+I+G4 was selected according to Bayesian Information Criterion.

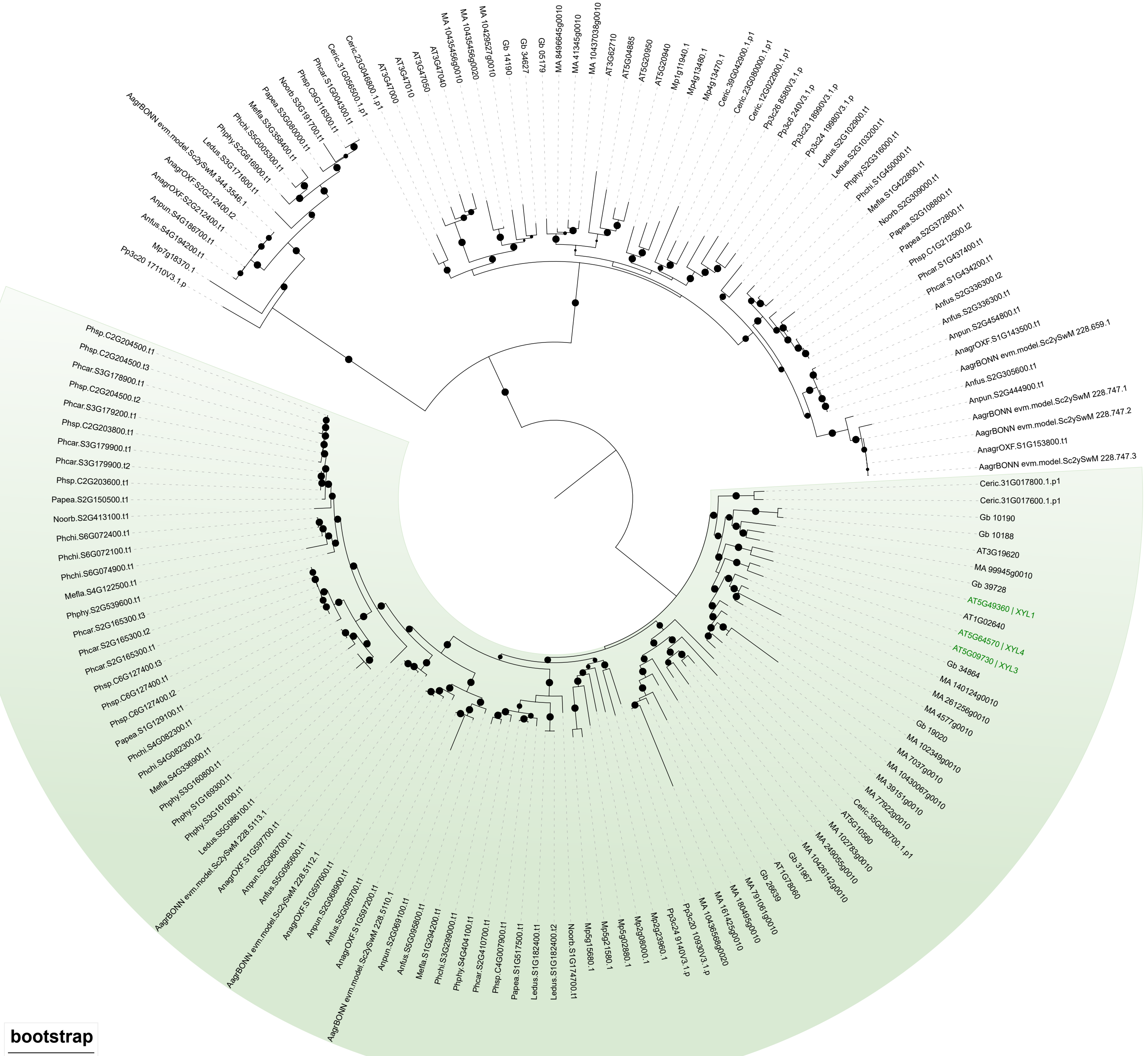

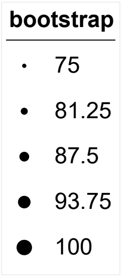

**Data S8 |** Phylogenetic tree for glycosylhydrolase 3 (GH3) family members. Multisequence alignment was performed using MAFFT in FFT-NS-i mode and IQ-TREE to generate a maximum likelihood tree with 1000 ultrafast bootstrap replicates. The best-fit evolutionary model WAG+I+G4 was selected according to Bayesian Information Criterion.

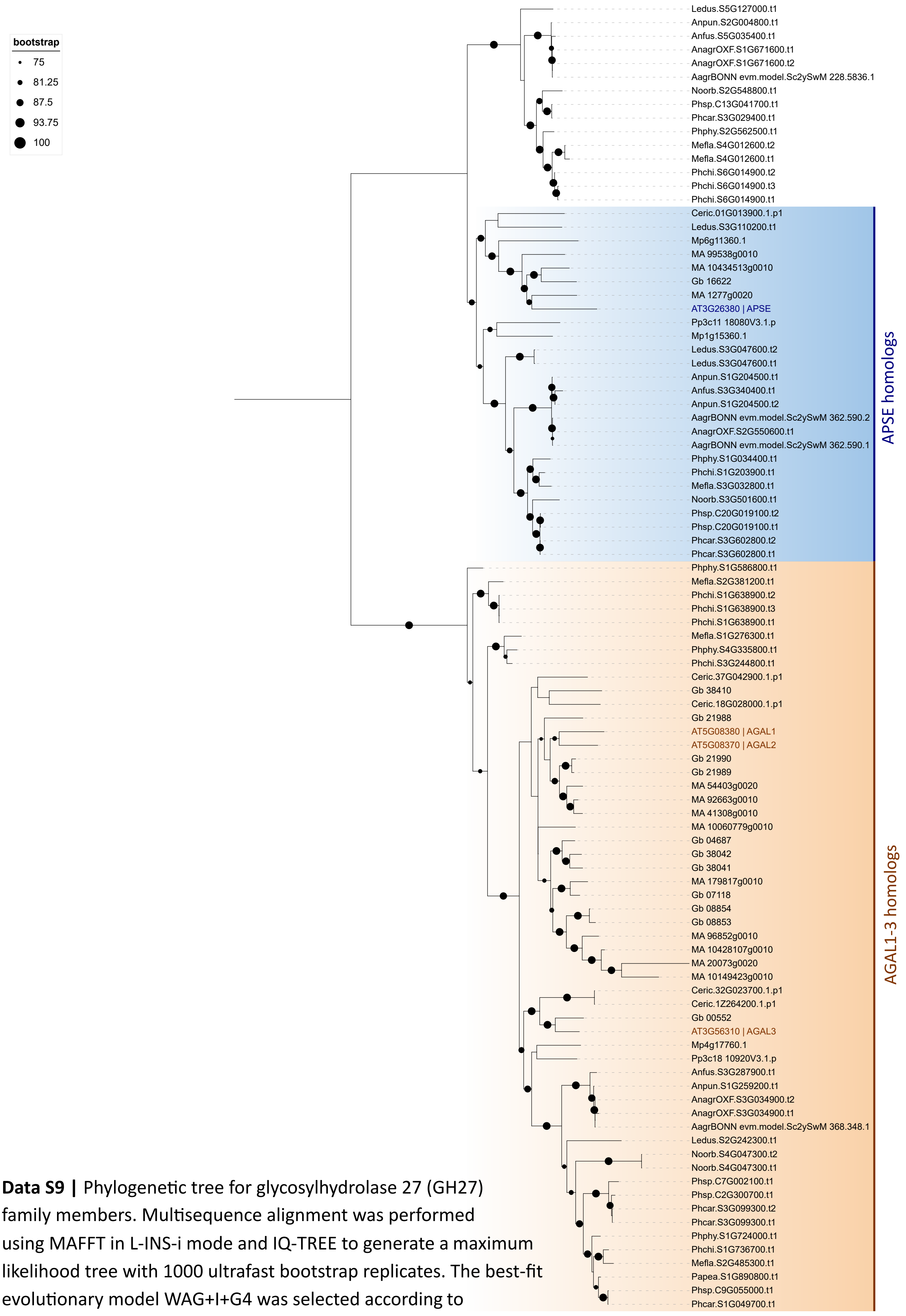

**Data S9 |** Phylogenetic tree for glycosylhydrolase 27 (GH27) family members. Multisequence alignment was performed using MAFFT in L-INS-i mode and IQ-TREE to generate a maximum likelihood tree with 1000 ultrafast bootstrap replicates. The best-fit evolutionary model WAG+I+G4 was selected according to Bayesian Information Criterion.

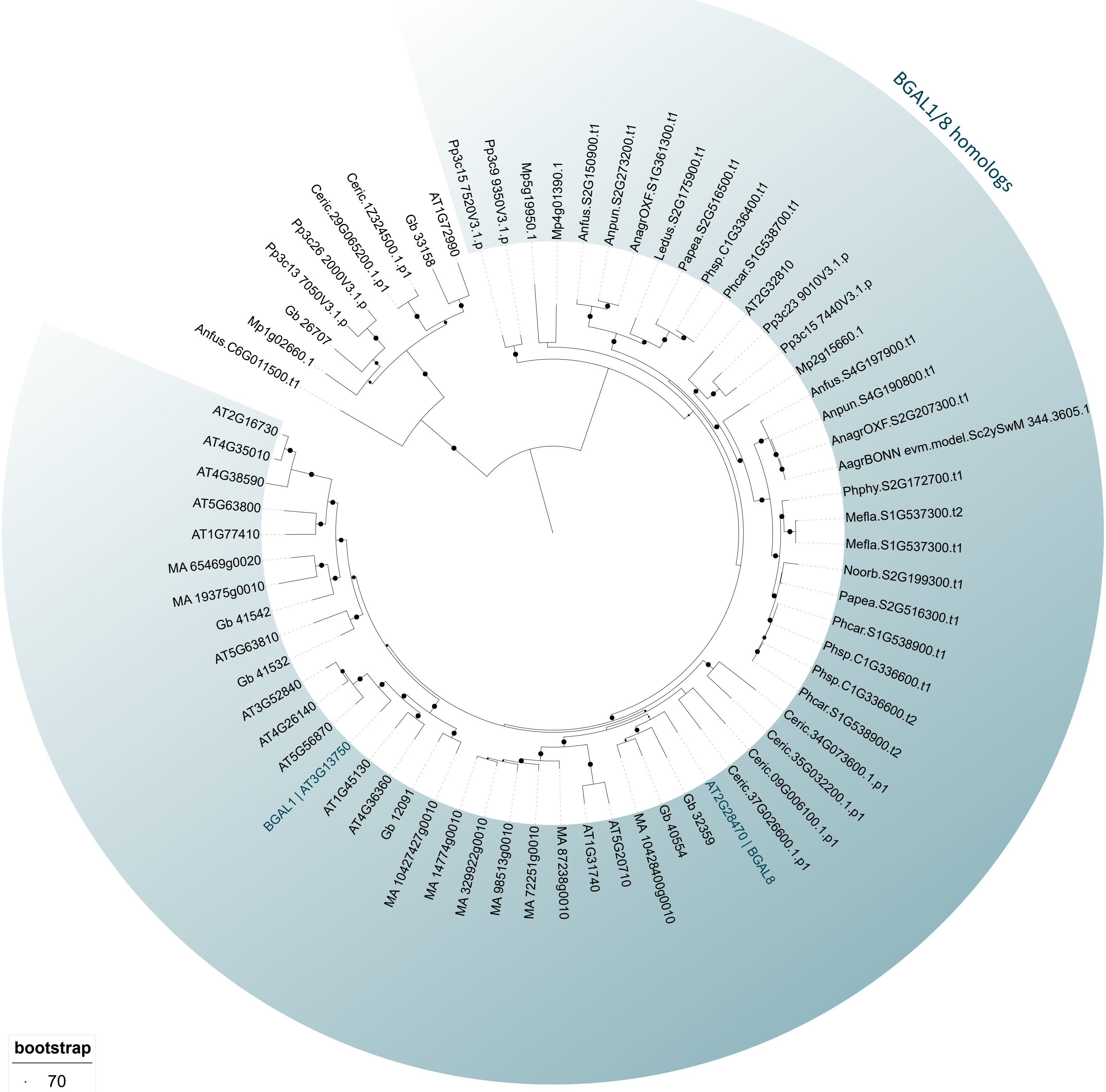

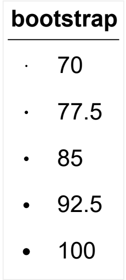

**Data S 10 | Phylogenetic tree for glycosylhydrolase 35 (GH35) family members.** Multisequence alignment was performed using MAFFT in L-INS-i mode and IQ-TREE to generate a maximum likelihood tree with 1000 ultrafast bootstrap replicates. The best-fit evolutionary model WAG+I+G4 was selected according to Bayesian Information Criterion.

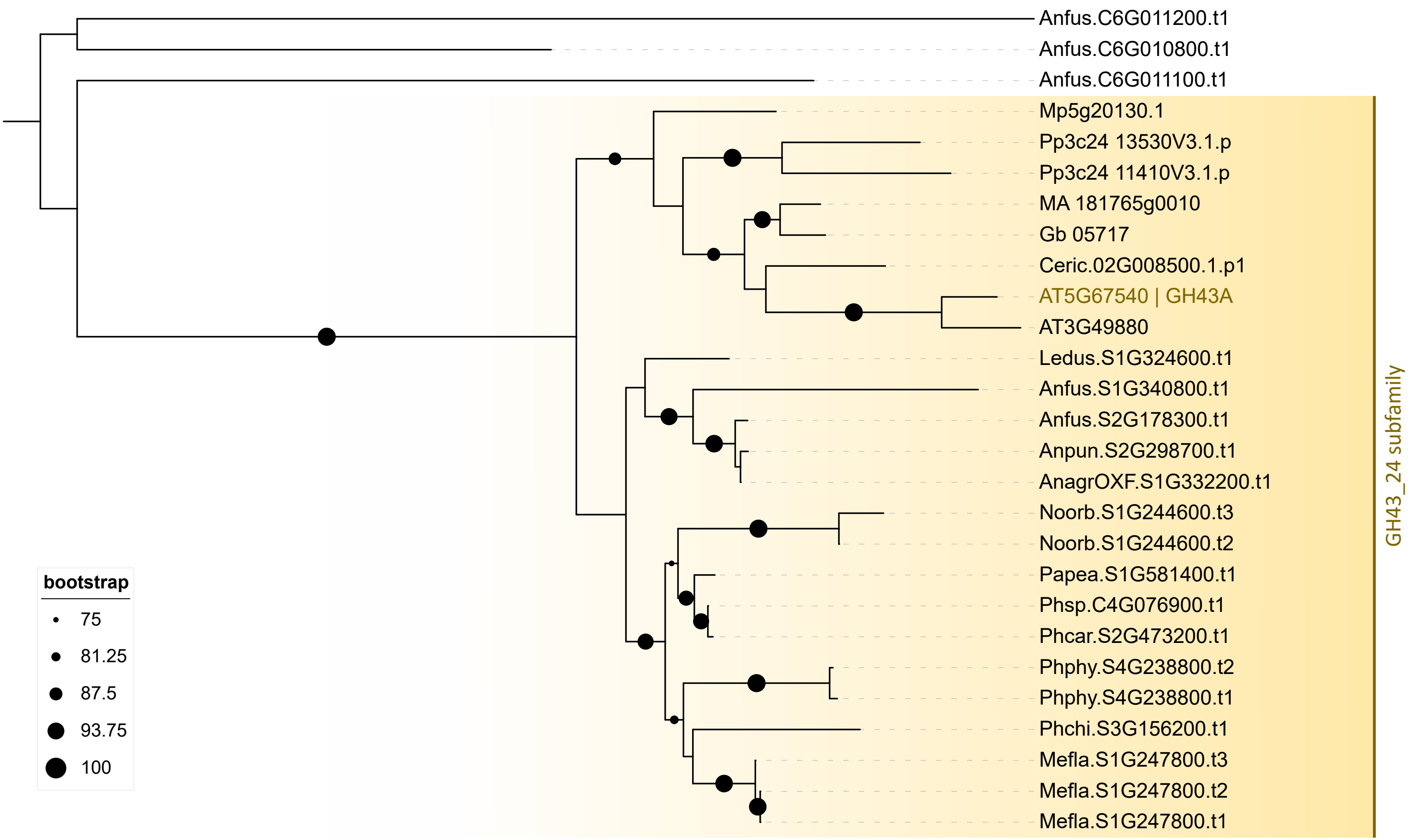

**Data S 11 |** Phylogenetic tree for glycosylhydrolase 43 (GH43) family members. Multisequence alignment was performed using MAFFT in L-INS-i mode and IQ-TREE to generate a maximum likelihood tree with 1000 ultrafast bootstrap replicates. The best-fit evolutionary model WAG+G4 was selected according to Bayesian Information Criterion.

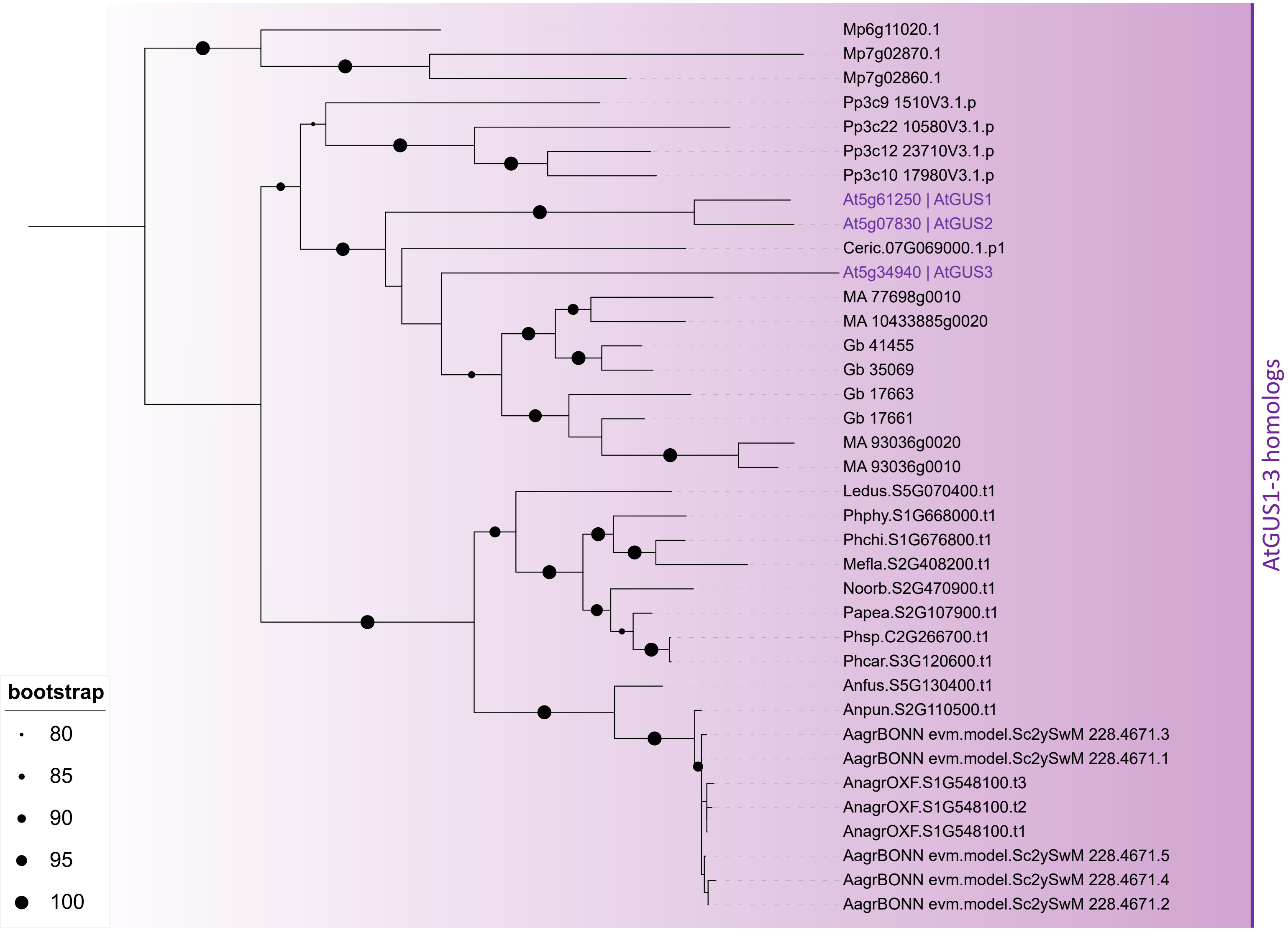

**Data S 12 |** Phylogenetic tree for glycosylhydrolase 79 (GH79) family members. Multisequence alignment was performed using MAFFT in L-INS-i mode and IQ-TREE to generate a maximum likelihood tree with 1000 ultrafast bootstrap replicates. The best-fit evolutionary model WAG+I+G4 was selected according to Bayesian Information Criterion.
